# North Pontic crossroads: Mobility in Ukraine from the Bronze Age to the early modern period

**DOI:** 10.1101/2024.05.24.595769

**Authors:** Lehti Saag, Olga Utevska, Stanislav Zadnikov, Iryna Shramko, Kyrylo Gorbenko, Mykola Bandrivskyi, Dmytro Pavliv, Igor Bruyako, Denys Grechko, Vitalii Okatenko, Gennadi Toshev, Svitlana Andrukh, Vira Radziyevska, Yurii Buynov, Viktoriia Kotenko, Oleksandr Smyrnov, Oleg Petrauskas, Borys Magomedov, Serhii Didenko, Anatolii Heiko, Roman Reida, Serhii Sapiehin, Viktor Aksonov, Oleksii Laptiev, Svyatoslav Terskyi, Viacheslav Skorokhod, Vitalii Zhyhola, Yurii Sytyi, Mari Järve, Christiana Lyn Scheib, Kyriaki Anastasiadou, Monica Kelly, Mia Williams, Marina Silva, Christopher Barrington, Alexandre Gilardet, Ruairidh Macleod, Pontus Skoglund, Mark G. Thomas

**Author notes:** Corresponding authors Lehti Saag, Pontus Skoglund, Mark G. Thomas. Estonian Biocentre, Institute of Genomics, University of Tartu, Tartu 51010, Estonia (Lehti Saag). Lead Contact: Lehti Saag.

## Abstract

The North Pontic region, which encompasses present-day Ukraine, was a crossroads of migration as it connected the vast Eurasian Steppe with Central Europe. We generated shotgun-sequenced genomic data for 91 individuals dating from around 7,000 BCE to 1,800 CE to study migration and mobility history in the region, with a particular focus on historically attested migrating groups during the Iron Age and the medieval period, such as Scythian, Chernyakhiv, Saltiv and Nogai associated peoples. We infer a high degree of temporal heterogeneity in ancestry, with fluctuating genetic affinities to present-day Western European, Eastern European, Western Steppe and East Asian groups. We also infer high heterogeneity in ancestry within geographically, culturally and socially defined groups. Despite this, we find that ancestry components which are widespread in Eastern and Central Europe have been present in the Ukraine region since the Bronze Age.

## Introduction

Migration has been a major factor shaping human societies, culture, biology and genomes through time. Previous ancient DNA (aDNA) research indicates that, to a first order of approximation, the genomes of present-day Europeans comprise ancestries from three major Holocene groups of people ^1,2^: 1) indigenous hunter-gatherers (HGs); 2) Near Eastern early farmers, arriving in Europe ∼8,000 years ago (ya) ^3^; 3) Steppe pastoralists, who migrated into Europe ∼5,000 ya ^4^. However, the detailed genetic history of any given region is necessarily more complex, calling for more focused and local-scale studies.

One such, and so far relatively understudied region is present-day Ukraine in the Northern Black Sea (Pontic) region, which is historically and archaeologically known as a contact zone between European and Asian populations. Archaeological and genetic data indicate not only ancestries from the above three broad-scale sources, but that it was the first area with early farmer habitation that the steppe pastoralists reached on their westward migration ^4–7^. The subsequent admixture between these two groups is particularly interesting as it was quickly followed by the emergence of the Corded Ware, Sintashta, Andronovo and Zrubna (Srubnaya) cultures over wide areas of central Eurasia ^1,2,8^. Furthermore, due to southern Ukraine forming part of the Eurasian Steppe, which reaches from Hungary in the west all the way to Northeast China in the east, the area has been in the path of extensive if punctuated genetic and cultural flow ^9^.

The accessibility of the Ukrainian Steppe, the presence of an extensive hydrological system, the abundance of raw materials, and the combination of fertile forest-steppe soils with open steppe spaces, large forests and mountain ranges attracted not only various nomadic groups, but also representatives of ancient sedentary civilizations who founded colonies on the Black Sea coast ^9^.

For centuries, migration took place in the steppe and forest-steppe belt of Ukraine, moving in several directions (Figure 1). These migrations were driven by various processes, including periodic aridization, the development of new subsistence strategies and economies, intertribal cultural contacts and conflicts, trade, demographic pressures, and expansion of nomad influence zones. Major migration flows came from the Carpathian-Danavian region, the Southern Urals and Volga region, Central Asia, the North Caucasus, etc., and intensive population movements also occurred within the territory of Ukraine ^10–12^.

**Figure 1.**
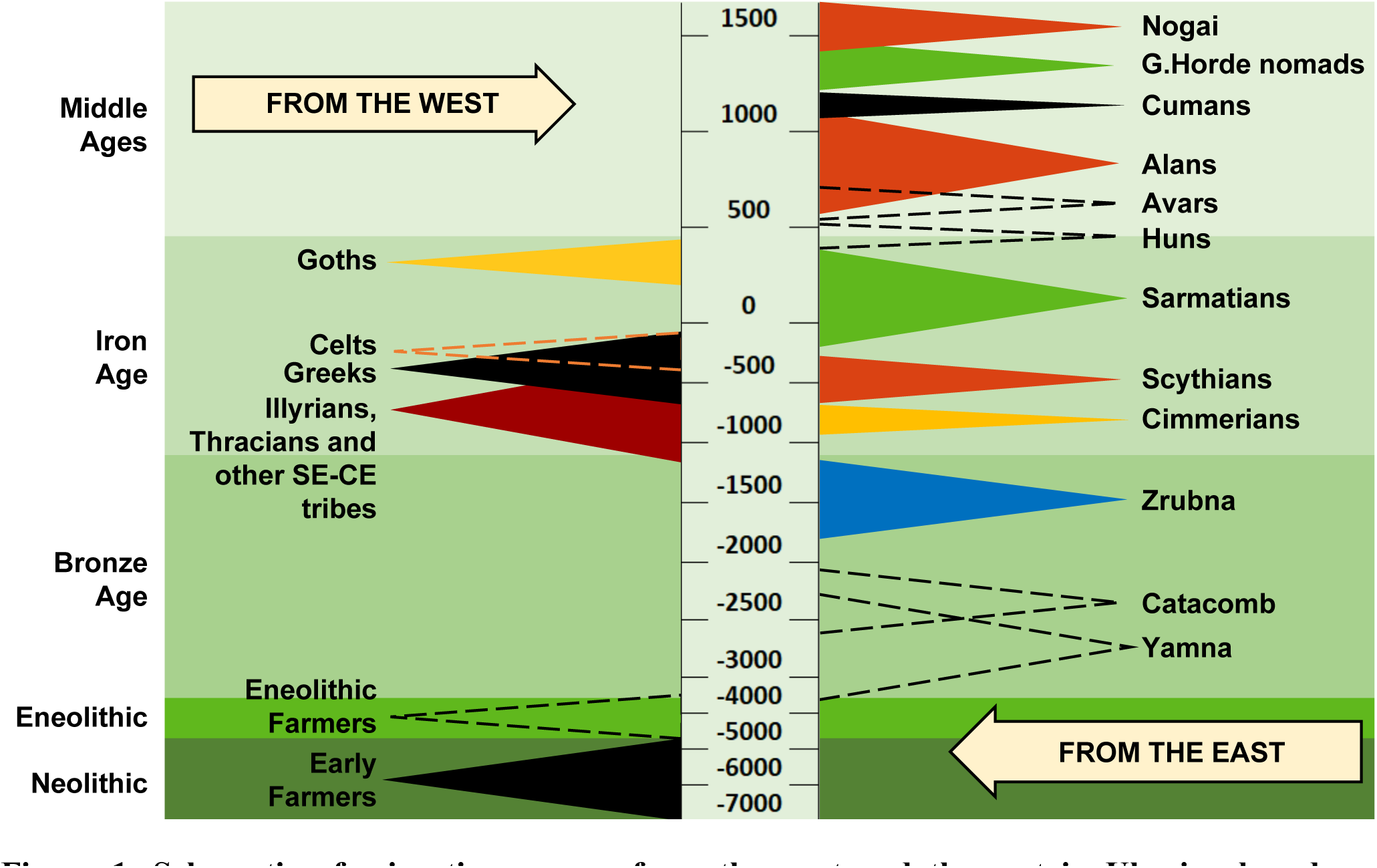
Schematic of migrating groups from the west and the east in Ukraine based on archaeological material. Negative values on the timeline denote years BCE, positive values years CE. See also Table S1.

At the end of the Bronze Age and beginning of the Early Iron Age, the most archaeologically conspicuous activities of the North Pontic steppes were associated with Cimmerians and their military campaigns in Asia Minor ^13^. The Cimmerians were followed by Scythians and Sarmatians, Early Iron Age political and military tribal unions ^9,14^ with variable combinations of local and East Asian ancestry, as indicated by previous ancient DNA studies ^7,15–18^. At this time, the Northern Black Sea coast was covered with a network of urbanized Greek colonies ^19,20^. In the forest-steppe zone, the contemporary settled populations were associated with the previous Tshinets Cultural Circle ^21^ (Lusatian, Vysotska cultures), as well as with Central European migrants of Hallstatt and La Tène periods (Illirians, Thracians and Celts) ^22–25^. According to written and archaeological sources, peoples that are considered the predecessors of the Slavs (associated with the Zarubinetska culture) had already been present in the Ukraine region during the La Tène and Roman periods, from the 3rd century (c.) BCE onwards ^26^.

The beginning of the Migration Period in the Ukraine region is associated with the arrival of Germanic tribes such as Goths, and the formation of the multi-ethnic Chernyakhiv culture, which included other peoples who already inhabited the region ^27,28^. In the 2nd–4th centuries, the Huns – nomadic people from Central Asia – appeared in the North Pontic steppe, and their westward migration led to significant economic, cultural and social changes in Europe. This period is associated with the emergence of a new ethnolinguistic group, the Slavs, who spread throughout much of eastern Europe during 5th–7th c. CE^29,30^.

In the 8th–10th centuries, а significant part of Ukraine was under the control of Khazar Khaganate. In Ukrainian archaeology this is represented by the Saltiv culture, which is thought to have been shared among multiple ethnic groups (Alans, Bulgars, Turks, Slavs, Magyars, etc.). During the same period, there was a process of unification of the Slavic tribes, and in the 9th c. CE the state of Kievan Rus was formed ^31^.

The development of Slavic statehood took place against the background of constant nomadic incursions from the east. In the period from the 11th to the 13th c. CE, waves of Pechenegs, Torques and Cumans entered the North Pontic region from Central Asia, and the most substantial invasion in terms of military strength and consequences was that of the Mongols of the Golden Horde in the 13th c. CE ^32^. By the 15th c. CE, remnants of the Golden Horde population, such as the Nogai, were still living in the North Pontic steppes ^33^. Since the 16th c. CE, Slavs have been the majority ethnolinguistic group in the region of Ukraine ^31,32^.

To date, published ancient genomic data from Ukraine are available mostly for Mesolithic to Bronze Age individuals, some Iron Age Scythians and only a few individuals from other periods ^6,7,16,34–41^. Due to the need to distinguish between groups of individuals and for brevity, we will refer to individuals by the archaeological culture context with which they have been associated. Importantly, it should not be assumed that there is a direct link between culture and genetic ancestry ^42^.

Genomic studies of HGs in Ukraine indicate closer genetic affinities to Eastern, rather than Western European HGs ^6,37,41^. Early farmers in the region, represented by Trypillia and Globular Amphora Culture-associated individuals, show indications of admixture with HGs ^6,36^. Eneolithic Cernavodă I and Usatove Culture-associated individuals display admixture with steppe ancestry groups ^39^. Genomic data from four Yamna culture-associated individuals have been published, with one displaying admixture with European early farmers ^6,7^. There is considerable heterogeneity in the 13 Scythian genomes available, with various combinations of early farmer, HG/steppe and also East Asian ancestry present ^7,16^. The three published Chernyakhiv individuals resemble modern Europeans, but do not form a genetically homogenous group ^7^. These results provide substantial evidence of unresolved genetic structure and population discontinuity in Ukraine, and fragmented relationships between genetic ancestry and material culture associations, starting from the Neolithic, pointing to a need for more paleogenomic data. Furthermore, no data are available from the Middle Ages; such data would extend our understanding of the demographic history of the area.

In this study, we shed light on the demographic history of Ukraine from the Bronze Age to the early modern period. We set out to examine the genetic ancestries of people living in the North Pontic region during different time periods and associated with various cultural groups. We do not encompass the entirety of local cultures, but focus on those introduced by migrants, which include a wide range of nomadic warrior groups (Figure 1). We investigate if individuals associated with specific cultural groups are genetically homogenous or if genetic structure exists within them. More specifically, we assess the extent to which admixture occurs between local and migrant groups, and to what extent autochthonous peoples are culturally assimilated into migrant groups. For Scythian-associated individuals, we explore genetic differences between individuals with nomad *vs* local and elite *vs* non-elite archaeological assignments as well as between different areas of Ukraine.

## Results

### Samples and archaeological background

We extracted DNA from the apical tooth roots and bone fragments of 128 individuals from 33 archaeological sites in present-day Ukraine (Table S1, Methods). The 91 individuals that were chosen for further sequencing and analyses (Figure 2, Table 1) yielded on average 49% endog-enous DNA (Table S1). We shotgun-sequenced these individuals to an average genomic coverage of >0.01× (n = 13), >0.1× (n = 9), >0.3× (n = 34), >0.5× (n = 27), and >1× (n = 8) (Table 1, Table S1). The presented genome--wide data are derived from 1 Neolithic individual (UkrN; 7,000–6,000 BCE), 9 individuals from the Bronze Age and from the Final Bronze Age to the beginning of the Iron Age (UkrBA, UkrFBA/EIA; 3,000–700 BCE), 6 individuals from the beginning of the Early Iron Age (UkrEIA; 900– 700 BCE), 29 individuals from the Scythian period of the Early Iron Age (UkrEIA; 700–300 BCE), 6 individuals from the end of the Early Iron Age (UkrEIA; 400–1 BCE), 12 individuals from the later Iron Age (UkrIA; 1–400 CE), 9 individuals from the Early Middle Ages (UkrEMA; 800–900 CE), and 19 individuals from the Middle Ages to the early modern period (UkrMA, UkrEM; 900–1,800 CE) (Figure 2, Table S1, Methods). We analysed the data in the context of published ancient and modern genomes, which have been assigned to groups based on ancestry, geography, time period and archaeological culture, as defined in the Allen Ancient DNA Resource ^43^. Throughout this study we term these as AADR groups (Tables S3–S4). Previously published genomes from individuals with the same archaeological associations ^7^ were analysed and discussed alongside individuals sequenced in this study.

**Figure 2.**
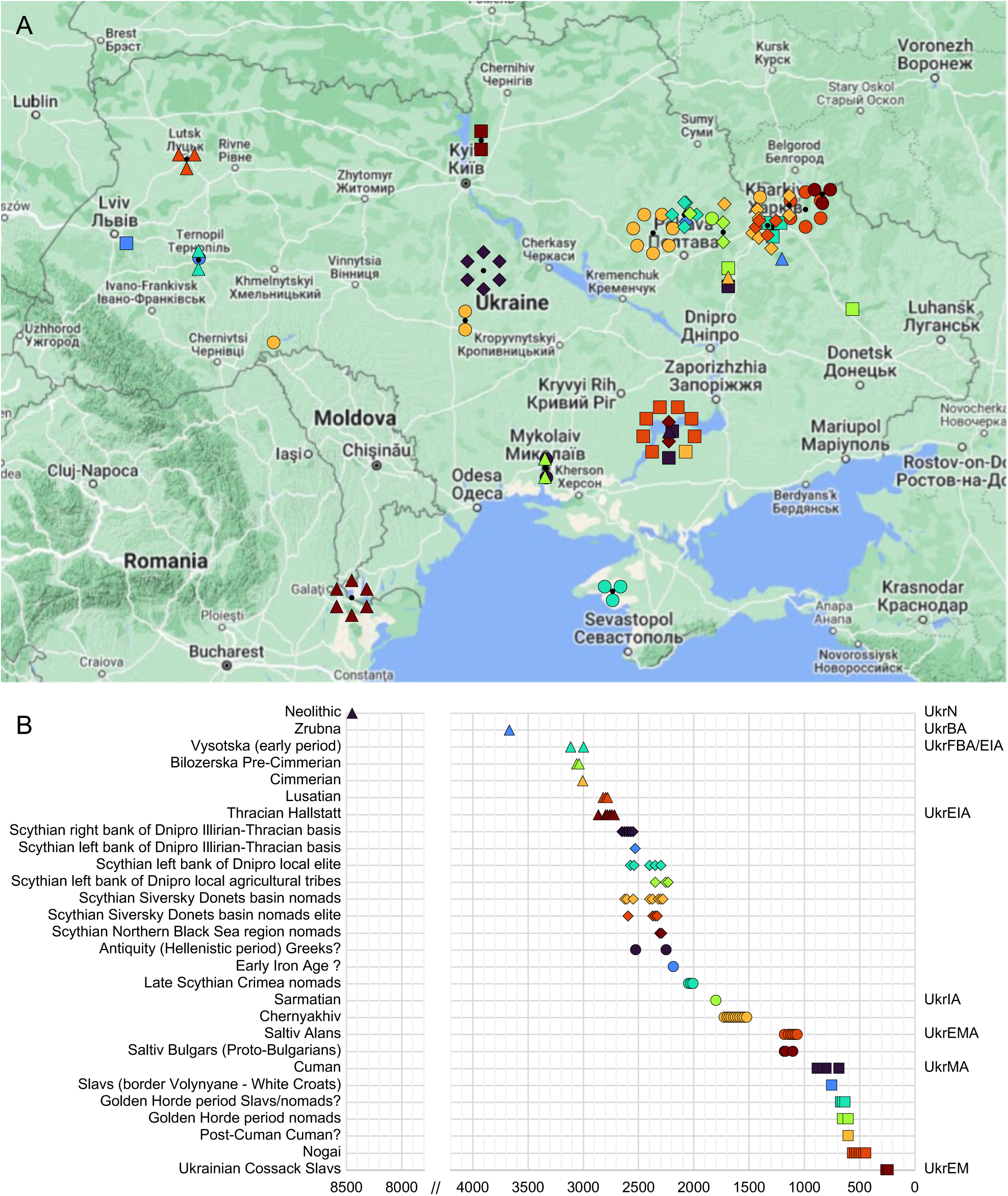
Map of the geographical locations of the individuals of this study and timeline showing the dates of individuals in archaeological groups. Symbols on (**A**) the map correspond to those on (**B**) the timeline. The dates (years before present) used on the timeline are midpoints of the 95% calibrated date estimates or archaeological date range estimates with jitter. See also Table S1.

**Table 1.** Archaeological information, genetic sex, mtDNA and Y chromosome haplogroups and average genomic coverage of the individuals of this study. Date (cal BCE/CE) – calibrated using OxCal v4.4.4 and IntCal20 atmospheric curve ^74,75^; MT hg – mitochondrial DNA haplogroup; Y hg – Y chromosome haplogroup; Av. Cov. – average genomic coverage. See also Tables S1, S2.

| Individual | Site | Region | Group | Date | Sex | MT hg | Y hg | Av. Cov. |
| --- | --- | --- | --- | --- | --- | --- | --- | --- |
| UKR008 | Mamay-Gora | Zaporizhzhia | UkrN_? | 7000–6000 BCE | XY | U4c2 | R1b-V88 | 0.087 |
| UKR055 | Sukha Gomilsha | Kharkiv | UkrBA_Zrubna | 1873–1566 cal BCE | XX | H5a1 | - | 0.035 |
| UKR170 | Petrykiv | Ternopil | UkrFBA/EIA_Vysotska_Early | 1300–800 BCE | XY | U2e2a1 | R1a-M198 | 0.054 |
| UKR171 | Petrykiv | Ternopil | UkrFBA/EIA_Vysotska_Early | 1278–1055 cal BCE | XY | U5a1a2b | R1a-M459 | 0.057 |
| UKR149 | Dykyi Sad | Mykolaiv | UkrFBA_Bilozerska_Pre-Cimmerian | 1200–1000 BCE | XX | U4a1 | - | 1.310 |
| UKR150 | Dykyi Sad | Mykolaiv | UkrFBA_Bilozerska_Pre-Cimmerian | 1200–1000 BCE | XY | N1a1a1a1 | R1a-M198 | 0.096 |
| UKR066 | Kumy | Kharkiv | UkrFBA/EIA_Cimmerian | 1195–919 cal BCE | XY | HV1a1a | Q1b-YP4000 | 0.451 |
| UKR167 | Rovantsi | Volyn | UkrFBA/EIA_Lusatian | 1000–700 BCE | XX | J1c3g | - | 1.950 |
| UKR168 | Rovantsi | Volyn | UkrFBA/EIA_Lusatian | 1000–700 BCE | XX | K1a4a1e | - | 0.852 |
| UKR169 | Rovantsi | Volyn | UkrFBA/EIA_Lusatian | 1000–700 BCE | XY | H1an1 | R1a-M459 | 0.085 |
| UKR000 | Kartal | Odesa | UkrEIA_ThracianHallstatt_2 | 900–798 cal BCE | XX | U4a2a | - | 1.170 |
| UKR001 | Kartal | Odesa | UkrEIA_ThracianHallstatt | 900–700 BCE | XX | K1a | - | 1.020 |
| UKR002 | Kartal | Odesa | UkrEIA_ThracianHallstatt | 900–700 BCE | XY | U5b1a | C1a-Z38888 | 0.422 |
| UKR005 | Kartal | Odesa | UkrEIA_ThracianHallstatt | 900–700 BCE | XY | T2 | ? | 0.026 |
| UKR006 | Kartal | Odesa | UkrEIA_ThracianHallstatt | 900–700 BCE | XX | U5a1h | - | 0.052 |
| UKR007 | Kartal | Odesa | UkrEIA_ThracianHallstatt | 996–831 cal BCE | XY | T1a1 | E1b-V13 | 0.313 |
| UKR035AB | Medvyn | Kyiv | UkrEIA_Scythian_RightDnipro_IIIThr | 700–600 BCE | XY | U5a1g1 | R1a-Z283 | 1.690 |
| UKR036 | Medvyn | Kyiv | UkrEIA_Scythian_RightDnipro_IIIThr_2 | 773–426 cal BCE | XY | C4b | R1a-Z645 | 0.738 |
| UKR039 | Medvyn | Kyiv | UkrEIA_Scythian_RightDnipro_IIIThr | 700–600 BCE | XY | U5a1a1a | R1a-M420 | 0.060 |
| UKR042 | Medvyn | Kyiv | UkrEIA_Scythian_RightDnipro_IIIThr | 779–539 cal BCE | XX | T2a2 | - | 0.170 |
| UKR043 | Medvyn | Kyiv | UkrEIA_Scythian_RightDnipro_IIIThr | 700–600 BCE | XX | U5a2b | - | 0.372 |
| UKR044 | Medvyn | Kyiv | UkrEIA_Scythian_RightDnipro_IIIThr | 700–600 BCE | XX | H6a1b | - | 0.424 |
| UKR078 | Bilsk hillfort | Poltava | UkrEIA_Scythian_LeftDnipro_IIIThr | 755–408 cal BCE | XX | H6a1b | - | 0.819 |
| UKR083 | Bilsk hillfort | Poltava | UkrEIA_Scythian_LeftDnipro_LocAgr | 500–300 BCE | XY | U4d3 | R1a-M198 | 0.080 |
| UKR087 | Bilsk hillfort | Poltava | UkrEIA_Scythian_LeftDnipro_LocEl | 650–600 BCE | XX | H3v+ | - | 0.400 |
| UKR088 | Bilsk hillfort | Poltava | UkrEIA_Scythian_LeftDnipro_LocEl | 761–420 cal BCE | XX | U2b | - | 0.457 |
| UKR089 | Bilsk hillfort | Poltava | UkrEIA_Scythian_LeftDnipro_LocEl | 500–300 BCE | XY | H+152 | E1b-V13 | 0.449 |
| UKR090 | Bilsk hillfort | Poltava | UkrEIA_Scythian_LeftDnipro_LocEl | 400–300 BCE | XY | H11a1 | E1b-V13 | 0.450 |
| UKR091 | Bilsk hillfort | Poltava | UkrEIA_Scythian_LeftDnipro_LocEl | 500–400 BCE | XY | H+152 | E1b-V13 | 0.327 |
| UKR095 | Kolomak | Kharkiv | UkrEIA_Scythian_LeftDnipro_LocAgr | 389–204 cal BCE | XY | J2b1a2a | R1a-Z93 | 0.699 |
| UKR096 | Kolomak | Kharkiv | UkrEIA_Scythian_LeftDnipro_LocAgr_2 | 382–199 cal BCE | XX | T2a1a | - | 1.030 |
| UKR105 | Kupievakha | Kharkiv | UkrEIA_Scythian_SivDon_Nom | 798–552 cal BCE | XY | J1d6 | R1a-YP5018 | 0.644 |
| UKR101 | Nyzhnia Gyivka | Kharkiv | UkrEIA_Scythian_SivDon_Nom | 400–300 BCE | XX | I1a1 | - | 0.375 |
| UKR114 | Nyzhnia Gyivka | Kharkiv | UkrEIA_Scythian_SivDon_Nom_2 | 500–400 BCE | XX | H4a1c | - | 0.290 |
| UKR104 | Grishkivka | Kharkiv | UkrEIA_Scythian_SivDon_Nom | 500–350 BCE | XY | H6a1b | R1a-Z645 | 0.919 |
| UKR109 | Vesele | Kharkiv | UkrEIA_Scythian_SivDon_Nom | 400–300 BCE | XY | H5a1 | Q1b-L330 | 0.530 |
| UKR110 | Vesele | Kharkiv | UkrEIA_Scythian_SivDon_Nom | 400–300 BCE | XY | U3a1b | R1b-Z2105 | 0.430 |
| UKR111 | Cheremushna | Kharkiv | UkrEIA_Scythian_SivDon_Nom_2 | 775–540 cal BCE | XY | K2 | R1a-YP5018 | 0.506 |
| UKR113 | Mala Rogozianka | Kharkiv | UkrEIA_Scythian_SivDon_Nom | 650–550 BCE | XY | HV0a | Q1b-L330 | 0.716 |
| UKR116 | Karavan | Kharkiv | UkrEIA_Scythian_SivDon_NomEl_3 | 775–516 cal BCE | XY | U2e1f1 | R1a-Z645 | 0.598 |
| UKR131 | Pisochyn | Kharkiv | UkrEIA_Scythian_SivDon_NomEl | 500–300 BCE | XY | I1c1a | R1b-M269 | 0.484 |
| UKR132 | Pisochyn | Kharkiv | UkrEIA_Scythian_SivDon_NomEl | 500–300 BCE | XY | X2f | R1b-L23 | 0.872 |
| UKR133 | Pisochyn | Kharkiv | UkrEIA_Scythian_SivDon_NomEl | 500–300 BCE | XY | T1b | R1b-Z2105 | 0.925 |
| UKR013 | Mamay-Gora | Zaporizhzhia | UkrEIA_Scythian_NBlaSea_Nom | 400–300 BCE | XX | HV2a3 | - | 0.398 |
| UKR014 | Mamay-Gora | Zaporizhzhia | UkrEIA_Scythian_NBlaSea_Nom | 400–300 BCE | XY | I4a | R1a-Z645 | 0.290 |
| UKR152 | Oleksandrivskyi<br>necropolis | Mykolaiv | UkrEIA_Antiquity_Greeks?_1 | 392–206 cal BCE | XY | HV1b | E1b-V13 | 0.818 |
| UKR153 | Oleksandrivskyi<br>necropolis | Mykolaiv | UkrEIA_Antiquity_Greeks?_2 | 746–401 cal BCE | XY | W6b | R1a-M459 | 0.573 |
| UKR174 | Petrykiv | Ternopil | UkrEIA_? | 359–104 cal BCE | XX | HV2 | - | 0.534 |
| UKR051 | Maslynny | Crimea | UkrEIA_LateScythian_Cri_Nom | 150–1 BCE | XX | U7a3a* | - | 0.648 |
| UKR052 | Maslynny | Crimea | UkrEIA_LateScythian_Cri_Nom | 150–1 BCE | XY | R1a1a | R1b-M269 | 0.066 |
| UKR053 | Maslynny | Crimea | UkrEIA_LateScythian_Cri_Nom | 150–1 BCE | XX | HV1a1 | - | 0.029 |
| UKR160 | Liubivka | Kharkiv | UkrIA_Sarmatian_SivDon | 1–300 CE | XX | H16+ | - | 1.220 |
| UKR045 | Lehedzyne | Cherkasy | UkrIA_Chernyakhiv_2 | 300–400 CE | XX | U4c1 | - | 0.375 |
| UKR047 | Lehedzyne | Cherkasy | UkrIA_Chernyakhiv_2 | 300–400 CE | XX | H2a2a | - | 0.138 |
| UKR049 | Komariv-1 | Chernivtsi | UkrIA_Chernyakhiv_3 | 169–338 cal CE | XX | HV0a | - | 0.433 |
| UKR102 | Zolochiv | Kharkiv | UkrIA_Chernyakhiv_2 | 300–400 CE | XY | H26 | E1b-V13 | 0.755 |
| UKR121 | Shyshaky | Poltava | UkrIA_Chernyakhiv_2 | 300–400 CE | XX | H1 | - | 0.347 |
| UKR122 | Shyshaky | Poltava | UkrIA_Chernyakhiv_1 | 300–400 CE | XX | K1b2b | - | 0.334 |
| UKR123 | Shyshaky | Poltava | UkrIA_Chernyakhiv_2 | 300–400 CE | XY | H7 | R1a-Z93 | 0.681 |
| UKR125 | Shyshaky | Poltava | UkrIA_Chernyakhiv_1 | 300–400 CE | XX | K1a4a1b | - | 0.309 |
| UKR126 | Shyshaky | Poltava | UkrIA_Chernyakhiv_1 | 131–325 cal CE | XY | H1a3 | R1a-Z645 | 0.389 |
| UKR128 | Shyshaky | Poltava | UkrIA_Chernyakhiv_2 | 229–361 cal CE | XX | K1a+ | - | 0.390 |
| UKR129 | Shyshaky | Poltava | UkrIA_Chernyakhiv_1 | 245–401 cal CE | XX | K1c2 | - | 0.296 |
| UKR134 | Verkhni Saltiv | Kharkiv | UkrEMA_Saltiv_Alans_2 | 800–900 CE | XX | A16 | - | 0.879 |
| UKR135 | Verkhni Saltiv | Kharkiv | UkrEMA_Saltiv_Alans_1 | 800–900 CE | XY | A+ | R1b-L23 | 0.422 |
| UKR136 | Verkhni Saltiv | Kharkiv | UkrEMA_Saltiv_Alans_1 | 800–900 CE | XY | W3 | R1b-M269 | 0.445 |
| UKR137 | Verkhni Saltiv | Kharkiv | UkrEMA_Saltiv_Alans_2 | 800–900 CE | XY | F2c1 | C2a-Y11606 | 0.359 |
| UKR138 | Verkhni Saltiv | Kharkiv | UkrEMA_Saltiv_Alans_1 | 800–900 CE | XY | H27+ | R1b-Z2105 | 0.441 |
| UKR139 | Verkhni Saltiv | Kharkiv | UkrEMA_Saltiv_Alans_1 | 671–874 cal CE | XX | W6 | - | 0.343 |
| UKR143 | Bochkove | Kharkiv | UkrEMA_Saltiv_Bulgars_1 | 800–900 CE | XX | U5a1a1 | - | 0.569 |
| UKR144 | Bochkove | Kharkiv | UkrEMA_Saltiv_Bulgars_2 | 671–874 cal CE | XX | B4c1a2a | - | 0.390 |
| UKR147 | Bochkove | Kharkiv | UkrEMA_Saltiv_Bulgars_1 | 683–883 cal CE | XY | D4 | R1a-M417 | 0.152 |
| UKR012 | Mamay-Gora | Zaporizhzhia | UkrMA_Cuman | 1100–1430 CE | XY | J1b1b1 | R1a-Z645 | 0.374 |
| UKR027 | Velyka Znamianka | Zaporizhzhia | UkrMA_Cuman | 1100–1200 CE | XY | H5a1 | N1a-Y10755 | 0.317 |
| UKR056 | Kumy | Kharkiv | UkrMA_Cuman_2 | 991–1149 cal CE | XX | B5b2a2 | - | 0.832 |
| UKR166 | Zvenigorod | Lviv | UkrMA_WhiteCroat_Slavs | 1100–1300 CE | XX | H1a | - | 1.040 |
| UKR068 | Donets hillfort | Kharkiv | UkrMA_GoldenHorde_Slav/Nom? | 1200–1400 CE | XX | U5b1e1a | - | 0.784 |
| UKR069 | Donets hillfort | Kharkiv | UkrMA_GoldenHorde_Slav/Nom? | 1200–1400 CE | XY | X2d1a | R1b-Z2105 | 0.531 |
| UKR070 | Donets hillfort | Kharkiv | UkrMA_GoldenHorde_Slav/Nom? | 1200–1400 CE | XY | U5a1a1a | ? | 0.019 |
| UKR063 | Kumy | Kharkiv | UkrMA_GoldenHorde_Nom | 1300–1400 CE | XX | U2e1b | - | 0.285 |
| UKR074 | Ploske | Donetsk | UkrMA_GoldenHorde_Nom | 1200–1400 CE | XX | A1a | - | 0.333 |
| UKR028 | Mamay-Surka | Zaporizhzhia | UkrMA_Post-Cuman_Cuman? | 1300–1400 CE | XX | J1c2e1 | - | 0.394 |
| UKR016 | Mamay-Gora | Zaporizhzhia | UkrMA_Nogai_1 | 1400–1500 CE | XY | M7c1a1a1 | R1b-Y14051 | 0.322 |
| UKR017 | Mamay-Gora | Zaporizhzhia | UkrMA_Nogai_2 | 1400–1500 CE | XY | C4a1a4a | C2a-M504 | 0.764 |
| UKR018 | Mamay-Gora | Zaporizhzhia | UkrMA_Nogai_3 | 1400–1500 CE | XX | B4d1'2'3 | - | 0.260 |
| UKR020 | Mamay-Gora | Zaporizhzhia | UkrMA_Nogai_2 | 1400–1500 CE | XY | M7b1a1a1 | J1a-Y5321 | 0.607 |
| UKR021 | Mamay-Gora | Zaporizhzhia | UkrMA_Nogai_3 | 1400–1500 CE | XX | C4a1a4a | - | 0.633 |
| UKR022 | Mamay-Gora | Zaporizhzhia | UkrMA_Nogai_2 | 1400–1500 CE | XX | U2e2a1a2 | - | 0.387 |
| UKR024 | Mamay-Gora | Zaporizhzhia | UkrMA_Nogai_1 | 1400–1500 CE | XY | M65a+ | G2a-Z6679 | 0.523 |
| UKR164 | Vypovziv | Chernigiv | UkrEM_Cossack_Slavs | 1600–1800 CE | XX | H3a1 | - | 0.444 |
| UKR165 | Vypovziv | Chernigiv | UkrEM_Cossack_Slavs | 1600–1800 CE | XX | U4a2b | - | 0.272 |

### Southern European ancestry is present in Ukraine in the Late Bronze Age to pre-Scythian Iron Age period

The mitochondrial DNA (mtDNA) of the Late Bronze Age and pre-Scythian Iron Age (LBAEIA; 3,000–700 BCE) individuals belonged to haplogroups (hgs) U, HV, H, T, K, J and N1a (Table 1, Table S1), while the Y chro-mosomes (chrY) of most males belonged to hg R1a (Table 1, Tables S1–S2), as has been shown previously for much of northern Europe after the steppe migrations ^1,2^. However, the Cimmerian male belonged to chrY hg Q1b-YP4004 (common in Central Asia and Siberia ^44^, but not in Europe ^45^), and one Thracian Hallstatt individual to C1-Y83490 (sub-branches of C-V20 are found in ancient Europeans in the Stone Age ^46–48^, but rare in Europe today ^49^) (Table 1, Tables S1–S2).

We performed principal component analysis (PCA) and ADMIXTURE analysis using autosomal data of modern individuals (“modern” ADMIXTURE) (Table S3) and projecting ancient individuals (Table S4) onto the components. ADMIXTURE analysis was also performed without modern individuals, using only ancient individuals with up to 90% missing positions after pruning the dataset to decrease linkage disequilibrium (maximum 5 individuals per group) (Methods, Table S4), and later also projecting all ancient individuals onto the components (“ancient” ADMIXTURE). The Zrubna individual of this study (1,873–1,566 cal BCE) and the previously published Zrubna individual from Ukraine (781–511 cal BCE) ^7^ (UkrBA_Zrubna) cluster on PCA with individuals from present-day Russia associated with the same culture (Figure 3A, Figures S1–S2A).

**Figure 3.**
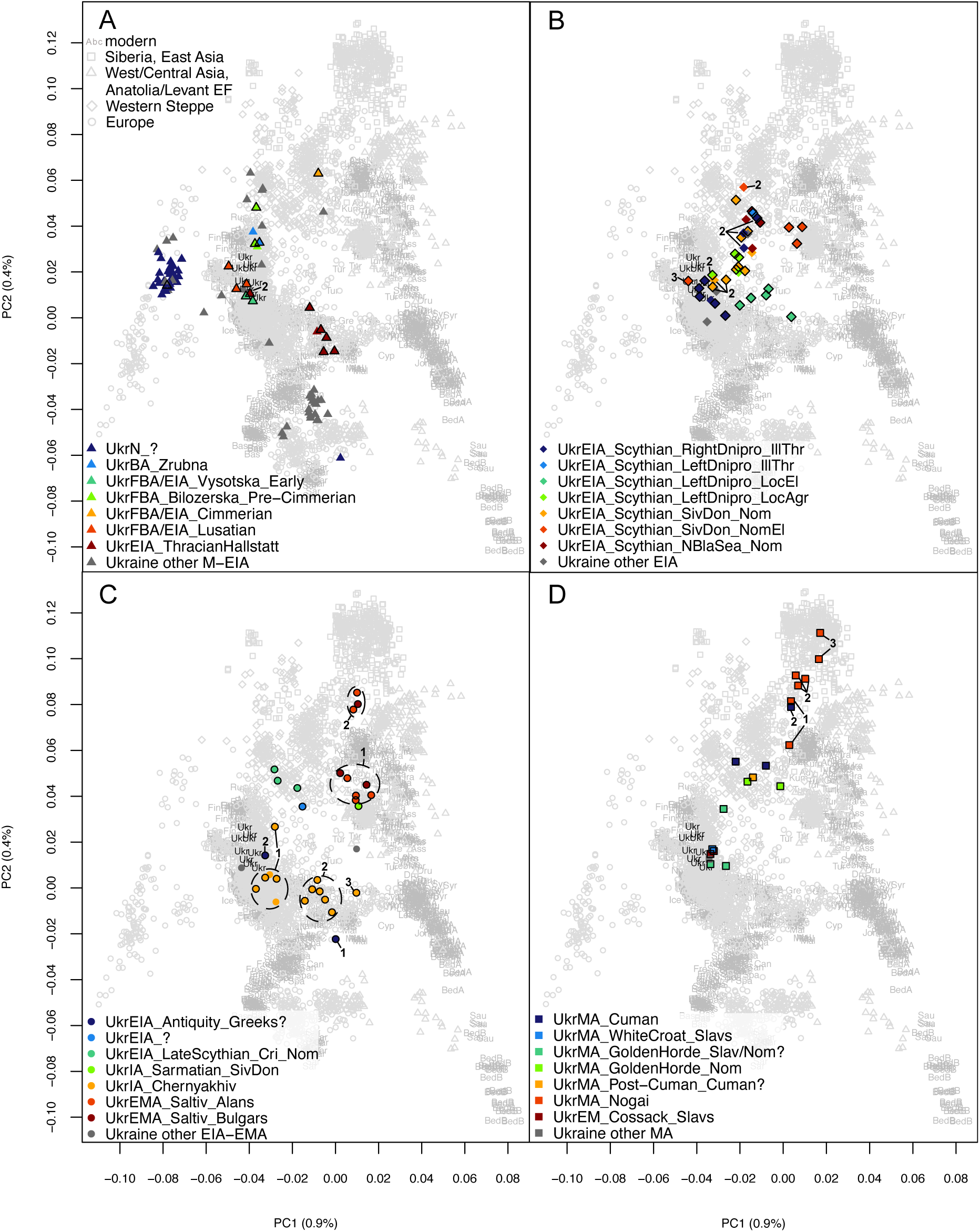
Principal component analysis results. Principal component analysis results of modern West Eurasians with ancient individuals projected onto the first two components (PC1 and PC2). Ukrainian groups from (**A**) Late Bronze Age and pre-Scythian Iron Age (3,000–700 BCE), (**B**) the Scythian period of Early Iron Age (700–300 BCE), (**C**) post-Scythian Iron Age until Early Middle Ages (400 BCE–900 CE), (**D**) Middle Ages and early modern period (900–1,800 CE). Newly reported individuals are indicated with a black outline. Genetic subgroups used in subsequent analyses are indicated with black numbers. Modern Ukrainians are shown in black. See also Figures S1–S2, Tables S3–S4.

In both PCA and ADMIXTURE, Zrubna individuals are similar to Yamna individuals, but some Zrubna genomes show signs of admixture with early farmers (Figure 3A, Figures S1, S2A, S3, S4). Bilozerska Pre-Cimmerians (archaeologically dated to 1,200–1,000 BCE, one radiocarbon dated to 1281–1058 cal BCE ^7^; UkrFBA_Bilozerska_Pre-Cimmerian) appear genetically similar to Zrubna individuals (Figures 3A, Figures S1, S3, S4CF). Early Vysotska (1,300–800 BCE, one 1,278–1,055 cal BCE; UkrFBA/EIA_Vysotska_Early) and Lusatian (1,000–700 BCE; UkrFBA/EIA_Lusatian) individuals appear similar to Northern and Eastern European individuals from the Iron Age to modern times (including modern Ukrainians) (Figures 3A, 4, Figures S1, S3, S4). The Cimmerian individual (1,195– 919 cal BCE; UkrFBA/EIA_Cimmerian) clusters on the PCA plot with Western Steppe individuals (Figure 3A, Figure S1), including previously published Cimmerians from Moldova (Figure S2A), but with a bigger East Asian genetic influence compared to Bilozerska Pre-Cimmerians (Figure 4, Figures S3–S4). Most of the Early Iron Age Thracian Hallstatt individuals (900–700 BCE, two 996–830 cal BCE; UkrEIA_ThracianHallstatt) cluster with Southern Europeans on PCA (Figure 3A, Figure S1) while previously published Hallstatt individuals from the Czech Republic cluster with Central European individuals (Figure S2A). A bigger early farmer influence compared to earlier individuals from Ukraine can also be seen on ADMIXTURE (Figure 4, Figures S3–S4). One Thracian Hallstatt individual (900– 798 cal BCE; UkrEIA_ThracianHallstatt_2) is similar to Early Vysotska and Lusatian individuals in both analyses (Figures 3A, 4, Figures S1, S3, S4).

**Figure 4.**
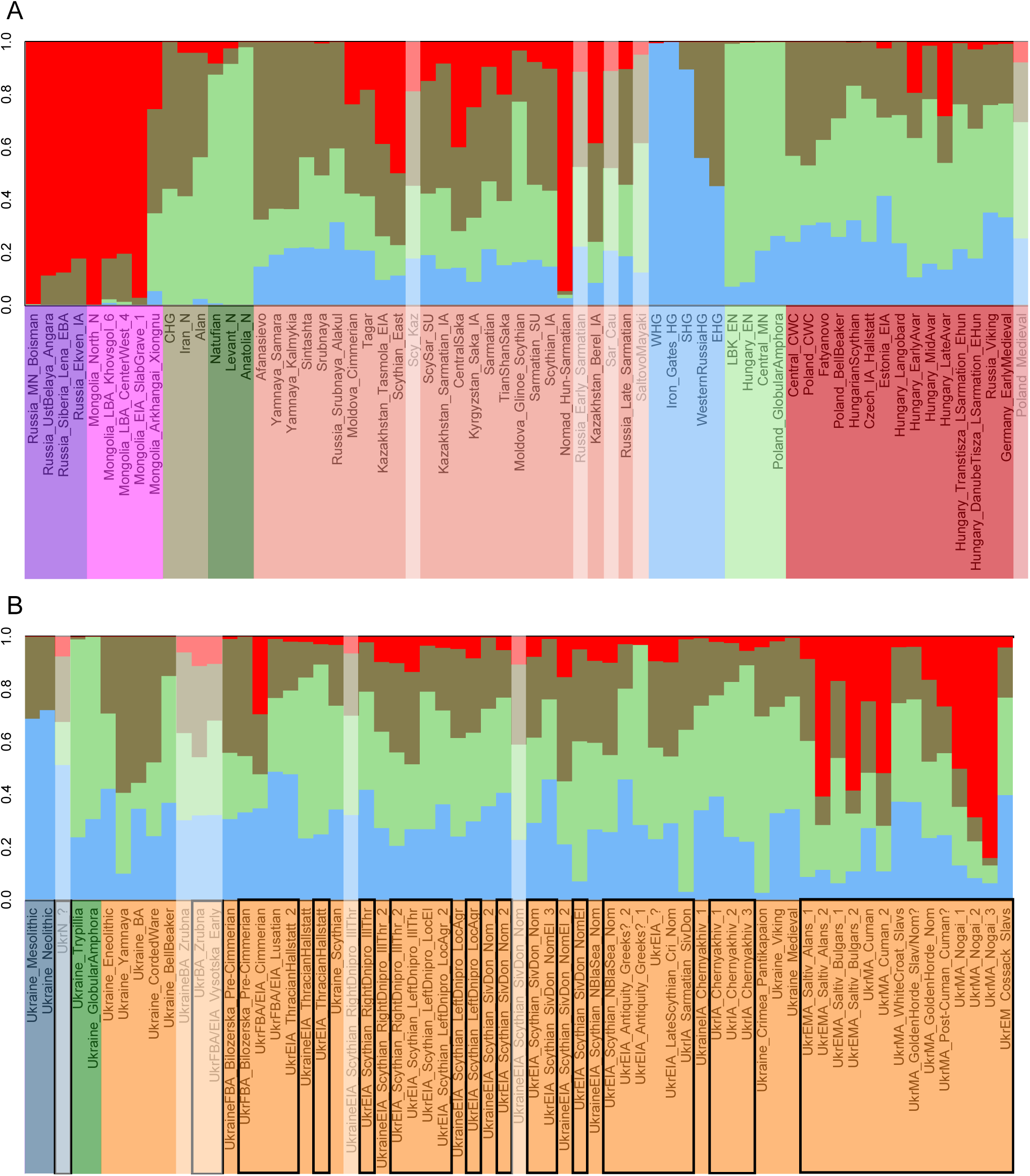
Subset of ADMIXTURE analysis results with genetic structure calculated on ancient individuals. Population averages of (**A**) Eurasia-wide, (**B**) Ukrainian ancient groups at K4. See also Figures S3–S4, Tables S3–S4.

Then, we tested the cladality of the groups assigned based on archaeological context or identified in PCA and ADMIXTURE by calculating f4 statistics of the from f4(Mbuti, ancient group; Ukrainian group 1, Ukrainian group 2) with a wide set of ancient groups (Table S4) and considering the Ukrainian groups cladal if at least 95% of the f4 results were not significantly different from 0 (−3 ≤ |Z| ≤ 3). The cladality of UkrBA_Zrubna and UkrFBA_Bilozerska_Pre-Cimmerian is confirmed, but UkrFBA/EIA_Vysotska_Early and UkrFBA/EIA_Lusatian are non-cladal with Lusatian having an additional affinity with some Siberian, East/West/Central Asian, Western Steppe and post-steppe migration European groups (but not with any European early farmers) (Tables S5–S6). UkrEIA_ThracianHallstatt_2 is cladal with contemporary UkrFBA/EIA_Lusatian (Tables S5–S6). Furthermore, we tested the cladality of the groups of this study with previously published groups that are associated with the same cultures and saw that UkrBA_Zrubna is cladal with AADR group Russia_Srubnaya_Alakul ^16^ and UkrEIA_ThracianHallstatt_2 with Czech_IA_Hallstatt_2 (an “outlier” individual of AADR group Czech_IA_Hallstatt ^50^), but UkrFBA/EIA_Cimmerian is non-cladal with AADR group Moldova_Cimmerian ^16^, the latter having relatively more affinity with some Siberian and East Asian groups (Tables S5–S6).

We used qpAdm to model the ancestry proportions of the archaeological/genetic groups. First, we tested different combinations of an early farmer group (5 total), a Yamna-associated group (3 total) and an East Asian (Mongolian) group (3 total) to find the distal sources that can be used to model the highest number of the groups of this study with significant results (Table S7). The model including Middle Neolithic individuals from Germany (AADR_Central_MN ^8^), Yamna-associated individuals from Ukraine (AADR_Ukraine_Yamnaya ^6^) and Slab-grave culture-associated individuals from Mongolia (AADR_Mongolia_EIA_SlabGrave_1 ^51^, AADR_Mongolia_EIA_SlabGrave used for brevity) as sources is among the models with the highest number of significant plausible results (27 out of 39 modelled groups) (Table S7). The best performing model with two Ukrainian sources includes Trypillia culture-associated individuals from Ukraine (AADR_Ukraine_Trypillia ^6,36^), AADR_Ukraine_Yamnaya and AADR_Mongolia_SlabGrave and gives significant plausible results for 26 out of 39 groups (Table S7). When models with one or two of the sources dropped are included, both up to three source models described above (AADR_Central_MN/AADR_Ukraine_Trypillia, AADR_Ukraine_Yamnaya, AADR_Mongolia_SlabGrave) produce a significant plausible result for 28 out of the 39 groups (Figure 5A, Figure S5A, Tables S8–S9). Four groups of this study (UkrFBA/EIA_Lusatian, UkrEIA_Scythian_LeftDnipro_LocAgr_2, UkrMA_WhiteCroat_Slavs, UkrMA_Nogai_2) do not produce a significant result with any of the tested three-source models (Table S7). All other groups, except for UkrEMA_Saltiv_Alans_1, give a significant result when modelled from AADR_Mongolia_SlabGrave, AADR_Ukraine_Yamnaya and AADR groups Central_MN/Poland_GlobularAmphora/Ukraine_GlobularAmphora/Ukraine_Trypillia ^6,8,36,52^ with the point estimate of the early farmer ancestry proportion differing by a maximum of 6% between the highest p-value model and the model with AADR_Ukraine_Trypillia (Table S7). Because of the small variability in the proportions, we use the model with AADR_Ukraine_Trypillia to describe all groups (Figure 5A, Table S7) and provide the model with AADR_Central_MN as comparison (Figure S5A, Table S9). Furthermore, we show the highest p-value three-way model out of all 315 tested models for each group (Figure S5B, Table S10). None of the UkrBA_Zrubna and UkrFBA/EIA_Vysotska_Early individuals have enough data (>100,000 SNPs) to be modelled, but UkrFBA_Bilozerska_Pre-Cimmerian can be modelled as mostly AADR_Ukraine_Yamnaya, some AADR_Ukraine_Trypillia and a small amount of AADR_Mongolia_SlabGrave (79–89% + 9–18% + 1–5%) and UkrFBA/EIA_Cimmerian as mostly AADR_Ukraine_Yamnaya and some AADR_Mongolia_SlabGrave (61–66% + 0% + 34–39%) (Figure 5A, Table S8). UkrEIA_ThracianHallstatt can be put together from some AADR_Ukraine_Yamnaya and mostly AADR_Ukraine_Trypillia, while UkrEIA_ThracianHallstatt_2 the other way around (19–25% + 75– 81% + 0% and 60–69% + 31–40% + 0%, respectively) (Figure 5A, Table S8).

**Figure 5.**
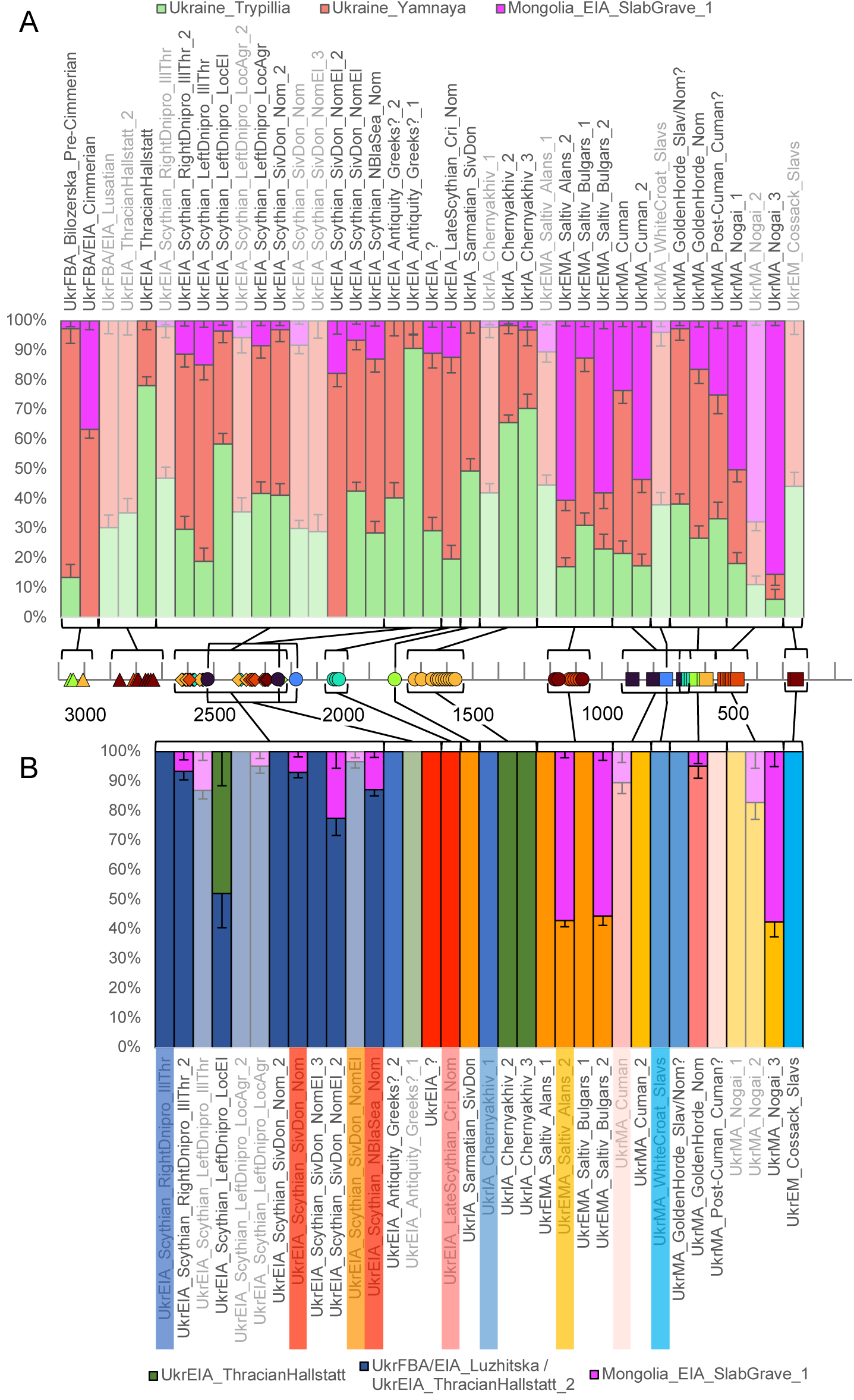
qpAdm admixture modelling results. (**A**) distal qpAdm models of admixture between AADR groups Ukraine_Trypillia, Ukraine_Yamnaya and Mongolia_SlabGrave, (**B**) proximal qpAdm models, tested using the autosomal positions of the 1240K dataset. Models with non-significant pvalues (p<0.05) are semi-transparent. The dates used on the timeline are midpoints of the 95% calibrated date estimates or archaeological date range estimates with jitter. See also Figure S5, Tables S3–S4, S7– S11.

To assess whether there is a sex bias in the ancestry distribution of any groups in this study, we calculated outgroup f3 statistics with a wide set of ancient groups (Table S4) on both autosomal (Table S12) and chrX (Table S13) data – the former being more representative of the male and the latter of the female lineage – and compared the results with each other. The Zrubna-associated group shows a trend of higher similarity to Levantine/Anatolian and European early farmers on chrX, and to Neolithic to Early Bronze Age Western Steppe groups and European HGs on autosomes (Figure S6A), consistent with admixture after the Yamna migration into Europe taking place mostly between males from the steppe (with a large proportion of Eastern HG ancestry) and local early farmer females, as has been inferred for Corded Ware Culture groups ^53–56^.

### The Scythian period of Early Iron Age is characterised by Eastern European and Western Steppe ancestries

The mtDNA variation during the Scythian period (SEIA; 700–300 CE) is similar to the earlier time window, but additionally includes mtDNA hgs I and X, and one individual carries hg C4 (Table 1, Table S1), which today has a mostly Asian distribution ^57^. The chrY lineages include R1a, R1b, E1b as well as Central/East Asian ^44^ Q1b-L330 (Table 1, Tables S1–S2).

Kinship analysis reveals some close genetic relationships among the Scythian period individuals in this study and previously published individuals from the same sites from Järve et al. 2019 (7). Burial 1 in kurgan 22 in Medvyn in Kyiv region includes two genetically identical samples (UKR035 and UKR038) that most likely come from the same individual (male, mtDNA hg U5a1g1, chrY hg R1a) (Figure S7A, Methods) and were later merged together for further analyses (UKR035AB). However, the burial also includes another male (MJ-14; mtDNA hg H6a1b, chrY hg R1a) who is a first degree relative of the previously described male and also of a female individual (UKR044; mtDNA hg H6a1b), while the latter two are not closely related (Figure S7A). This means that MJ-14 is likely the son of UKR035AB and UKR044 (Figure S7A). Furthermore, there is another pair of first degree relatives from the same site – two males, MJ-33 (mtDNA hg U5a2a2a, chrY hg R1a) and UKR036 (mtDNA hg C4b, chrY hg R1a), who could be father and son in either direction (Figure S7A). At the Bilsk hillfort in Poltava region, the analysis identified another pair of genetically identical samples (UKR089 and UKR091; mtDNA hg H+152, chrY hg E1b), but these come from different kurgans that were excavated in different years, so it is likely that they were identical twins (Figure S7B, Methods). These inferred twins are first degree related to another male (UKR090; mtDNA hg H11a1, chrY hg E1b) who is likely their father (Figure S7B).

Scythian period individuals (nine of whom are radiocarbon dated to 798–199 cal BCE) were divided into groups based on their geographic location: right (i.e. west) or left (i.e. east) bank of Dnipro and Siversky Donets basin in the forest-steppe, Northern Black Sea region in the steppe. The groups were further divided based on the demographic association inferred from archaeological context: Illirian-Thracian, local or nomad, agriculturalist or elite. The groups are named using the following structure, separated by underscores: time period, cultural association, geographic location, demographic association, genetic subgroup (where relevant) (Figure 2, Table 1, Table S1, Methods). Most of the Scythian individuals from the right bank of Dnipro with Illirian-Thracian associations (UkrEIA_Scythian_RightDnipro_IllThr), one individual from local agricultural tribes of the left bank of Dnipro (UkrEIA_Scythian_LeftDnipro_LocAgr_2), some of the non-elite nomad individuals from Siversky Donets basin (UkrEIA_Scythian_SivDon_Nom_2) and one elite nomad individual from the same area (UkrEIA_Scythian_SivDon_NomEl_3) are similar to previous Early Vysotska and Lusatian individuals and also modern Ukrainians in both PCA and ADMIXTURE (Figures 3B, 4, Figures S1, S2B, S3, S4). The rest of the Scythian individuals from the right bank of Dnipro with Illirian-Thracian associations (UkrEIA_Scythian_RightDnipro_IllThr_2) and from local agricultural groups of the left bank of Dnipro (UkrEIA_Scythian_LeftDnipro_LocAgr) are more similar to Western Steppe individuals (including previously published Scythian-related individuals from the region) (Figures 3B, 4, Figures S1, S2B, S3, S4). The same is true for most of the non-elite nomad individuals from Siversky Donets basin (UkrEIA_Scythian_SivDon_Nom) and one elite nomad individual from the same region (UkrEIA_Scythian_SivDon_NomEl_2), the individual from the left bank of Dnipro with Illirian-Thracian associations (UkrEIA_Scythian_LeftDnipro_IllThr), as well as the four steppe nomads from the Northern Black Sea region (UkrEIA_Scythian_NBlaSea_Nom) (Figures 3B, 4, Figures S1, S2B, S3, S4). Local elite individuals from the left bank of Dnipro (UkrEIA_Scythian_LeftDnipro_LocEl) have greater genetic affinity to Southern European individuals (somewhat similarly to Scythians from Moldova) (Figures 3B, 4, Figures S1, S2B, S3, S4). Most of the elite nomads from the Siversky Donets basin (UkrEIA_Scythian_SivDon_NomEl) (n=3) share highest similarity with individuals from the Caucasus (Figures 3B, 4, Figures S1, S3, S4).

The f4-based cladality test supports UkrEIA_Scythian_RightDnipro_IllThr being cladal with UkrEIA_Scythian_SivDon_Nom_2 and UkrEIA_Scythian_LeftDnipro_LocAgr_2, but not with UkrEIA_Scythian_SivDon_NomEl_3, which has relatively more affinity with some East Asian groups and European HGs (Tables S5–S6). Also, UkrEIA_Scythian_SivDon_Nom is cladal with UkrEIA_Scythian_RightDnipro_IllThr_2, UkrEIA_Scythian_LeftDnipro_LocAgr and UkrEIA_Scythian_NBlaSea_Nom. However, UkrEIA_Scythian_SivDon_Nom shares more with several European groups compared to UkrEIA_Scythian_LeftDnipro_IllThr and less with some Siberian and East Asian groups than UkrEIA_Scythian_SivDon_NomEl_2. Importantly, UkrEIA_Scythian_RightDnipro_IllThr is cladal with preceding UkrEIA_ThracianHallstatt_2 and UkrFBA/EIA_Vysotska_Early, as well as HungarianScythian_1 (the main subgroup of AADR group HungarianScythian ^17^) (Tables S5–S6). Furthermore, UkrEIA_Scythian_SivDon_NomEl is cladal with Scy_Kaz_2 (one individual of AADR group Scy_Kaz ^7^) (Tables S5–S6).

With distal qpAdm modelling, UkrEIA_Scythian_RightDnipro_IllThr and UkrEIA_Scythian_SivDon_Nom_2 can be modelled as approximately half AADR_Ukraine_Yamnaya and half AADR_Ukraine_Trypillia, with a small amount of AADR_Mongolia_SlabGrave ancestry (50– 58% + 40–48% + 1–4% on average) (Figure 5A, Table S8). UkrEIA_Scythian_SivDon_NomEl has slightly more AADR_Mongolia_SlabGrave ancestry (48–54% + 40–46% + 5–8%), while UkrEIA_Scythian_SivDon_NomEl_3 can be modelled without it (65–77% + 23–35% + 0%) (Figure 5A, Table S8). UkrEIA_Scythian_RightDnipro_IllThr_2, _LeftDnipro_LocAgr, _SivDon_Nom and _NBlaSea_Nom can be put together from mostly AADR_Ukraine_Yamnaya, some AADR_Ukraine_Trypillia and some additional AADR_Mongolia_SlabGrave ancestry (53–61% + 29–36% + 9–12% on average), while UkrEIA_Scythian_LeftDnipro_IllThr has slightly less of AADR_Ukraine_Trypillia ancestry and more of the other two (61–71% + 14–23% + 13–17%) (Figure 5A, Table S8). UkrEIA_Scythian_SivDon_NomEl_2, however, can be modelled without AADR_Ukraine_Trypillia ancestry (78–87% + 0% + 13–22%), while UkrEIA_Scythian_LeftDnipro_LocEl needs more than other Scythian groups (34–42% + 55–62% + 2– 5%) (Figure 5A, Table S8).

Next, we also modelled the groups of this study starting from the Scythian period using relevant groups from earlier periods as sources (proximal qpAdm modelling) (Table S11). The analysis showed that UkrEIA_Scythian_RightDnipro_IllThr, _SivDon_Nom_2 and _SivDon_NomEl_3 can be modelled as 100% UkrFBA/EIA_Lusatian or UkrEIA_ThracianHallstatt_2 ancestry, while UkrEIA_Scythian_RightDnipro_IllThr_2, _SivDon_Nom, _SivDon_NomEl_2 and _NBlaSea_Nom show an additional input from AADR_Mongolia_SlabGrave (4–10%, 5–9%, 17–28% and 11–15%, respectively) (Figure 5B, Table S11). UkrEIA_Scythian_LeftDnipro_LocEl can be modelled as approximately half UkrEIA_ThracianHallstatt_2 and half UkrEIA_ThracianHallstatt (41–63% + 37– 59%) ancestry (Figure 5B, Table S11).

### Appearance of ancestry profiles from Southern Europe, the Caucasus and Central Asia during Post-Scythian Iron Age (Hellenistic period) until Early Middle Ages

The mtDNA lineages carried by individuals from post-Scythian Iron Age (Hellenistic period) and early medieval period (IAEMA; 400 BCE–900 CE) in Ukraine include many of those already seen in earlier periods, as well as mtDNA hg W and one case of R1a1a (Table 1, Table S1), which is found in the Caucasus ^58^, but very rarely observed in modern-day Eastern Europe ^59^. Furthermore, Saltiv Culture associated individuals carry lineages belonging, among others, to mtDNA hgs A, B, D and F (Table 1, Table S1), all frequent in East Asian populations today and rare in Europe ^60–63^. The chrY lineages present in the individuals from this period are again R1a, R1b and E1b, and Central East Asian ^64^ C2a-Y11606 in one Saltiv-associated individual (Table 1, Tables S1–S2).

The dataset includes two individuals that have been tentatively associated with Greeks of the Hellenistic period (Table 1, Table S1). One of the individuals (392–206 cal BCE; UkrEIA_Antiquity_Greeks?_1) has highest affinity with Southern Europeans, but the other (746–401 cal BCE; UkrEIA_Antiquity_Greeks?_2) is most similar to Nortern and Eastern European individuals (including modern Ukrainians) (Figures 3C, 4, Figures S1, S2C, S3, S4). An individual with no specific archaeological association (359–104 cal BCE; UkrEIA_?) and all three available Late Scythians (150– 1 BCE; UkrEIA_LateScythian_Cri_Nom) have highest affinity with Western Steppe individuals, including some of the Scythian period individuals of this study (see the previous subsection) (Figures 3C, 4, Figures S1, S3, S4). The Chernyakhiv Culture associated individuals (300–400 CE, seven 131– 530 cal CE) can be divided into three genetic subgroups – 6 individuals are most similar to Eastern/Central Europeans (UkrIA_Chernyakhiv_1), 6 to continental Southern Europeans (clustering with Thracian Hallstatt associated individuals of this study; Figure S1) (UkrIA_Chernyakhiv_2) and 1 individual has an even more southern genetic profile, clustering with modern Cypriots (UkrIA_Chernyakhiv_3) (Figures 3C, 4, Figures S1, S3, S4). Importantly, most of the individuals are from one site (Shyshaky, Poltava region), but are divided between the first and second genetic subgroup (Table 1, Table S1). The one available Sarmatian (1–300 CE; UkrIA_Sarmatian_SivDon) and most of the Saltiv Culture associated Alans and Bulgars (800–900 CE, two 671–883 cal CE; UkrEMA_Saltiv_Alans/Bulgars_1) have the highest affinity with individuals from the Caucasus, similarly to previously published Saltiv Culture individuals and Alans, and most of the Siversky Donets basin elite nomads from this study, but not with previously published Sarmatians (Figures 3C, 4, Figures S1, S2C, S3, S4). Three of the Saltiv Culture associated Alans and Bulgars from the same sites as the previously mentioned individuals (800–900 CE, one 671–874 cal CE; UkrEMA_Saltiv_Alans/Bulgars_2) are most similar to Central Asians, unlike any previously published Saltiv or Alan associated individual (Figures 3C, 4, Figures S1, S2C, S3, S4).

The f4-based cladality tests confirm that Alans and Bulgars are cladal with each other within genetic subgroups 1 and 2 (Tables S5–S6). Furthermore, UkrEIA_Antiquity_Greeks?_2 is cladal with UkrIA_Chernyakhiv_1, as is UkrEIA_? with UkrEIA_LateScythian_Cri_Nom, but UkrIA_Sarmatian_SivDon is non-cladal with UkrEMA_Saltiv_Alans_1, the latter sharing more ancestry with some Siberian and East Asian groups (Tables S5–S6). Importantly, UkrIA_Chernyakhiv_1 is cladal with preceding UkrEIA_Scythian_RightDnipro_IllThr, as is UkrEIA_LateScythian_Cri_Nom with preceding UkrEIA_Scythian_SivDon_Nom and UkrEMA_Saltiv_Alans_1 with preceding UkrEIA_Scythian_SivDon_NomEl (Tables S5–S6). Interestingly, UkrEMA_Saltiv_Alans_1 is cladal with AADR group SaltovoMayaki ^17^ (from Russian Caspian Steppe), but not with AADR group Alan ^17^ (from Russian Caucasus), the latter having more affinity with West/Central Asian, Western Steppe and European groups (Tables S5–S6).

Distal qpAdm modelling indicates that UkrEIA_Antiquity_Greeks?_1 can be modelled as a small amount of AADR_Ukraine_Yamnaya and otherwise almost entirely AADR_Ukraine_Trypillia (5–14% + 86–95% + 0%), while UkrEIA_Antiquity_Greeks?_2 and UkrIA_Sarmatian_SivDon model as near equal proportions of the same sources (54–65% + 35–46% + 0% and 47–55% + 45–53% + 0%, respectively) (Figure 5A, Table S8). UkrIA_Chernyakhiv_1 can be put together from mostly AADR_Ukraine_Yamnaya, slightly less of AADR_Ukraine_Trypillia and a small amount of AADR_Mongolia_SlabGrave ancestry (52–59% + 39–45% + 1–4%) while UkrIA_Chernyakhiv_2 and _3 have more AADR_Ukraine_Trypillia ancestry (30–35% + 63–68% + 1–3% and 21–32% + 66–75% + 1–5%, respectively) (Figure 5A, Table S8). UkrEIA_?, UkrEIA_LateScythian_Cri_Nom and UkrEMA_Saltiv_Bulgars_1 can be explained by combining ancestry from mostly AADR_Ukraine_Yamnaya, some AADR_Ukraine_Trypillia and slightly less of AADR_Mongolia_SlabGrave (52–73% + 15–35% + 9–15% overall) (Figure 5A, Table S8). On the other hand, UkrEMA_Saltiv_Alans_2 and _Bulgars_2 can be modelled as around equal proportions of AADR_Ukraine_Yamnaya and AADR_Ukraine_Trypillia, but mostly AADR_Mongolia_SlabGrave ancestry (16–25% + 16–24% + 57–61% on average) (Figure 5A, Table S8).

Using proximal qpAdm modelling, UkrEIA_Antiquity_Greeks?_2 and UkrIA_Chernyakhiv_1 can be modelled as 100% UkrEIA_Scythian_RightDnipro_IllThr while UkrEIA_? and UkrEIA_LateScythian_Cri_Nom as 100% UkrEIA_Scythian_SivDon_Nom or UkrEIA_Scythian_NBlaSea_Nom (Figure 5B, Table S11). UkrIA_Sarmatian_SivDon, UkrEMA_Saltiv_Alans_1 and _Bulgars_1 can be put together from 100% UkrEIA_Scythian_SivDon_NomEl, but UkrEMA_Saltiv_Alans_2 and _Bulgars_2 also show 54–59% AADR_Mongolia_SlabGrave (Figure 5B, Table S11). Chernyakhiv_2 and _3 can be modelled as 100% UkrEIA_ThracianHallstatt (Figure 5B, Table S11).

The outgroup f3 analysis reveals that UkrIA_Chernyakhiv_3 has more affinity with Levantine/Anatolian early farmers on chrX compared to autosomes (Figure S6B, Tables S12–S13), suggesting that its similarity to southern populations comes more from the female rather than the male mediated migration.

### East Asian as well as Eastern European genomes during the Middle Ages and early modern period

The mtDNA variation in individuals from medieval to early modern (MAEM; 900–1,800 CE) Ukraine includes lineages that are frequent in Europe (mtDNA hgs U, H, J and X2; Table 1, Table S1) as well as some that are more frequent in Asia (mtDNA hgs A, B, C and for the first time in this study also M65a and M7; Table 1, Table S1). The chrY lineages include those that are common in modern Eastern Europe (R1a, R1b, N1a-Y10755 and G2a-Z6679), as well as lineages frequent in Siberia and East Asia (C2a-M504) ^64^ or the Near East (J1a-P58) ^65^ (Table 1, Tables S1–S2).

Two Cuman period individuals (900–1,400 CE; UkrMA_Cuman), one from the Post-Cuman period (1,300–1,400 CE; UkrMA_Post-Cuman_Cuman?) and two Golden Horde related nomads (1,200–1,400 CE; UkrMA_GoldenHorde_Nom) cluster with Western Steppe individuals on PCA, but show more of the “East Asian” ancestry component using ADMIXTURE, when compared to preceding Scythians (Figures 3D, 4, Figures S1, S3, S4). Another of the Cuman-associated individuals (991–1,149 cal CE; UkrMA_Cuman_2) clusters with Central Asians, showing a larger eastern influence (Figures 3D, 4, Figures S1, S3, S4). The White Croat Slav (1,100–1,300 CE; UkrMA_WhiteCroat_Slavs) and Golden Horde period individuals whose archaeological associations do not distinguish between Slavs and nomads (1,200–1,400 CE; UkrMA_GoldenHorde_Slav/Nom?), are similar to the first genetic subgroup of Chernyakhiv individuals, and also to modern Ukrainians (Figures 3D, 4, Figures S1, S3, S4). The Nogai-associated individuals of this study (1,400–1,500 CE) from Mamay-Gora in the Zaporizhzhia region can be divided into three genetic subgroups, the first of which (UkrMA_Nogai_1) is similar to the genetically more eastern Saltiv and Cuman associated individuals, while we infer an even larger eastern ancestry component in the remaining two subgroups (UkrMA_Nogai_2/3), the third clustering with Mongolians on PCA (Figures 3D, 4, Figures S1, S3, S4). The two available Cossack Slav associated individuals (1,600–1,800 CE; UkrEM_Cossack_Slavs) have highest genetic affinity with the preceding Slav-related individuals as well as modern Ukrainians (Figures 3D, 4, Figures S1, S3, S4).

The UkrMA_Cuman individual is cladal with UkrMA_Post-Cuman_Cuman? in an f4-based test, but non-cladal with UkrMA_GoldenHorde_Nom, the former having relatively higher genetic affinity with some Siberian and East Asian groups (Tables S5–S6). What is more, UkrMA_WhiteCroat_Slavs is cladal with UkrMA_GoldenHorde_Slav/Nom? (Tables S5–S6) in the same test, as is, UkrEMA_Saltiv_Alans_2 with UkrMA_Cuman_2 (Tables S5–S6). Importantly, UkrMA_WhiteCroat_Slavs is cladal with preceding UkrIA_Chernyakhiv_1 and succeeding UkrEM_Cossack_Slavs (Tables S5–S6).

Using distal qpAdm modelling, UkrMA_Cuman, UkrMA_Post-Cuman_Cuman? and UkrMA_GoldenHorde_Nom can be explained with around half of AADR_Ukraine_Yamnaya, some AADR_Ukraine_Trypillia and some AADR_Mongolia_SlabGrave ancestry (35–62% + 18–39% + 14– 28% overall) (Figure 5A, Table S8). UkrMA_GoldenHorde_Slav/Nom? and UkrEM_Cossack_Slavs can be modelled as mostly AADR_Ukraine_Yamnaya, some AADR_Ukraine_Trypillia and a small amount of AADR_Mongolia_SlabGrave ancestry (53–62% + 37–45% + 1–2% on average) (Figure 5A, Table S8). For UkrMA_Cuman_2 and UkrMA_Nogai_1 we infer some AADR_Ukraine_Yamnaya and AADR_Ukraine_Trypillia, but around a half of AADR_Mongolia_SlabGrave ancestry (25–36% + 14– 22% + 48–56% overall), while UkrMA_Nogai_3 can be modelled with a small amount of AADR_Ukraine_Trypillia, some AADR_Ukraine_Yamnaya, but mostly AADR_Mongolia_SlabGrave ancestry (3–9% + 5–12% + 84–87%) (Figure 5A, Table S8).

Using proximal qpAdm modelling we found that UkrMA_Cuman_2 can be modelled as 100% UkrEMA_Saltiv_Alans_2, and UkrMA_Post-Cuman_Cuman? in turn as 100% UkrMA_Cuman_2 (Figure 5B, Table S11). UkrMA_GoldenHorde_Nom can be modelled as 91–99% UkrEIA_LateScythian_Cri_Nom and 1–9% AADR_Mongolia_SlabGrave ancestry, while UkrMA_Nogai_3 models as 37–48% UkrEMA_Saltiv_Alans_2 and 52–63% AADR_Mongolia_SlabGrave ancestry (Figure 5B, Table S11). UkrMA_WhiteCroat_Slavs and UkrMA_GoldenHorde_Slav/Nom? can be sourced from 100% UkrIA_Chernyakhiv_1, and UkrEM_Cossack_Slavs in turn from 100% from UkrMA_WhiteCroat_Slavs (Figure 5B, Table S11).

### Heterogeneity in Ukraine and elsewhere in Western Eurasia

As a means of estimating relative genetic heterogeneity, we analysed multidimensional Euclidean distances between individuals from PCA, grouped in the four date ranges discussed above: Late Bronze Age and pre-Scythian Early Iron Age (LBAEIA); Scythian period Early Iron Age (SEIA); post-Scythian Iron Age to Early Middle Ages (IAEMA); and Middle Ages and early modern period (MAEM) (Table S1). Considering the top principal components as continuous axes of covariation in ancestry, obtained through PCA following ^66^, we calculated Euclidean distance across the first 25 components to represent similarity or dissimilarity in genetic ancestry. This showed a general trend of high heterogeneity amongst the investigated individuals, with the exception of the SEIA date-group, for which it is somewhat decreased (Figure 6E). To contextualize these PCA-based heterogeneity estimates, we generated a baseline for Western Eurasia over the same time periods. Up to 100 randomly geographically distributed subgroups of individuals (within the same geographic distance of one another as the Ukraine study individuals in each date range) were also analysed in the same way. Plots of kernel density estimations for these show a shift towards higher Euclidean distances between the Ukraine genomes of this study when compared to the baseline in all date ranges. Notably, no matched random subsets from the Western Eurasia data returned higher average Euclidean distances than those obtained from the Ukraine genomes reported in this study (Figure 6A–D).

**Figure 6.**
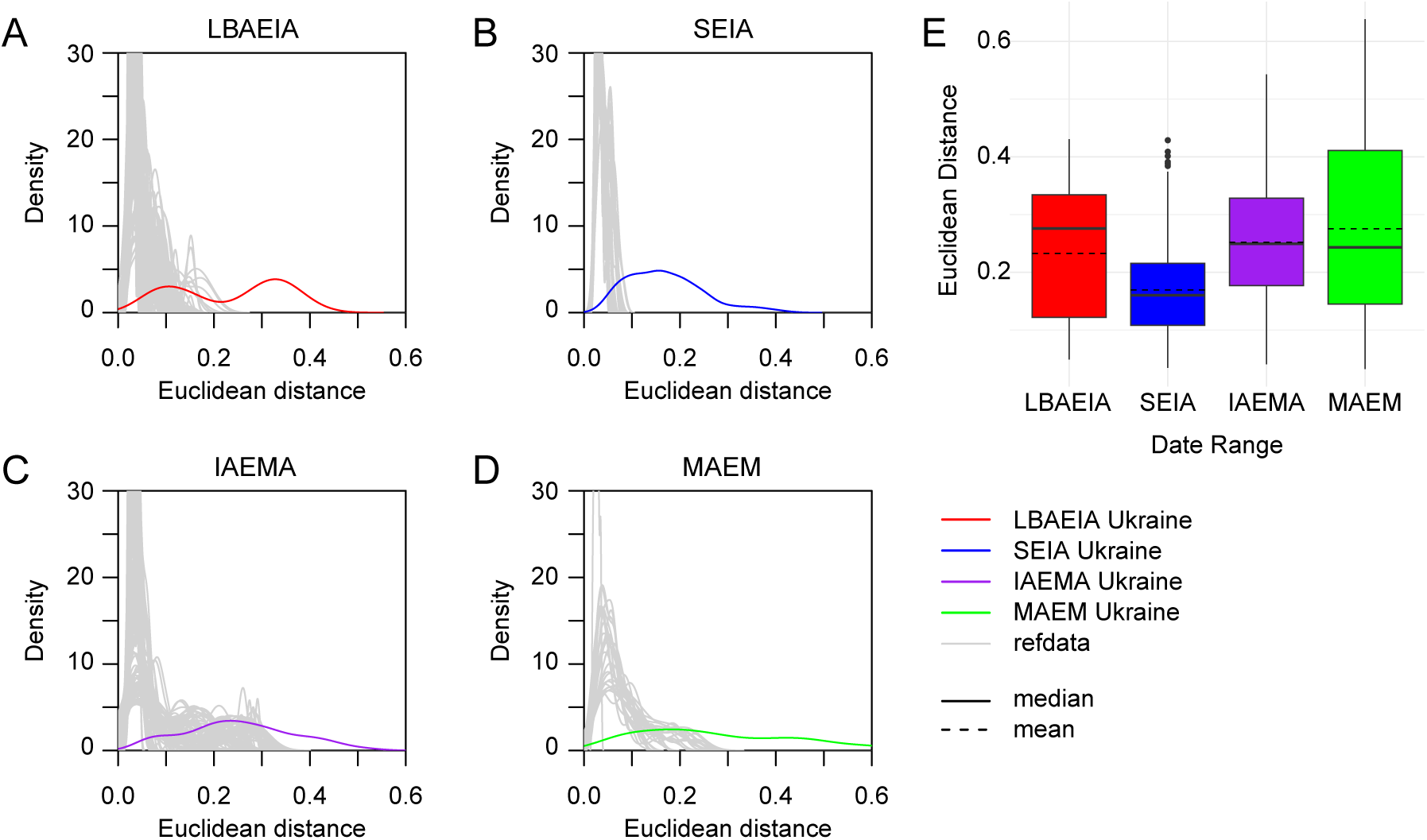
Investigating genetic heterogeneity in ancestry using PCA data. Kernel density estimations (KDE) of Euclidean distance sets for genomes of this study and corresponding comparative subsets (refdata), for (**A**) Late Bronze Age and Pre-Scythian Early Iron Age (LBAEIA, up to 105 distances used for KDE, 100 comparative subsamples), (**B**) Scythian period Early Iron Age (SEIA, up to 351 distances used for KDE, 63 comparative subsets), (**C**) post-Scythian Iron Age to Early Middle Ages (IAEMA, up to 325 distances used for KDE, 100 comparative subsets), (**D**) Middle Ages and Early Modern period (MAEM, up to 171 distances used for KDE, 28 comparative subsets). A cutoff at density 30 is used on the y-axis for better visibility of the Ukraine data. (**E**) Boxplot of Euclidean distances from the first 25 principal components of PCA analysis across date ranges (LBAEIA n=15, SEIA n=27, IAEMA n=26, MAEM n=19). Dashed lines indicate the mean. See also Table S1, Tables S3–S4.

## Discussion

The geographical location, landscape and ecotypes of the Ukraine region has made it a place of intersection and interaction between eastern and western neighbours, which has left its mark on the genetic composition of local populations. The genetic profiles of Ukrainian Mesolithic and Neolithic HGs are intermediate between Eastern and Western European HGs ^6,37,41^. The following early farmers (associated with Trypillia and Globular Amphora Culture) are similar to those from the rest of Europe ^6,36^. Peoples of the Bronze Age Yamna culture from the east were not only assimilated in the North Pontic region but were also the source of demic flow further into Europe ^1,2^.

In the post-Yamna period, a clear genetic differentiation of the BA/EIA populations is observed. Eastern and steppe affiliated groups are genetically similar either to Yamna people (BA Zrubna and FBA Bilozerska individuals) or individuals from the eastern side of the Western steppe (FBA/EIA Cimmerians). The genomes of western forest-steppe individuals are more similar to those of Central/Eastern (FBA/EIA Vysotska and Lusatian individuals) and Southeastern (Thracian Hallstatt) Europeans. Notably, we detect a signal of sex-biased admixture evident in the Zrubna individuals. Since the Zrubna culture is a product of migration from the eastern Pontic Caspian steppe into Europe similarly to Corded Ware Culture ^8,16^, the scenario of the migrating people being mostly men who then admixed with local women in Europe, which has previously been suggested for Corded Ware Culture ^53,56,67^, is also plausible for Zrubna.

During the Scythian period, we infer geographic structure that fits well with archaeology ^11^ and reflects their spread throughout the North Pontic steppe and forest-steppe. Most of the individuals associated with the Scythian culture from the western part of Ukraine (Illirian-Thracian basis) are genetically “local”, whereas genomes from the eastern part of Ukraine share more genetic ancestry with Western Steppe and East Asian populations. Individuals from the eastern part of Ukraine, archaeologically associated with the “local elite”, have more Southeastern European influences (Near Eastern ancestry) in their genome (including Y hg E1b), when compared to “local agriculturalists’’. Most of “nomad elite” individuals (Siversky Donets basin) have a genetic profile similar to that of people from the Caucasus. Patterns of genetic variation among social groups, as determined by the burial features and archaeological artifacts, appear more complex than those based on geography (Methods). Among the elite there are both individuals with a “local’’ genetic profile, and individuals with IA Western Steppe ancestry. There are intermediate profiles among local farmers and low-status nomads. Such patterns could be explained by the close incorporation of the Scythians into local society, and vice versa, including the interweaving of elites as part of the population admixture process.

It is natural to expect a sex-biased admixture in the case of nomadic invasion, and our data indirectly support such assumptions: there are 8 males and 7 females among Scythian groups with a local archaeological background (right and left bank of Dnipro), but 11 males and only 3 females among groups with a nomad archaeological background (Siversky Donets, Northern Black Sea) (Table S1). Scythian family burials in kurgans on both the right and the left banks of Dnipro, and kinship groups for two generations established for 8 samples, indicate sedentary lifestyle, at least among a part of those with non-nomadic roots. The low mobility of Scythians in the North Pontic forest-steppe has previously been inferred using Sr isotope analysis ^68^.

IA Chernyakhiv individuals from central and eastern forest-steppe of Ukraine form two genetic subgroups, one with a more Northern/Central and the other with a Southern European genetic profile, reflecting the polyethnicity of the Chernyakhiv group visible even within one location. The genetically more northern individuals may be potentially associated with the relocation of Goths into the North Pontic area. At the same time, the unusual Near Eastern maternal ancestry of the Chernyakhiv_3 individual from the Eastern Carpathians might help to explain previous archaeological observations. The Komariv-1 settlement, where this young woman was from, had the only known glass production point outside the Roman Empire ^69,70^; ancient glass production was concentrated primarily in the Middle East ^71^. The unusual Near Eastern ancestry might originate from eastern Mediterranean craft bearers living in the settlement.

The genetic composition of the EMA Alans and Bulgars examined in this study is similar to individuals from the Caucasus or Central Asia. The latter component indicates a constant influx of new migrants from Central Asia, which is consistent with previous archaeological claims ^72^. Our data suggest that the Central Asia affiliated groups lived in the region permanently, consisting of both men and women whose genetic profiles, including autosomal and mitochondrial data, show no evidence of mixing with the local people. The high degree of genetic similarity among individuals thought to be Alans and Bulgars based on their burial ritual (catacombs *vs* pit burials, respectively) and other features, is consistent with them coming from a single, genetically homogenous population. It is unlikely that funeral rites provide a robust means of distinguishing Alans from Bulgars. This idea is consistent with other archaeological interpretations, such as not only catacombs but also pit burials of the Saltiv culture in the forest-steppe of Ukraine being associated with Alans, who came from the Caucasus ^73^.

The Cuman, Golden Horde and Post-Cuman individuals of this study are genetically most similar to Western Steppe peoples. However, Nogai, the last nomadic group in the North Pontic, which is thought to have included remnants of Khazars, Pechenegs, Cumans as well as Mongol peoples, have genetic profiles indicating high levels of East Asian ancestry. The Nogai nomadic migrants with East Asian ancestry, similarly to Alans described above, include both males and females without genomic signals of admixture with autochthonous people. The genetic composition of contemporary medieval and early modern Slavs is similar to modern Ukrainians. Moreover, such genetic profiles are traceable among some individuals from previous periods since LBA and, apparently, are of local Eastern European ancestry.

To conclude, DNA analyses of samples taken from archaeological sites representing different regions of Ukraine in the chronological interval from around 9,000 BCE to 1,800 CE show that the ancient population had a diverse range of ancestries as a result of frequent movements, assimilation and contacts. From the Mesolithic until the time of Vysotska and Bilozerska cultures at the end of the Bronze Age, broad-scale ancestry proportions are similar to contemporary populations in the rest of Europe – first HGs, then early farmers and finally a mixture between early farmers and Steppe pastoralists. Starting from the Cimmerian time (EIA) until the Middle Ages, the appearance of eastern nomads in the Pontic region became a regular occurance. Their genetic composition varied from Yamna-like superimposed on the locals, as with Scythians and Cumans, to high degrees of East Asian ancestry and minimal local admixture, as with Alans-Bulgars and Nogai. During that time, nomadic populations were recorded in the steppe zone, whereas individuals from the rest of the Ukrainian region had mostly European ancestry, associated with local predecessors, as well as Thracians, Greeks, Goths etc.

The palimpsest of migration and population mixing in the Ukraine region will have contributed to the high genetic heterogeneity in geographically, culturally and socially homogenous groups, with different genetic profiles present at the same site, at the same time and among individuals with the same archaeological association. Nevertheless, the broad-scale “local” genetic profile, which is similar to modern Ukrainians, persists in the region through time. This ancestry composition can be traced back to the Zrubna individuals at least, and is seen among Vysotska and Lusatian individuals, Scythians from the west and contemporary agriculturalists from the east, among the Chernyakhiv population, as well as medieval and early modern Slavs. Despite clear signatures of high migration activity, including from East Asia, as well as extensive admixture, we infer a major autochthonous component to Ukrainian ancestry, at least since the Bronze Age.

## Acknowledgments

This project has received funding from the European Union’s Horizon 2020 research and innovation programme under the Marie Skłodowska-Curie grant agreement No 101029303. LS was further supported by Estonian Research Foundation grant PRG1027. OU was supported by the European Union’s Horizon 2020 research and innovation programme under the MSCA4Ukraine grant agreement No 1232832. IS was supported by the research program of the German Archaeological Institute ‘Documenting, Recording and Saving Ukrainian Archaeological Heritage’ (2023). RM was supported by an SSHRC doctoral studentship grant (G101449: ‘Individual Life Histories in Long-Term Cultural Change’). PS was supported by the European Molecular Biology Organisation, the Vallee Foundation, the European Research Council (grant no. 852558), the Wellcome Trust (217223/Z/19/Z), and Francis Crick Institute core funding (FC001595) from Cancer Research UK, the UK Medical Research Council, and the Wellcome Trust. MGT was supported by ERC Horizon 2020 research and innovation programme grant agreements: no. 951385 (COREX) awarded to MGT; no. 865515 (SUSTAIN) awarded to Maria Ivanova-Bieg; no. 324202 (NeoMilk) awarded to Richard Evershed; no. 788616 (YMPACT) awarded to Volker Heyd; and by Wellcome Senior Research Fellowship Grant 100719/Z/12/Z awarded to MGT. Data was processed at the Advanced Sequencing Facility and Scientific Computing at the Francis Crick Institute. Analyses were carried out using the facilities of the UCL Department of Computer Science High Performance Computing Cluster.

## Author contributions

Conceptualization: LS; funding acquisition: LS, PS, MGT; resources: OU, SZ, IS, KG, MB, DP, IB, DG, VO, GT, SA, VR, YB, VK, OS, OP, BM, SD, AH, RH, SS, VA, OL, ST, VS, YMS, PS, MGT; methodology: LS, RM; investigation: LS, MJ, CLS, KA, MK, MW, MS; formal analysis: LS, OU, CB, AG, RM; data curation: LS, OU; visualization: LS, RM; project administration: LS; supervision: PS, MGT; writing—original draft: LS, OU, SZ, IS; writing—review & editing: LS, OU, SZ, IS, KG, MB, DP, IB, DG, VO, GT, SA, VR, YB, VK, OS, OP, BM, SD, AH, RH, SS, VA, OL, ST, VS, YMS, MJ, CLS, KA, MK, MW, MS, CB, AG, RM, PS, MGT.

## Declaration of interests

The authors declare no competing interests.

## Supplemental information

Document S1. Figures S1–S7.

Table S1. Information about the individuals of this study. Related to Figure 2, Table 1.

Table S2. Y chromosome informative positions for haplogroup determination. Related to Table 1.

Tables S3-S4. Modern and ancient comparison populations used in different analyses. Related to Figures 3–5.

Tables S5-S6. f4 statistics’ results. Related to Figure 3.

Tables S7-S11. qpAdm modelling results. Related to Figure 5. Tables S12-S13. f3 statistics’ results. Related to Figure 3.

## Methods

### RESOURCE AVAILABILITY

#### Lead contact

Further information and requests for resources and reagents should be directed to and will be fulfilled by the lead contact, Lehti Saag.

#### Materials availability

This study did not generate new unique reagents.

#### Data and code availability

Shotgun sequencing data mapped to the human genome have been deposited at European Nucleotide Archive and are publicly available as of the date of publication. Accession numbers are listed in the key resources Table.

All original code has been deposited at Github and is publicly available as of the date of publication. DOIs are listed in the key resources Table.

Any additional information required to reanalyse the data reported in this paper is available from the lead contact upon request.

### EXPERIMENTAL MODEL AND STUDY PARTICIPANT DETAILS

The teeth used for DNA extraction were obtained with relevant institutional permissions from various collections in Ukraine: Museum of Archaeology of V. N. Karazin Kharkiv National University, Zaporizhzhia National University, I. Krypyakevich Institute of Ukrainian Studies of the National Academy of Sciences of Ukraine, Institute of Archaeology of National Academy of Sciences of Ukraine, Odesa Archaeological Museum, M. F. Sumtsov Kharkiv Historical Museum, Laboratory of Archeology and Ethnology of V. O. Sukhomlynskyi Mykolaiv National University, Mykolaiv Regional Museum of Local History, Center for the preservation and preservation of monuments of archaeology of the Poltava region for the sake of Poltava, Archaeological Research Laboratory of National Pedagogical Dragomanov University, Lviv Historical Museum.

DNA was extracted from 129 tooth or bone samples from ancient individuals from the territory of present-day Ukraine – 2 from the Neolithic (UkrN; 7,000–6,000 BCE), 14 from the Bronze Age to the beginning of the Iron Age (UkrBA, UkrFBA/EIA; 3,000–700 BCE), 7 from the beginning of the Early Iron Age (UkrEIA; 900–700 BCE), 46 from the Scythian period of the Early Iron Age (UkrEIA; 700–300 BCE), 8 from the end of the Early Iron Age (UkrEIA; 400–1 BCE), 16 from the later Iron Age (UkrIA; 1–400 CE), 12 from the Early Middle Ages (UkrEMA; 800–900 CE) and 24 from the Middle Ages to the early modern period (UkrMA, UkrEM; 900–1,800 CE) (Figure 2, Table S1). More detailed information about the archaeological periods and the specific sites and burials of this study is given in Methods.

### NEOLITHIC

The Neolithization of the territory of present-day Ukraine occurred during 7,000–5,000 BCE and spread from the Balkans through the Danube area via several migration waves, involving tribes of the Linear Pottery, Cucuteni-Trypillia and some other European cultures. Neolithic archaeological cultures found on the territory of Ukraine are Bug-Dniester culture, Dnipro-Donets culture, Azov-Dnipro culture, Pit-Comb Ware culture, etc. ^76^

Azov-Dnipro (Mariupol) culture *(Азово-Дніпровська культура)*

The landmarks of the Sur and Azov-Dnipro cultures are known from the Lower Dnipro region. The Azov-Dnipro culture landmarks are dated from the 6th to the beginning of the 5th millennium BCE. They are represented by both settlements (Semenivka, Stone Grave) and burial grounds (Lysa-Gora, Mamay-Gora, etc.). ^77^

**Mamay-Gora (Mamai-Hora)** (Мамай-Гора) (Archaeology, text – G. Toshev, S. Andrukh)

The Mamay-Gora hill is a multi-layered archaeological site located near the village Velyka Znamianka in the Vasilyevsky district (former Kamensko-Dneprovsky) of the Zaporizhzhia region, on the high left bank of the Kakhovka reservoir. This elevated location was used by steppe tribes as a burial place from the Neolithic to the 15^th^ c. CE. Now the burial ground occupies approximately 30 hectares. Its core is formed by 5 kurgans (three elongated and two round), stretching along the west-east line. ^78–82^

Since 1988, Mamay-Gora has been studied by archaeological expeditions of the Zaporizhzhia National University. During more than 34 years over 900 burial complexes (kurgans and ground burials) have been found there. The oldest are two Neolithic burial grounds, followed by the complexes of the Eneolithic, Bronze Age, and Scythian time (n=398). Later, the Sarmatians (n=2), the tribes of the post Golden Horde time (Cumans (n=60), Nogai (n=200)) were buried there. Currently the excavations are interrupted because of the Russian aggression in Ukraine.

The Neolithic burial ground under research is a family burial belonging to the first period of the Azov-Dnipro culture. It comprises 26 graves forming 2 rows stretched perpendicular to the Dnipro riverbed. Burials are in individual pits at different distances from each other. The skeletal remains lay stretched out on backs, with heads facing south or west. In most of the burials, artefacts were found: pendants made of red deer’s teeth, necklaces made of shells or stones, flint plates. In 13 burials, ochre was found on the bones, on the bottom of the pit or in the filling. Among the buried individuals there were men (n=6), women (n=4), unidentified adults (n=9) and children (n=7). ^83^

**Location:** 47.432845 N, 34.27259 E. Mamay-Gora tract, Velyka Znamianka village, Vasylivsky District, Zaporizhzhia Region.

**Excavations:** 1999.

**Excavation authors:** Svitlana I. Andrukh, Gennadi M. Toshev, Zaporizhzhia National University, Zaporizhzhia, Ukraine.

**Storage of anthropological materials:** Zaporizhzhia National University, Zaporizhzhia, Ukraine.

**Description of samples:**

One sample was taken for aDNA analysis.

**UKR008**. *Mamay-Gora, Neolithic burial 26.* Excavated in 1999. Human remains were found at a depth of 1.17 m. The contours of the pit were not traced. The individual was stretched out on the back with hands along the body. The head was oriented to the southeast/east. Chronology according to archaeology: 7,000–6,000 BCE. ^81^

### BRONZE AGE

#### Zrubna (Srubnaya)/Timber-Grave cultural and historical community (Зрубна культура) (Archaeology – V. Mikheev; text – I. Shramko, S. Zadnikov)

The Zrubna cultural and historical community was formed in the Late Bronze Age (1,800–1,100 BCE) and spread in the steppe and forest-steppe zones of Eastern Europe from the Dnipro river to the Ural mountains. The main variations are Berezhniv-Mayivska and Pokrovska Zrubna cultures. The Pokrovskа Zrubna culture (1,800–1,400 BCE) was widespread in the steppe and forest-steppe zones from Siversky Donets river to the Volga river. Separate sights are presented in the Urals. The Berezhniv-Mayivska Zrubna culture (1,700–1,100 BCE) was widespread in the steppe and forest-steppe area from Ingulets river to the Volga river. The main types of archaeological sites are settlements, kurgans and ground burials. The economy was based on cattle breeding and agriculture. Bronze casting production and bone processing were also developed.

#### **Sukha Gomilsha** *(Суха Гомільша)*

The kurgan is located near the centre of the northern part of the Sukha Gomilsha hillfort belonging to the Saltiv culture (8th to 10th c. CE). It stands out in the area like a small hill of 0.2–0.4 m in height and 21–22 m in diameter.

**Location:** 49.547122 N, 36.362542 E. Sukha Gomilsha village, Slobozhanska territorial community, Chuhuiv district, Kharkiv region.

**Excavations:** 1982.

**Excavation authors:** Volodymyr K. Mikheev, V. N. Karazin Kharkiv National University, Kharkiv, Ukraine.

**Storage of anthropological materials:** Collection of the Museum of the Archaeology, V. N. Karazin Kharkiv National University, Kharkiv, Ukraine.

**Description of samples:**

One sample was taken for aDNA analysis.

**UKR055.** *Sukha Gomilsha, burial 3.* Excavated in 1982. Inlet burial in kurgan. A human skeleton of poor preservation was found at a depth of 1.0 m from the modern surface level. The skeleton lay on the back, arms stretched along the body. The head was oriented to the west. There was a vessel near the head. The burial belonged to the Late Bronze Age and dated to 1,300–800 BCE according to archaeology. ^84,85^ Chronology according to ^14^C dating: 1873–1566 cal BCE (3404±35 BP).

### FINAL BRONZE AGE

#### **Bilozerska culture** (Білозeрська культура)

*(Archaeology, text – K. Gorbenko)*

The Bilozerska culture is an archaeological culture of the Final Bronze Age (1,200–900 BCE). It was defined as an archaeological culture by V.V. Otroshchenko, I.T. Chernyakov and V.P. Vanchugov in the 1980s. The name comes from the settlement on the shore of the Bilozersky estuary (now the Kakhovka Reservoir) in the city of Kamianka-Dniprovsk. The culture is widespread in the steppe zone of Ukraine and Moldova. A few sites were found on the Lower Don, on the territory of the Kuban and in Crimea.

The sites of the Bilozerska culture are represented by settlements alongside rivers and estuaries (Tudorovka, Voronivka, Zmiivka), kurgans (Shyroka Mohyla, Stepovy, Kochkovate) and ground burials (Brylivka, Shiroke, Budurzhel), complexes of foundry molds (Zavadivka, Novooleksandrivka), foundry workshops (Kardashinka), treasures of metal objects (Novogrigoryivka) and others. Presumably, the carriers of the culture tried to control the supply routes for raw materials, prestige products, and possibly grain.

In the Prut-Dniester interfluve, the Bilozerska culture has three periods (S. Agulnikov): early (end of the 13th to first half of the 12th c. BCE); middle (end of the 12th to first half of the 11th c. BCE); late (second half of the 11th to first half of the 10th c. BCE). Represented mainly by settlements, kurgans, ground burials, hillfort (Dyky Sad).

Approximately 165 settlements of Bilozerska culture are known. The maximum area of these reached 3.0–4.5 hectares. The layout of the settlements was street-type. The size of houses ranged from 15 to 90 m^2^.

Burials are connected with settlements and are best studied in the places where the settlements are clustered. So far, more than 800 burials have been identified. The population of the Bilozerska culture used a burial tradition that was different from the prevailing one in the steppe. Old graves were almost not used for burial needs. People built new graves or used ground cemeteries (kurgan, ground grave).

The population was heterogeneous. Anthropologists note the similarity of Bilozerska people with narrow-faced skulls from ground cemeteries belonging to the Noua culture. A common anthropological dynamic is traced for Bilozerska culture and Ukrainian Zrubna culture. In general, the local anthropological basis of Bilozerska culture seems clear, although external influences took place all over the Bilozerska culture world. ^86–93^

#### Dykyi Sad fortified settlement *(Городище Дикий сад)*

Dykyi Sad (Wild Garden) fortified settlement is located on a plateau at the confluence of the Southern Bug and Ingul rivers, in the historical centre of the modern city of Mykolaiv. It was built in the form of an oval elongated along the SE-NW axis. The total area of the preserved territory reaches more than 5 hectares. The settlement belongs to the Bilozerska archaeological culture (Bilozerska-Tudorovska community) of the Final Bronze Age (13/12–12/11 c. BCE). According to ^14^C dating, the time ranges within 1,186–925 BCE.

Residential, economic, defence, ritual and cult objects have been found on the territory of the settlement. It can be argued that during the heyday of the settlement, a clear system of uniform planning and construction was formed within the territory: a “citadel” surrounded by a moat, a “suburb” in the hemisphere of the outer moat, a “market”. The plan of the settlement corresponds to the classic concept of “urbs” (“city”).

In total, 53 archaeological sites in an area of more than 8,000 m^2^ have been investigated. Among them are 41 premises with yards, 3 utility pits outside the “suburb”, a moat around the “citadel” (with buried skulls), a moat around the entire settlement, remains of defensive structures along the moat of the citadel, ritual and cult ramp, the central site of the “citadel”, the economic and ritual site opposite the moat of the “citadel”, the central square of the remote “suburb”, 21 pits for economic and ritual purposes, a pit behind the outer moat. The pits were located on a flat area between the houses, forming a kind of central economic area of the “suburb”. Remains of ceramic dishes, burials of human skulls, animal and fish bones, remains of charred grains of common millet, barley, wheat, cultivated grapes and charcoal were found in the pits. ^90,91,94,95^

**Location:** 46.980272N, 31.983500E. Mykolaiv city (Naberezhna str.), Mykolaiv region.

**Excavations:** 2004, 2007.

**Excavation authors:** Kyrylo V. Gorbenko, Petro Mohyla Black Sea National University, Mykolaiv, Ukraine.

**Storage of anthropological materials:** Mykolaiv Regional Museum of Local History, Mykolaiv, Ukraine.

**Description of samples:**

Two samples were taken for aDNA analysis.

**UKR149.** *Dykyi Sad, the moat of the hillfort “citadel”. Excavation 13, lower layer.* The arc-shaped moat encloses the “citadel” of the settlement, stretching along the southeast-northwest axis. The length of the studied part is 130 m (the total length is approximately 140–150 m), width 5.0 m, depth 3.0 m. In the southern part of the moat, the stone foundation of a bridge (large limestone slabs) is located. The foundation is rectangular in plan, stretched across the moat along the north-south line with a slight deviation to the northeast. The dimensions are 3.64×2.0 m, and the width of the central part is 1.4 m due to the shifting of the stones. At the distance of 5 m to the west of the foundation, in the middle of the stone lining and among ceramic fragments, a human (female) skull was found. Chronology according to archaeology: 13–12 с. ВСЕ.

**UKR150**. *Dykyi Sad, the central site of the distant “suburb” of the settlement. Pit № 8.* The ritual pit had a rounded shape with an extension to the lower part. The upper and lower diameters were 0.5 m and 1.05 m, depth was 0.65 m. The pit was filled with grey humus loam (burnt soil), small and medium-sized rubble. At a depth of 0.25 m there was a stone backfilling (143 stones). Pot remains were found in the eastern part of the pit at a depth of 0.5 m. In the southern part at a depth of 0.45 m there was a human skull without a lower jaw, facing west. The skull was covered with small stones. The bottom of the pit was flat and clayey. Chronology according to archaeology: 12 с. ВСЕ.

### FINAL BRONZE AGE / EARLY IRON AGE

#### **<u>Vysotska culture</u>** (Висоцька культура)

*(Archaeology, text – M. Bandrivskyi)*

The Vysotska culture was one of the bright archaeological cultures of the end of the Bronze Age and the beginning of the Iron Age in Central-Eastern Europe. It occupied a small area in the west of Ukraine: from the upper course of the Zbruch river in the east to the upper course of the Western Bug in the west, and from the Volyn upland in the north to the middle course of the Seret and Strypa rivers in the south. Graveyards and individual burials, including kurgans, are the source of knowledge about this culture since settlements are almost unknown today.

The culture’s chronology includes three periods: early – from around 1250–1150 BCE (Bronze Age to Hallstatt period A1 (HaA1)); middle – 1150–950/920 BCE (HaA2–HaB1); late – 950/920–around 725 BCE (HaB3–HaC1).

As a rule, the burials were oriented to the south, graves were placed in rows, grave pits were almost completely absent (the remains were buried in the lower layer of the modern black soil, in recesses up to 15–20 cm, which resembled the body contours), many of the burials were likely covered by an earthen embankment that has not been preserved. The skeletons were on their backs in a straightened position, their hands mostly folded on the chests. Single burials are the most numerous, but also double burials are found with a man and a woman buried at the same time. In the late period of the culture, burials in stone tombs (cists) appeared.

The artefacts consisted of moulded dishes, which were placed around the head or less often near the feet or on the side, as well as jewellery and toiletries. For the late period, stone war hammers are known. Although the Vysotska culture is usually attributed to the Central European Urnfield culture, it differs by a strong military focus. The carriers of the culture had well-established trade relations with distant western centres of metalworking. ^23,96^

#### **Petrykiv cemetery** (Петриківський могильник)

The Petrykiv cemetery was found in 1987 by O. Sytnyk in Petrykiv village, which is a suburban area of Ternopil city. In 1995 the excavations were started. The cemetery is located in the eastern part of the Ternopil Plateau of Western Podillia, on the flat terrain of the right bank of the Seret river – the left tributary of the Dniester river – on the southeastern slopes. Fragments of ceramics, sometimes metal products, flint production waste are found throughout the territory. A total of 148 burials were found in the cemetery. The inhumation, cremation and reburial of individual bones and cenotaphs is described.^97^

**Location:** 49.534488 N, 25.579807 E. Petrykiv village, Ternopil district, Ternopil region, Ukraine.

**Excavations:** 1995–1996.

**Excavation authors:** Mykola Bandrivskyi, I. Krypyakevich Institute of Ukrainian Studies of the National Academy of Sciences of Ukraine, Lviv, Ukraine

**Storage of anthropological materials:** I. Krypyakevich Institute of Ukrainian Studies of the National Academy of Sciences of Ukraine, Lviv, Ukraine

**Description of samples:**

Three samples were taken for aDNA analysis.

**UKR170.** *Petrykiv, burial 35.* Excavated in 1995. The burial was found at a depth of 0.35–0.45 m. The size of the grave pit was 2.10 x 0.5 m. The skeleton was straightened with the head oriented to the south. The length of the skeleton was 1.77 m. The skull lay on the left temporal and facial bones. The cranial sutures were separated. Half of the lower jaw was lost. From the preserved molars and premolars, it can be judged that the individual was 18–20 years old at the time of death. The arms were folded on the chest. The legs were straightened, ankles lying next to each other, but the phalanges were absent. On the left side of the pelvic bones at a depth of 0.25 m lay a large fragment of a pot. Chronology according to archaeology: 13–9 c. BCЕ.

**UKR171.** *Petrykiv, burial 58.* Double burial. Excavated in 1996. The grave at a depth of 0.40 m contained remains of two skeletons, with their heads oriented to the south. Almost all bones were displaced from their original positions. Based on the placement, size, and other features of the preserved bones, we can assume that the main burial (UKR171) belonged to a man with very large bones, who laid on his back with straight legs. The skull was badly destroyed, the lower jaw was pushed 20 cm to the side. Only fragments of ribs and other bones were also preserved. On the left side, close to the man’s skeleton lay another, but much smaller and with a more delicate and feminine bone structure. Only the lower limbs including the ankle-foot joints, a part of the skull, the lower jaw and the bones of the forearms were preserved. Bones had intensely coloured bright green stains from bronze ornaments. At a depth of 0.40 m near the place where the frontal parts of the man’s and woman’s skulls touched, a small bronze ring was found. No other artefacts were found, except for small and probably redeposited fragments of pottery. Chronology according to archaeology: 13–9 c. BCЕ. Chronology according to ^14^C dating: 1,278–1,055 cal BCE (2967±30BP).

**UKR174.** *Petrykiv, burial 79.* Excavated in 1996. The burial was found at a depth of 0.3–0.5 m. It was accompanied by six vessels and an animal skull. Only the lower part of the legs and the skull were preserved. Considering the position of the bones, the head was initially oriented to the south but later the skull was moved lower to the right knee joint. The lower jaw was absent. The placement of the foot bones proves that they both faced east. It is also interesting that a large fragment of a pot lay between the heel bones of the ankle joints. A tulip-shaped pot, a bowl and two ladles were found in the burial. Chronology according to archaeology – 13–9 c. BCЕ. Chronology according to ^14^C dating: 359–104 cal BCE (2169±29BP). Based on the ^14^C date and the genetic profile difference, this individual was separated from other Vysotska culture individuals in analyses.

#### **Syncretic Ulvivok-Rovantsi cultural group (Lusatian culture, Vysotska culture)** Ульвівецько-Рованцівська культурна група (Лужицька культура, Висоцька культура) (Archaeology, text – D. Pavliv)

Syncretic Ulvivok-Rovantsi cultural group (Lusatian culture, Vysotska culture) formed on the border of the 2^nd^ and 1^st^ millennium BCE (from 10th c. to the first part of 7th c. ВСE) in the southwest of Volyn, in the contact zone of the Vysotska and Lusatian cultures.

Presumably, the population is related to the carriers of the Vysotska culture in western Ukraine and the Lusatian culture on territories west of the Bug up to Silesia. Certain features of the culture point to continuity with the kurgan cultures of the Middle Bronze Age and the Urnfield culture of Central Europe of the end of the Bronze Age, in particular the Urnfield culture of the Northern Alpine and Middle Danube regions.

Ulvivok-Rovantsi landmarks are represented by ritual necropolises on the high banks of the rivers Styr, Chornoguzka, Western Bug and Solokia. They have peculiar features of the funeral rite – combination of different types of inhumations and cremation, separate burials of skulls – and contain original complexes of ceramics, including special forms of funeral vessels. At the late stage the Ulvivok-Rovantsi culture was strongly influenced by the Lusatian culture. ^98–104^

#### **Rovantsi cemetery** (Рованцівський могильник)

The Rovantsi cemetery is located on the border of the Volyn Upland and Polissia, on the high left bank of the Styr River – the right tributary of the Pripyat River – near Lutsk, between the villages of Rovantsi and Boratyn. According to radiocarbon dating, the necropolis belongs to the second part of 9th to 8^th^c. BCE (Kyiv Radiocarbon Laboratory of the Institute of Geochemistry of the Science of the National Academy of Sciences of Ukraine, laboratory number Кі–9815). It is the easternmost landmark and the largest burial ground of the Ulvivok-Rovantsi group. The total area is approximately 2,000 square meters. The site was found by V. Shkoropad in 1986 and was studied in 1987, 1989–1990 by expedition led by Dmytro Pavliv.

The necropolis belongs to the ground type with a bi-ritual burial rite with a significant predominance of inhumation. In total, 80 inhumation burials arranged in rows stretched from east to west with heads oriented to the south, 12 cremation burials of three types, 16 separate burials of skulls, as well as pottery between the burials, were investigated at a depth of 0.4–0.9 m. Burials were accompanied by moulded dishes of various shapes: pots, bowls, mugs, jugs, special funerary ceramics. The pottery was decorated with complex ornaments. Some of the burials contained bronze items: temple pendants, diadems, breast ornaments, rings, wrist and ankle bracelets. ^105,106^

**Location:** 50.725306N, 25.361439E. Boratyn and Rovantsi villages, Lutsk district, Volyn region.

**Excavations:** 1987, 1989–1990.

**Excavation author:** Dmytro Pavliv, Ivan Krypiakevych Institute of Ukrainian Studies of National Academy of Sciences of Ukraine, Lviv, Ukraine

**Storage of anthropological materials:** Institute of Archaeology, National Academy of Sciences of Ukraine, Kyiv, Ukraine.

**Description of samples:**

Three samples were taken for aDNA analysis.

**UKR167.** *Rovantsi, burial 9.* The southern part of the cemetery. The depth was 0.7–0.8 m. There was a skeleton that was osteologically estimated to be a man (but genetically female) aged 30–40 years. The body was straightened, the head oriented to the southwest. The skull was lying on the occipital bone, tilted to the left side. The arms were bent at the elbows and placed on the chest and the stomach. There were no artefacts. Chronology according to archaeology: 9th to 8th c. BCЕ.

**UKR168.** *Rovantsi, burial 44.* The south-eastern part of the cemetery. The depth was 0.6–0.7 m. The body was straightened, the head oriented to the south. The skull rested on the right temporal bone. The arms were bent at the elbows, the forearm of the right hand was on the stomach, the left hand was raised to the skull. There were no artefacts. Chronology according to archaeology: 9th to 8th c. BCЕ.

**UKR169.** *Rovantsi, burial 69.* The southern part of the cemetery. The depth was 0.6–0.7 m. The body was straightened, the head oriented to the south. The skull was lying on the occipital bone. The arms were bent at the elbows, the forearm of the right hand was on the chest, the left hand was on the stomach. There were no artefacts. Chronology according to archaeology: 9th to 8th c. BCЕ.

### EARLY IRON AGE

#### **<u>Cimmerian culture</u>** (Кіммерійська культура)

*(Archaeology – I. Shramko; text – I. Shramko, S. Zadnikov)*

The Cimmerian culture (10th–8th c. BCE) landmarks in Ukraine are represented exclusively by inlet burials in Bronze Age kurgans. Not many artefacts have been found: mostly weapons, parts of a horse’s bridle, work tools, pottery. The Cimmerians are the first people in the Northern Black Sea region known from written sources. They were nomads, armed horsemen who fought on the territory of Asia Minor, mentioned in cuneiform texts.

#### **Kumy** (Куми)

The kurgan near the village of Kumy was discovered in 2010 during an archaeological survey ^107^. The site with kurgans is located in the fields of the Krasnograd Research Station, near the village of Kumy, Krasnograd District, Kharkiv Region. The kurgan, located on the edge of the watershed plateau between the right tributaries of the Berestova river (right tributary of the Orel river), has nine mounds of various sizes, most of which have been ploughed and are barely visible on the surface of the field. ^108,109^

**Location:** 49.3246500 N, 35.3687167 E. Kumy village, Krasnograd district, Kharkiv region, Ukraine.

**Excavations:** 2010.

**Excavation authors:** Iryna Shramko, Museum of Archaeology of V.N. Karazin Kharkiv National University.

**Storage of anthropological materials:** Museum of Archaeology of V.N. Karazin Kharkiv National University.

**Description of samples:**

One sample from the Early Iron Age was taken for aDNA analysis.

**UKR066.** *Kumy, kurgan 6, burial 5.* The inlet burial of a Cimmerian nomad. The burial was found at a depth of 0.7–0.9 m above the level of the reference point. The contours of the pit were not traced. The skeleton of an adult man (35–36 years old) was lying on his right side, with his head to the west. The bones of the right hand were slightly bent at the elbow and were stretched along the body. The left hand was missing. There were no artefacts. Chronology according to archaeology: 10th–9th с. ВСE. Chronology according to ^14^C dating: 1195–919 cal BCE (2865±39BP).

#### Thracian Hallstatt (Thraco-Cimmerian) culture *(Фракійський Гальштат)*

*(Archaeology, text – I. Break)*

The Thraco-Cimmerian culture still does not have the status of a distinct archaeological culture in historiography. It was described in 1920–1930 based on horse ammunition items from hoards belonging to the late Urnfield and early Hallstatt period in Central and Eastern Europe. The Kartal III burial ground is the only complete monument of Thracian-Cimmerian culture to date. ^110^

#### **Kartal** (Картал)

The Kartal settlement is a multi-layered archaeological site, located on the left bank of the Lower Danube, 1.5 km east of the village Orlovka. For this study, the materials from the burial ground of the third cultural horizon (Kartal III) were used. This horizon dates to the Middle Hallstatt period around 9th to 8th c. BCE. A total of 530 graves were excavated (about half of the entire cemetery). The features of the artefacts and the burial rite indicate that Kartal III belongs to the Thracian-Cimmerian culture. The population of the settlement consisted of two main ethnocultural components – Thracian and Iranian. Possibly, they represented the tribal world of the Western (South-Western) Balkans (Illyrians, Paeonians, Dardanians), as well as the indigenous people as a relic of the Indo-Aryan Late Bronze Age community (Zrubna-Sabatynivska and Bilozerska cultures). ^111^

**Location:** 45.319997 N 28.411989 E. Orlivka village, Izmail district, Odesa region.

**Excavations:** 2005, 2007, 2008.

**Excavation authors:** Igor Bruyako, South Ukrainian K. D. Ushinsky National Pedagogical University State Institution, Odesa.

**Storage of anthropological materials:** Odesa Archaeological Museum.

**Description of samples:**

Seven samples were taken for aDNA analysis, DNA was successfully extracted from six.

**UKR000.** *Kartal, burial 132.* The burial pit was not traced. The skull stood at a depth of 60 cm. The skeleton lay in a crouched position on its right side, with the head oriented to the south. The arms were bent at the elbows. Behind the head there was a lower jaw of small cattle. Opposite the front part of the skull there was a small black-clay jug. Chronology according to archaeology: 9th to 8th c. BCE. Chronology according to ^14^C dating: 900–798 cal BCE (2676±30 BP).

**UKR001.** *Kartal, burial 124.* The shape of the pit was close to rectangular. The depth of the contours of the pit was 75 cm, the size 210×110×15 cm. The skeleton was fragmented, the position crouched, on the right side, head oriented to the south. In front of the skull there was a black-clay cup-shaped vessel. Chronology according to archaeology: 9th to 8th c. BCE.

**UKR002.** *Kartal, burial 19.* The rectangular pit with strongly rounded corners was oriented along the NNW-SSE line. The contour of the pit was at the level of 80 cm. The length of the pit was 215–217 cm, the width in the middle 110 cm, the depth 25–30 cm. The skeleton lay in a crouched position on the right side, head oriented to SSE. The arm bones were bent at the elbows, the right hand was brought under the skull. There was a black polished goblet near the left hand, with its neck facing the front of the skull. Under the goblet an iron knife with a curved back and a straight blade lay. Chronology according to archaeology: 9th to 8th c. BCE.

**UKR005.** *Kartal, burial 103.* The pit has the shape of a wide rectangle with rounded corners, oriented along the N-S line. At a depth of 25–30 cm a human skull was found. Further, the contours of the burial pit were fully revealed. The size of the pit was 1.7 x 1.15–1.2 m, the depth 45 cm. The skeleton lay on the back. The legs were bent at the knees, arms at the elbows. The hands were crossed on the chest. Body orientation to SSE. Near the skull there was a moulded rhyton-shaped vessel. A bronze item was found under the lower thoracic vertebrae – the tip of a belt in the form of an elongated conical cylinder. On the chest lay a fragment of a polished bowl, decorated with flutes along the edge and inner surface. Chronology according to archaeology: 9th to 8th c. BCE.

**UKR006.** *Kartal, burial 109.* The contour of a very large pit had the shape of a rectangle and was found at a depth of 100–110 cm. Orientation on the SE-NW line. The size of the pit was 260×120–130 cm, depth up to 30 cm. At the centre of the pit’s bottom, a trough-like depression was clearly visible, in which the skeleton was located. The skeleton was poorly preserved. The skull was crushed, the bones of the limbs were not completely preserved. The skeleton was in a crouched position and lay on the right side, with the head oriented to SSE. In front of the skull there was a low black-clay vessel (bowl) and a bronze fibula. Very small (1–1.5 mm) white beads were found around the 4^th^ to 5^th^ cervical vertebrae. Between the left shoulder and the back of the skull there was a small bronze plaque with a loop on the back. Chronology according to archaeology: 9th to 8th c. BCE.

**UKR007.** *Kartal, burial 126.* The burial pit has not been traced. The skeleton lay at a depth of 60– 75(80) cm. The skeleton was in a crouched position and lay on the right side, with the head oriented to SSE. The arms were bent at the elbows, the hands directed to the chin. The artefacts consisted of a small black-clay vessel located near the skull. Chronology according to archaeology: 9th to 8th c. BCE. Chronology according to ^14^C dating 996–831 cal BCE (2767±29 BP).

#### **<u>Scythian culture</u>** (Скіфська культура)

*(Text – I. Shramko, S. Zadnikov)*

On the territory of Ukraine, Scythian culture was widespread in the forest-steppe and steppe zones of the Northern Black Sea Region in the 7th–4th c. BCE. On the Dnipro left bank forest-steppe, the Scythians appear in the second half and last quarter of the 7th c. BCE. The Scythian tribes, their daily life and traditions were described by the ancient Greek historian Herodotus in the middle of the 5th c. BCE.

The landmarks of Archaic Scythian period include kurgans, the burial artefacts of which show signs of the early Scythian culture complex: certain types of weapons, horse bridles, items made in the Scythian “animal style”, as well as imported items of Egyptian or West Asian origin, brought by nomads who moved to the forest-steppe through the Caucasus. One of the largest Scythian kurgan necropolises of the early period was found in the basin of the Sula river (the left tributary of the Dnipro). Large kurgans were studied in the basin of the Vorskla river (left tributary of the Dnipro) and Sukha Grun river (tributary of the Psel river), as well as in the Siversky Donets river basin.

From the last quarter of the 6th c. BCE the presence of new nomadic groups, which advanced to the forest-steppe area from the east and through the Caucasus, is recorded. A change in the material culture occurred in the burials of that time, *e.g.*, antique dishes became more common. Settled population groups were moving from western forest-steppe regions, occupying new territories of Dnipro-Donetsk area and populating the eastern regions of the Ukrainian forest-steppe. Plenty of new unfortified settlements arose. Archaic Scythia was replaced by Classical Scythia. This period is well known from the cultural artefacts of numerous kurgans and settlements of the 5th–4th c. BCE.

The Scythians were establishing relations with the local settled agricultural population, as well as with other contemporary ancient centres since the Archaic period. ^112^

#### Scythian period. Forest-steppe zone of the right bank of Dnipro (Правобережний лісостеп)

*(Archaeology -H. Kovpanenko, B. Levchenko; text – D. Grechko)*

In Scythian times, the forest-steppe zone of the right bank of Dnipro was densely populated by a settled population of local origin ^113^. The natural conditions of the region were favourable for various groups of nomads and semi-nomads that appeared there in the 7th–4th c. BCE ^14,114^. In the second half of the 6th c. BCE, changes in climate and the accompanying shift of natural zones caused the southward migration of forest tribes that left monuments of the Podhirtsiv type ^115^. For these reasons, Porossia (area of the Ros River basin, southward from Kyiv) became a place of active population movement and close interaction between different cultural groups.

Early Scythian kurgans in the Ros River basin have attracted the attention of Scythologists for a long time. Most of the burials were investigated before 1917. Excavations of the Medvin necropolis were carried out in the 1970s and 1980s ^116,117^.

#### **Medvyn** (Медвин)

There are several kurgan groups near the village Medvyn, Boguslav district, Kyiv region, in the Girchakiv forest tract. They are located on the elevated right bank of the Khorobra river (a tributary of the Ros river, a right tributary of the Dnipro). This necropolis belonged to the forest-steppe agricultural population, which preserved archaic burial traditions (decarnation through exposure to the elements and scavangers) ^118^. The time range when the necropolises were used can be attributed to the Zhabotyn period and the beginning of the early Scythian period (second half of the 8th until third quarter of the 7th c. BCE) ^117^.

The burial rite of the Medvyn necropolis has direct analogies in the burial ground of the early Zhabotyn period near the village of Tyutky in the Southern Bug basin, where these traditions have no local basis^119^. Burials with a similar set of artefacts are found in the earlier dated kurgans of Saharna-1 burial ground (Cigleu) in forest-steppe Moldova ^120^. These facts allow to assume the movement of the population from Middle Transnistria (the oldest complexes) through Pobuzhzhia (Tyutky, Nemyriv, Vyshenka-2) to Porossia in the early Zhabotyn period. The migrants moved into regions sparsely populated by people of the late Chernolis culture, where mixing of different ethno-cultural groups occurred. The funeral rite and the set of moulded dishes indicate either the participation of the Chornolis-Zhabotyn population of Porossia in the genesis of this population, or the influence of migrants on the material culture ^121^.

**Location:** 49.40542 N, 30.846856 E. Medvin village, Bila Tserkva district, Kyiv region.

**Excavations:** 1973, 1984.

**Excavation authors:** Halina Kovpanenko, Institute of Archaeology, National Academy of Sciences of Ukraine, Kyiv; Borys Levchenko, Communal institution “Museum of History of Boguslav Region” of Boguslav City Council, Kyiv Region.

**Storage of anthropological materials:** Institute of Archaeology, National Academy of Sciences of Ukraine, Kyiv.

**Description of samples:**

Ten samples were taken for aDNA analysis, seven yielded a sufficient amount of DNA for further study.

**UKR036.** *Medvyn, tract Girchakiv Lis, Group I, kurgan 1, burial 1.* The burial was in a ground pit with a latitudinal orientation. The skeleton was stretched out on its back, the head was oriented to the northwest. The burial was accompanied by artefacts similar to the Hryshkov sets – a quiver set, a spear, meat food ^122^. Chronology according to archaeology: second half of the 5th c. BCE. ^14^C calibrated date 773–426 cal BCE (2481±31 BP).

**UKR042.** *Medvyn, tract Girchakiv Lis, Group I, kurgan 3/1973, burial 1.* The burial was in a rectangular ground pit with a small dromos entrance on the south side. The wooden floor was damaged during the secondary penetration to the grave. The bones of a 35–40-year-old man and a 25–30-year-old woman (UKR042) lay in two elongated parallel clusters with a certain system. Several moulded vessels were found *in situ* near the bones. The southern part of this composition was disturbed because of the secondary penetration into the grave for ritual manipulation. The burial artefacts included a set of moulded dishes (a bowl, three cups and a ladle), bronze and bone arrowheads, mushroom-shaped bone parts of a quiver, necklaces of cowrie shells and opaque glass, and an iron knife. ^116^. Chronology according to archaeology: second half of 7th c. BCE. Chronology according to ^14^C dating: 779–539 cal BCE (2503±30 BP).

**UKR039.** *Medvyn, tract Girchakiv Lis, Group II, kurgan 3/1973, burial 1.* The burial was in a rectangular ground pit with a size of 2.9×1.95 m, 0.6 m deep, with a small dromos entrance on the southern side and two wooden pillars near it. The roof was not preserved. The filling was very dense, indicating secondary penetration. The burial was probably disturbed for a ritual purpose, as the long bones were lying chaotically in the central part of the grave and the skull was closer to the southern wall. Near the skull, pointed to the south, lay an iron spearhead. ^116^. Chronology according to archaeology: second half of 7th c. BCE.

**UKR035A, UKR035B, UKR043, UKR044.** *Medvyn, kurgan 22, field numbers 403, 405, 406.* The burial was in a rectangular pit with a dromos on the south side. In ancient times, the pit was covered with longitudinal and transverse oak logs. Only fragments remain from the ceiling, what indicates a secondary entry into the grave. Judging by the number of skulls, the remains of at least ten individuals (including UKR035, UKR043, UKR044) were placed in the central part of the pit. The bones of nine skeletons were laid out in a certain system, and the skulls were placed near the northern and southern edges. The bones of the “western” and “eastern” skeletons were partially laid out in pseudo-anatomical order. These are completely excarnated secondary burials, buried after the complete loss of ligaments and consist of separate bones. The remains of the dismembered individual, who was buried last, are of interest. It includes the bones of the arm, leg, lumbar spine and pelvis, which were not in the correct anatomical placement. Judging by the destruction of the roof and the presence of bowls above the level of the grave’s bottom, there was a secondary penetration to the grave for the reburial of the dismembered individual. Hence, the burial complex of kurgan No. 22 was a two-act burial (secondary burials with reburial). Some of the bones had traces of cutting tools and teeth of predators. During the reburial of the dismembered skeleton, the entire central part of the chamber was opened. The bones of the ancestors were collected in a “pack” or a pile, and parts of the dismembered body were placed next to or together with it. It was difficult to determine who was buried, it can be assumed that it was a woman. The burial was accompanied by various artefacts. In the south-eastern corner of the pit, near the dromos, three large bowls were found. One of them was lying upside down. Another was above the dismembered skeleton, 35 cm above the bottom, with a ladle with a handle inside. The third one was 50 cm above the bottom, a little away from the others. Next to one skull there was a moulded cup, an iron knife, a spinning wheel, and scattered necklaces. A bronze pendant was found near the northwestern edge of the bone remains. Chronology according to archaeology: third quarter of the 7th c. BCE.

#### Scythian period. Forest-steppe zone of the left bank of Dnipro, Vorskla group of landmarks *(Лівобережний лісостеп, Ворсклинська група пам’яток) (Text – S. Zadnikov, I. Shramko)*

The Scythian culture of the Vorskla group of landmarks is represented by unfortified settlements, hillforts, kurgans, and ground burials. It was formed in pre-Scythian times, in the second half to the end of the 8th c. BCE, when the Basarab tribes – carriers of the Middle Hallstatt culture – moved from the right to the left bank of the Dnipro River, to the basin of the Vorskla River and some tributaries of the Psel River. According to written sources, the tribes of the Neurs, Budins and Gelons lived in this territory in the second half of the 6th to 5th c. BCE and the city of Gelon existed at the last quarter of the 6th to 4th c. BCE. According to archaeological data, the territory of the Vorskla basin was populated by settled tribes, who were engaged in agriculture, cattle breeding and crafts. From the third quarter of the 7th c. BCE the population was in trade relations with the ancient centres of the Northern Black Sea region. From the end of the 8th to the end of the 6th c. BCE local people had tight connections with the Illirian-Thracian tribes of the Northern Balkans, Central Europe, the right bank of the Dnipro and the Scythians, who moved into the region in several waves from the second half to the end of the 7th c. BCE. Burial traditions have regional characteristics, reflecting the diversity of society and complex social composition of the population. ^123–127^.

#### **Bilsk fortified settlement (Bilsk Horodische, Bilsk Hillfort)** (Більське городище) (Archaeology – B. Shramko, I. Shramko, S. Zadnikov; text – I. Shramko, S. Zadnikov)

Bilsk fortified settlement (8th–4th centuries BCE) is known as the largest fortified settlement of the Early Iron Age in Europe and as a major craft, trade, religious and political centre of forest-steppe Scythia. Most researchers identify it with the city of Gelon described in Herodotus’ “History”. It is located in the forest-steppe area on the left bank of the Dnipro river, on the watershed plateau between the rivers Vorskla (tributary of the Dnipro) and Sukha Grun (tributary of the Psel).

The settlement consists of three fortifications connected via an earthen rampart about 35 km in length and occupies an area of 5,000 hectares. There are kurgan necropolises westward of the settlement, in the locations of Skorobir, Osnyagi, Peremirky, Pereshchepyne, and Marchenki. Small groups of kurgans dated to the later period are located within the settlement. On its territory and beyond, separate earthen burials and a large earthen necropolis have been found.

Some unfortified satellite settlements were discovered around the main settlement. The earliest of those were founded in the second half to the end of the 8th c. BCE by migrants from the right bank of the Dnipro, who were carriers of the Basarab archaeological culture. Until the end of the 6th c. BCE, the population maintained contacts with the Hallstatt cultures.

The first excavations were carried out in 1906 by V. O. Gorodtsov. Since 1958 the hillfort and necropolises have been consistently studied by Ukrainian researchers from Kharkiv, Poltava, Donetsk, Kyiv, as well as researchers from Germany. In addition to knowledge of the rich material culture and extensive trade relations of the Scythians with other cultural centres, the complex social and ethnic composition of the population has been traced. The diversity of the society is recorded, first of all by the funeral rite, the presence of graves of representatives of the local aristocracy, elite women’s burials, etc. 124,128–136

**Location:** 50.093324 N, 34.596018 E. Bilsk village, Poltava district, Poltava region.

**Excavations:** 1975, 1978–1980, 1983, 1987–1988, 1990, 1994, 2009, 2011–2021.

**Excavation authors:** Borys Shramko, Iryna Shramko, Stanislav Zadnikov, V.N. Karazin Kharkiv National University, Kharkiv.

**Storage of anthropological materials:** Museum of Archaeology of V.N. Karazin Kharkiv National University.

**Description of samples:**

17 samples were taken for aDNA analysis, seven samples yielded a sufficient amount of DNA for further study.

**UKR078.** *Bіlsk, Western fortification, ash hill 10, field number 7675, square 21–22/Щ–Э, depth 0.60 m, pit 12.* Excavated in 2011. The burial has features of the Illirian-Thracian basis. The pit was found at a depth of 0.50 m, its bottom was at the depth of 1 m from the modern level. The diameter of the pit was 2.45 x 2.70 m. Only a human skull was found in the ashy filling, as well as fragments of moulded ware, a fragment of the wall of a Greek amphora with two red stripes (2nd–3rd quarters of 6th c. BCE), two bronze arrowheads (three blade, one basic), one iron three-bladed arrowhead, a fragment of a bone cheek-piece designed in an animal style, two fragments of iron bracelets. Chronology according to archaeology: the middle of 6th с. ВСE. Chronology according to ^14^C: 755–408 cal BCE (2445±37 BP).

**UKR083.** *Bіlsk, Eastern fortification, excavation 32, field number 205/32-1988, square Я-14, depth 0.50 m., household pit 8.* Excavated in 1988. Supposedly, the individual belonged to the local agricultural population. The pit was found at a depth of 0.20 m, its bottom was at the depth of 0.8 m. It was of a round shape with a diameter of 2.33 m. There were fragments of moulded ware, a bronze bracelet made of thin wire, four clay cone-shaped sinkers in the black soil filling, as well as fragments of a human skull. Chronology according to archaeology: second half of the 5th–4th c. BCE.

**UKR087.** *Bіlsk, Skorobir kurgan cemetery, kurgan 1/2013, burial 1.* Excavated in 2013. The individual belonged to the local elite, the burial has features of the Illirian-Thracian basis. The kurgan had a height of 0.2 m and a diameter of 19 m. Annual ploughing practically levelled the kurgan – the kurgan had almost no external signs. From the southwest (at a depth of 1.55 from the rapper), the burial chamber was adjoined by a dromos on which the burial of a man lying on his back with his head oriented to the west was found. A bronze arrowhead was lodged in the distal part of his left tibia. The burial was built as a wooden crypt. Remains of wooden planks can be traced along the walls of the burial chamber and in its filling. On the chamber’s ceiling there were three sets of horse bridles, which included iron bits, cheek-pieces, bronze clips and zoomorphically decorated plaques. The main burial was looted, but some items (mainly in the southern part of the chamber) remained *in situ*. At the bottom of the southern corner there was a Greek amphora, in the southwestern corner a jar. Along the western wall of the burial chamber there were a 2.27 m long iron spear and several sets of horse bridles (iron bits with cheek-pieces). At 0.2–0.3 m to the east, there were remains of a leather quiver with 147 bronze tips. In the central part, at the bottom of the burial, a part of the skeleton and the skull of a goat were found. A row of beads was found in the eastern part. Fragments of a bronze pin and an iron knife were also found in mixed filling. Not far from the western wall, separate bones and skull fragments of a man and a woman (UKR087) were piled up. A gold plaque 1×1 cm was found in the man’s jaw. Thus, the main burial was a pair (man, woman) and belonged to the elite. Chronology according to archaeology: last quarter of the 6th c. BCE.

**UKR088.** *Bіlsk, Skorobir kurgan cemetery, kurgan 19.* Excavated in 1975. Supposedly, the buried individual belonged to the local elite. The kurgan was looted. The height was 0.8 m, the diameter about 29 m. The rectangular grave pit with dimensions of 2.5×3.3 m was elongated along the SW-NE line. On the south-western side there was an entrance corridor with a width of 1.1 m and a length of 3 m. The floor of the corridor was sloping. The bottom of the grave was at a depth of 1.8 m from the top of the kurgan. The grave was built as a wooden crypt with four main pillars and two small additional pillars at the entrance. The walls and floor were lined with wood. The skeleton was badly damaged. Among the bone remains there were a cranial lid, fragments of the tubular bones of a man’s limbs, a human shoulder blade. Fragments of the lower and upper jaw of a woman were found near the south-eastern wall (UKR088). The finds included an iron knife with a bone handle, pottery fragments, remains of sacrificial food (large bones of a goose), a gold plaque shaped as a stylized hare. Plaques of this type as well as knives with a bone handle were widespread in kurgans of the 4th c. BCE. Chronology according to archaeology: 4th с. ВСE. Chronology according ^14^C dating: 761–420 cal BCE (2467±28BP).

**UKR089.** *Bіlsk, Skorobir kurgan cemetery, kurgan 6.* Excavated in 1979. Supposedly, the buried individual belonged to the local elite. The kurgan was looted. The height of the kurgan was 1.3 m, diameters were 46 m and 42 m. The burial was in a grave pit with dimensions of 3.1×2.2 m. The bottom of the pit was at a depth of 2.3 m from the top of the kurgan. The grave was built as a wooden crypt, made of birch logs 15–20 cm thick, covered with a wooden roof, which was badly damaged by robbers. The bottom of the grave was lined with birch bark. The kurgan was robbed twice. The human skeleton was badly damaged, the bone remains scattered. It was probably oriented along the SW-NE line. The remains of a skull were found in the southwest corner of the grave, and the tubular bones of the legs lay near the northern wall. Fragments of narrow bronze plates from a belt, remains of an iron bit and an iron awl were found. Chronology according to archaeology: 5th–4th c. BCE.

**UKR090.** *Bіlsk, Skorobir kurgan cemetery, kurgan 20.* Excavated in 1975. Supposedly, the buried individual belonged to the local middle-level elite. The kurgan was looted. The height was 1.2 m, diameter 28 m. At a depth of 0.85 m a robbery passage was seen. The grave pit had dimensions of 3.5×4.0 m. The pit was stretched out along NW-SE line and covered with logs. The bottom was at a depth of 2.65 m from the top of the kurgan. The grave had a wooden floor. Most of the human bones were destroyed or thrown into the robbery passage. Only a fragment of the lower jaw of an adult was found near the NE wall of the grave. Among the things scattered around the grave were fragments of a moulded pot, an amphora, a fragment of a bowl and a fragment of a Greek black-glazed vessel of the 4th c. BCE, which determines the chronology according to archaeology: 4th c. BCE.

**UKR091.** *Bіlsk, Pereshchepynе kurgan cemetery, kurgan 3, burial 2.* An inlet burial. Excavated in 1980. Supposedly, the buried individual belonged to the local high-level elite. The height of the kurgan was 0.9 m. The kurgan contained two chronologically close burials, main (1) and additional (2). Burial 2 (UKR091) belonged to a 20–30-year-old man and had rich grave goods. The pit of this burial had the shape of an irregular rhombus with slightly rounded corners (2.9×3.1×2.55×2.75 m), with a depth of 1.3 m. Along the north-eastern wall, a stretched skeleton was laid on the back, with the head oriented to N-NW. The bones were in the correct anatomical order, but almost completely decayed. The remains of the skull and teeth were better preserved, which made it possible to establish the age at death. The artefacts included an iron sword with a golden scabbard, two spears with iron pointed tips and iron shafts. To the northwest of the head of the skeleton there were the remains of a quiver with arrows that had bronze sleeve tips. A total of 57 bronze arrowheads and 5 of their fragments were found. Also, an iron knife, lekythos, Greek ceramic jugs were found in the burial. Chronology according to archaeology: end of 5th until beginning of 4th c. BCE.

#### **Kolomak** (Коломак)

*(Archaeology – V. Radzievska; text – S. Zadnikov, I. Shramko)*

The settlement is located on the cape of an unnamed tributary of the Kolomak River (a tributary of the Vorskla River). The settlement consisted of the main yard (5.4 hectares) and suburb area (8 hectares). Excavations revealed that it was inhabited by the local forest-steppe population of the Scythian period (the second half of the 6th to 4th c. BCE). The territory of the settlement was densely populated. During excavations, 14 dwellings, numerous pits and a well were found. A big number of objects of material culture were found, namely knives, sickles, hoes, arrowheads, clay figurines, fragments of local moulded ware, fragments of Greek amphorae. Features of attacks were traced. The first attack dates back to the 6th–5th c. BCE, when the settlement suffered from the raids of the steppe Scythians. After that it was restored and strengthened with an additional rampart and ditch on the eastern side. In the second half of the 5th c. BCE the territory of the settlement decreased. At the border of 4th to 3rd c. BCE the settlement was again defeated by nomads and ceased to exist. ^137–140^

**Location:** 49.864876 N, 35.277719 E. Kolomak city, Kharkiv region, Ukraine. The headland on the left bank of a tributary of the Kolomak River, which is a left tributary of the Vorskla River.

**Excavations:** 1987, 1991.

**Excavation authors:** Vira E. Radzievska, Museum of Archaeology of V. N. Karazin Kharkiv National University, Kharkiv.

**Storage of anthropological materials:** Museum of Archaeology of V.N. Karazin Kharkiv National University, Kharkiv.

**Description of samples:**

Five samples were selected, DNA was extracted from two of them.

**UKR095**. *Kolomak, Field number 1198/IV-87, pit 51 (fragment of the skull cap of an adult, skull 3).* The burial has been attributed to local agricultural tribes. Pit 51 was found at a depth of 0.40 m from the modern level, size 3.10×3.20 m. The depth of the bottom gradually increased from 0.80 m in the western part to 1.10 m in the eastern part, 1.15 m in the southwest. A large amount of household waste typical for this period was recorded: fragments of moulded ceramics (over 200 specimens), quartzite chips (5 pieces), clay plaster (5 pieces). In the garbage filling of the pit, starting from 0.60 m, scattered human bones were found. Separate parts of at least 4 skeletons (including a tooth – 1198/IV-87, UKR095) and one relatively complete skeleton of a subadult without traces of a ritual were found. All human bones were of poor preservation. Chronology according to archeology: 6th–4th/3rd c. BCE. Chronology according to ^14^C dating: 389–204 cal BCE (2241±27 BP).

**UKR096.** *Kolomak, Field number 1915/IV-88; square БД-3.* A human lower jaw was found in the cultural layer of the settlement, in excavation 4, at a depth of 0.40 m, among cultural remains attributed to local agricultural tribes. Chronology according to archaeology: 6th–4th/3rd с. ВСE. Chronology according to ^14^C dating: 382–199 cal BCE (2220±25BP).

#### **Kupievakha** *(Куп’єваха)*

*(Archaeology – S. Berestnev; text – I. Shramko, S. Zadnikov)*

A group of 76 kurgans is located on the watershed plateau on the right bank of the Berezivka River (right tributary of the Vorskla River). ^123–125,127,141,142^

**Location:** 50.207055 N, 35.285673 E. Kupevaha village, Bohodukhiv district, Kharkiv region, Ukraine.

**Excavations:** 1980, 1992–1993, 2003.

**Excavation authors:** Serhii Berestnev, V.N. Karazin Kharkiv National University, Kharkiv.

**Storage of anthropological materials:** Museum of Archaeology of V.N. Karazin Kharkiv National University, Kharkiv.

**Description of samples:**

Two samples were taken from the burials, DNA was extracted from one of them. Since Kupievakha is located very close to the Siversky Donets basin and this sample is the only nomad sample from the left bank of Dnipro region, it is grouped together with Siversky Donets nomads in analyses.

**UKR105.** *Kupyevaha, kurgan 23, burial 1.* The burial has been attributed to nomads. The primary grave. The pit is square-shaped (3.5×3.6 m), its corners oriented to NE-SW and NW-SE directions. The bottom was at a depth of 2.6 m. On the bottom, along the western wall, lay the skeleton of a child. It was stretched out on the back with the head oriented to the south. The remains of the buried adult individual (UKR105) had been disturbed by robbers: closer to the south-western wall, the parietal part of the skull and individual bones of the limbs were found out of anatomical order. Beads (carnelian, amber, glass?), a small gold plaque of a hemispherical shape with an internal loop, fragments of iron items (a knife) and a Greek amphora made on the island Lesbos were found in the burial. Chronology according to archaeology: the end of 6th to beginning of 5th c. BCE. Chronology according to ^14^C dating cal. 798–552 cal BCE (2547±26BP).

#### <u>Scythian period. Forest-steppe zone of Dnipro-Donets, Siversky Donets group of landmarks</u> (Дніпро-Донецький лісостеп, Сіверськодонецька група пам’яток) (Text – I. Shramko, S. Zadnikov)

The Scythian culture of the Siversky Donets group of landmarks is represented by unfortified settlements, hillforts and kurgans. Until the second half of the 6th c. BCE, the territory was occupied only by nomads, whereas later numerous settlements of agricultural tribes appeared there. The region was settled by migrants.

The graveyards formed compact groups. The main type of burials of the Siversky Donets group of landmarks were ground pits with wooden floor and crypts. Individuals were buried stretched out on the back. Southern orientation of the head prevailed. The ritual use of fire was widespread. Child burials in kurgans were almost completely absent. Along with the burials of the main population of the region in the Scythian period, the graves of other ethnic groups have also been traced.

Tribes that inhabited the Dnipro-Donets forest-steppe zone in Scythian times are classified as Herodotus’ Budins, whose tribal union included the Melankhlens tribe. The local features of the region were sTable from the second half of the 6th to the 5th c. BCE and began to change due to external influence at the end of the 5th and in 4th c. BCE. In the last quarter of the 4th c. BCE, it is possible to assume a Sauromatian-Sarmatian invasion in the region. ^143–148^.

#### Cheremushna (Черемушна)

(Archaeology – Yu. Buynov; text – I. Shramko, S. Zadnikov)

The kurgan burial consisting of about 150 kurgans is located on the edge of the plateau of the left bank of the Cheremushna River (the left tributary of the Mzha River). The burial arose in the Bronze Age and can be classified as medium-sized (group of 50–150 kurgans). Burials were made in ground pits and in inlet catacombs. Also, ritual burning was traced. ^149^

**Location:** 49.854878 N, 35.818011 E. Cheremushna village, Bohoduhiv district, Kharkiv region.

**Excavations:** 2002.

**Excavation author:** Yurii Buynov, V. N. Karazin Kharkiv National University.

**Storage of anthropological materials:** Museum of Archaeology of V. N. Karazin Kharkiv National University, Kharkiv, Ukraine.

**Description of samples:**

One sample was taken for aDNA analysis.

**UKR111.** *Cheremushna, kurgan 10, burial 1.* A primary grave. The burial has been attributed to nomads. The height of the kurgan was 1.25 m. The burial pit was rectangular-shaped (3 x 2.1 m), its depth was 1.7 m. The burial was robbed. Human bones were found in the filling of the lower part of the chamber. At the skull base, traces of its cutting were visible, and on the inner side of the parietal bone there was a round hole, formed as a result of its attachment to a wooden pole. The following artefacts were found in the burial: fragments of an iron knife, four bronze arrowheads, fragments of iron horse bits, two bronze plates from a horse’s bridle, a bronze buckle, a fragment of a Greek amphora and a fragment of a silver fitting from a wooden vessel. Chronology according to archaeology: end of 5th – beginning of 4th c. BCE. Chronology according to ^14^C dating: 775–540 cal. BCE (2496±26BP).

#### **Кaravan** (Караван)

*(Archaeology – V. Okatenko; text – I. Shramko, S. Zadnikov)*

The necropolis consisting of four kurgans is located on a high plateau on the left bank of the Merefa River between the Karavan village and the Lyubotin city, Kharkiv region. ^143–145,148,150–152^

**Location:** 49.925983 N, 35.880553 E. Karavan village, Lyubotyn territorial community, Kharkiv region.

**Excavations:** 2013.

**Excavation authors:** Vitalii Okatenko, State enterprise research centre “Security Archaeological Service of Ukraine” of the Institute of Archeology, National Academy of Sciences of Ukraine, Kyiv.

**Storage of anthropological materials:** Institute of Archaeology, National Academy of Sciences of Ukraine, Kyiv Ukraine.

**Description of samples:**

One sample was taken for aDNA analysis.

**UKR116.** *Кaravan, kurgan 2, burial 2, (central grave (primary burial)).* The burial has been attributed to nomadic elite. The ground embankment was about 5 m in height and was completely destroyed during the excavation. The pit was 1.7 m deep from the ancient surface and had a rectangular shape measuring 4.8 x 4 m. The burial was robbed. In the burial, the upper part of a Greek amphora, eight gold plaques decorated with ornament, fragments of a silver rhyton decorated with a bull’s head, a bronze dish with gilding, bronze arrowheads, fragments of an iron three-looped cheek-peice, and fragments of a moulded scoop were found. Chronology according to archaeology: the first quarter of 6th c. BCE. Chronology according to ^14^C dating: 775–516 cal BCE (2491±28BP).

#### **Mala Rogozianka** *(Мала Рогозянка)*

*(Archaeology – Y. Buynov; text – S. Zadnikov, I. Shramko)*

The group of five kurgans is located 1 km to the north of the Mala Rogozianka village, on a high plateau on the left bank of the Udy River. The distance to the nearest settlements is several km. In terms of size, it belongs to the group of small necropolises. A moat was traced around the kurgan. There were several types of burials: crypts in a ground pit, wooden crypts with pillars and a dromos. Buried individuals were laid on their backs, with heads oriented to the southwest. The burials of nobility were accompanied by the dependent persons. ^143^.

**Location:** 50.13547 N, 35.909989 E. Mala Rogozianka village, Bohoduhiv district, Kharkiv region.

**Excavations:** 1989.

**Excavation authors:** Yurii Buynov, V. N. Karazin Kharkiv National University.

**Storage of anthropological materials:** Museum of Archaeology of V.N. Karazin Kharkiv National University.

**Description of samples:**

One sample was taken for aDNA analysis.

**UKR113.** *Mala Rogozianka, kurgan group I, kurgan 1, burial 3.* A primary grave. Attributed as nomadic. The height of the kurgan was 1–1.1 m. The pit was of rectangular shape (3 x 2.5 m), a wooden crypt was built in it. The skeleton was poorly preserved. It lay on the back with the head oriented to the west. Arrowheads (33 bronze, 4 bone, 1 iron) and a moulded pot were found. Chronology according to archaeology: the first quarter to first half of the 6th c. BCE.

#### Nyzhnia Gyivka *(Нижня Гиївка)*

*(Archaeology – Y. Buinov; text – S. Zadnikov, I. Shramko)*

The burial ground near Nyzhnia Gyivka village is located on the edge of the plateau on the right bank of the Merefa River. It consists of 83 kurgans and belongs to medium-sized burial grounds (50–150 kurgans) of the Siversky Donets group of landmarks. Burials of 4th c. BCE were made in ground pits with “shoulders” around the perimeter. Ritual burning has been detected. ^143–148,150,151^

**Location:** 49.914637N, 35.935222 E. Nyzhnia Gyivka village, Lubotyn city, Kharkiv region.

**Excavations:** 1994.

**Excavation authors:** Yurii Buynov, V.N. Karazin Kharkiv National University, Kharkiv, Ukraine.

**Storage of anthropological materials:** Museum of Archaeology of V.N. Karazin Kharkiv National University, Kharkiv.

**Description of samples:**

Three samples were taken for aDNA analysis, two samples yielded a sufficient amount of DNA for further study.

**UKR101.** *Nyzhnia Gyivka, kurgan 3, burial 1.* An inlet burial. Attributed as nomadic. The height of the kurgan was 0.5 m. The burial was found at a depth of 0.85 m from the benchmark. The contours of the grave-pit were not traced. The skeleton lay stretched out on its back, with the head oriented to the south. There were bronze bracelets on the hands, a temporal ring near the skull, and 18 pyramidal pendants made of blue glass on the chest. Chronology according to archaeology: 4th c. BCE.

**UKR114.** *Nyzhnia Gyivka, kurgan 3, burial 2 (central grave (primary burial)).* The burial is attributed as nomadic. A rectangular burial pit 2.5 x 1.6 m with a depth of 1.8 m. The skeleton, poorly preserved, without a skull, was oriented with its head to the south. A bronze arrowhead was found near the shoulder. Chronology according to archaeology: the end of 5th – 4th c. BCE.

#### **Gryshkivka** *(Гришківка)*

*(Archaeology – D. Grechko; text – S. Zadnikov, I. Shramko)*

The kurgan is located on the plateau of the left bank of the Velika Vylovka River (the right bank of the Mzha River, the right tributary of the Siversky Donets River), 4 km to the east of the watershed along which the Muravsky Way to the steppe passed. It consists of 53 kurgans. In total, 18 graves under 13 kurgans were excavated. All the graves were represented by rectangular pits oriented along the W-E line. Five kurgans were surrounded by moats, where the remains of feasts were found. On five occasions, head orientation to the NW and SE was fixed. Twice, the skeletons of women were placed with their heads facing east. Stretched on the back was the most common body position. The artefacts of men’s burials were represented by bronze arrowheads and iron spearheads. Women’s burials were accompanied by jewellery (necklaces, pendants, bronze and iron bracelets).

The necropolis functioned from the end of the 5th to the first half of the 4th c. BCE. The specific features of the burial rite leave no doubt that it belonged to the steppe Scythians, who moved to the forest steppe in search of free pastures. Burial artefacts are standard for the Scythian culture of the end of the 5th– 4th c. BCE. The necropolis reflects the process of displacement of impoverished groups of nomads from the steppe in the 5th–4th c. BCE. ^153,154^

**Location:** 49.671775N, 36.167723 E. Grishkivka village, Zmiiv district, Kharkiv region.

**Excavations:** 2006.

**Excavation authors:** Denys Grechko, Institute of Archaeology, National Academy of Sciences of Ukraine.

**Storage of anthropological materials:** Museum of Archaeology of V.N. Karazin Kharkiv National University.

**Description of samples:**

Two samples were taken for aDNA analysis, one sample yielded a sufficient amount of DNA for further study.

**UKR104.** *Grishkivka, kurgan 26, burial 2.* The height of the kurgan was about 1 m. Two simultaneous burials attributed to nomads were found there. The pit of burial 2 had a rectangular shape 2.6 x 1.3 m and a depth of 1.3 m. An iron dart, an iron knife with a bone handle, and 29 bronze arrowheads were found. Chronology according to archaeology: last quarter of the 5th – first half of the 4th c. BCE.

#### **Pisochyn** *(Пісочин)*

*(Archaeology – V. Borodulin; text – S. Zadnikov)*

The kurgan necropolis is located on the right bank of the Luchka River (the right tributary of the Udy river), near the southern outskirts of the Pisochyn settlement, Kharkiv district, Kharkiv region. This family necropolis began to form in the second half of 5th c. BCE and continued to function until the end of 4th c. BCE. In total, 35 kurgans with a height varying from 0.3 to 4 m have been explored. The excavated burials date to the period from the middle of 5th to the end of 4th c. BCE. Among artefacts, weapons and gold jewellery were found. The necropolis belongs to the so-called “military retinue’s kurgan necropolises”, and accordingly, the buried individuals had high social status. ^155^

**Location:** 49.945 N, 36.0975 E. Pisochyn settlement, Kharkiv district, Kharkiv region.

**Excavations:** 1978–1980.

**Excavation authors:** Vyacheslav Borodulin, M. F. Sumtsov Kharkiv Historical Museum, Kharkiv.

**Storage of anthropological materials:** M. F. Sumtsov Kharkiv Historical Museum, Kharkiv.

**Description of samples:**

Three samples were taken for aDNA analysis.

**UKR131.** *Pisochyn, kurgan 8.* The height of the kurgan was 6.2 m. Two burials were found in the kurgan. Burial 1 (the inlet burial) was a wooden crypt measuring 3.3 x 3.2 m, oriented along the N-S line. The burial was collective and has been attributed to the nomadic elite. Only teeth and decay of a light-yellow colour remained from the buried individuals. In the middle of the tomb, a woman lay with the head oriented to the south. In her skull area, deciduous teeth were found, as well as the remains of a headdress consisting of gold plates depicting a snake-footed goddess. The earrings were a golden ring with zoomorphic blue glass pendants. There were 15 round gold plaques in the chest area and 7 gold rings on the hands. Also, glass beads, iron tweezers, a moulded jug, black glazed kantharos, and a bronze mirror were found in the burial. The second buried individual (woman) lay parallel to the first, with the head oriented to the south. Teeth and gold earrings were found near the skull, gold plaques depicting a deer were found near the chest, and a gold ring was found near the left hand. The third buried individual (of unknown gender) lay along the northern wall, the head oriented to the east, without burial artefacts. Burial 2 (primary burial) was a double burial, which had been robbed. In a pit of 2.5 x 1.8 m there was a wooden tomb. The skeleton of a teenager (UKR131) was lying on the back along the northern wall, with the head oriented to the east. Foot bones, tibia and partial femurs remained from the second buried individual. Two bronze arrowheads were found. Chronology according to archaeology: third quarter of 4th c. BCE.

**UKR132.** *Pisochyn, kurgan 18.* The height of the kurgan was 6.7 m. The kurgan contained two burials attributed to the nomadic elite. Burial 1 was a robbed double burial. The burial pit was rectangular in shape. The skeleton of a woman was stretched on the back along the northern wall, the head was oriented to the south. Nearby, a bronze mirror with an iron handle, an iron scoop, black glazed kantharos, a bronze cauldron, bones of sacrificial food, and an iron knife with a bone handle were found. The bones of the second skeleton (UKR132) were scattered in the plundered pit, where 13 gold buttons, around 10 glass beads, a Greek amphora, and two iron darts were found. Burial 2 was in the centre of the kurgan. The burial pit was rectangular in shape and measured 4.2 x 2.2 m. The burial was robbed. Human bones, 5 glass beads, 2 gold and 3 glass amphora-like beads, 3 iron pendants were scattered along the bottom of the chamber. Chronology according to archaeology: mid-third quarter of 4th c. BCE.

**UKR133.** *Pisochyn, kurgan 6.* The burial has been attributed to the nomadic elite. The height of the preserved part of the kurgan was 4 m. The burial pit was rectangular in shape and measured 3.2 x 2.25 m. The skeleton lay along the western wall, with its head oriented to the south in a straightened position.

Near the skull, a golden neckband (hryvnia) and a silver earring were found. There were silver rings on the fingers. In the belt area, there were the remains of a leather belt with bronze plates, a iron shield plates, a quiver with 75 bronze arrowheads. Also, a spearhead and two darts, a Greek pottery jug, and an iron knife with a bone handle were found. Chronology according to archaeology: mid-second half of 4th c. BCE.

#### **Vesele** (Веселе)

*(Archaeology -B. Shramko; text – S. Zadnikov, I. Shramko)*

The group of six kurgans is located on the edge of the watershed plateau, on the right bank of the Murom River (right tributary of the Kharkiv River – Udy River, Siversky Donets basin), between Liptsy village and Vesele village. The burial was founded in the Bronze Age, in pre-Scythian times, and had been used during the Scythian period. Two types of graves were discovered under the kurgans: ground pits with supporting pillars and a ceiling, as well as a catacomb. The burial was explored in 1978 by an expedition of the Kharkiv University under the leadership of B. A. Shramko. ^143–145,148,151,156^

**Location:** 50.183969N, 36.496846 E. Vesele village, Kharkiv district, Kharkiv region.

**Excavations:** 1978.

**Excavation authors:** Boris Shramko, V.N. Karazin Kharkiv National University, Kharkiv.

**Storage of anthropological materials:** Museum of Archaeology of V.N. Karazin Kharkiv National University, Kharkiv.

**Description of samples:**

Two samples were taken for aDNA analysis.

**UKR109, UKR110.** *Vesele, kurgan 4, burial 1.* The burial was attributed as nomadic. The kurgan was 1.65 m high. The burial chamber was made as a catacomb. The entrance chamber had two steps. The bottom of the grave was at a depth of 5.20 m above the level of the reference point. The burial was robbed. The remains of two skeletons were found in the grave, the bones were randomly scattered in the burial chamber. Two human skulls were found near the centre of the grave. One of them (UKR109) had a lifelong deformity of the skull that gave it an unusually elongated shape. Other body bones (UKR110) were found in different parts of the burial. The preserved things were also scattered. Fragments of an iron awl, an iron hook for hanging a quiver, a bronze quiver, an iron two-hole cheek-piece, an iron ring from a fishhook, two gold hemispherical plaques with black glazed Attic camphor of the 4th century BCE were found. Chronology according to archaeology: 4th c. BCE.

#### Scythian period. Steppe zone of the Northern Black Sea region (Причорноморський степ)

Scythian tribes appeared in the Northern Black Sea region in the 7th c. BCE. Most known Scythian landmarks in the region (fortifications, settlements, kurgans) are dated to the 2nd half of 5th to the 4^th^c. BCE. Scythians were engaged in agriculture and cattle breeding, and had close ties with the Greek colonies. A large number of burial sites are concentrated in the Lower Dnieper area, such as Chortomlyk, Solokha, Gaimanova Grave and others, related to the Scythian aristocracy. The Mamay-Gora, one of the largest in the Black Sea steppes, was a burial place mainly for the ordinary population.^78–82^

#### **Mamay-Gora** (Мамай-Гора)

*(Archaeology – G. Toshev, S. Andrukh; text – G. Toshev, I. Shramko, S. Zadnikov)*

The archaeological site has been described under the Neolithic Azov-Dnieper culture section.

In Scythian time, the Mamay-Gora was the burial ground of the ordinary population. It is one of the largest necropolises in the Black Sea steppes: more than 400 Scythian kurgans and ground burials have been found there. Individuals belonging to Scythian time (adults and children) were buried in catacombs and ground pits. They were stretched out on their backs, with their heads mainly to the west. The accompanying grave goods included household items, jewellery, weapons, and rare items of Greek origin. These artefacts date the Scythian burial of Mamay-Gora to the second half of the 7th to to the first third of the 3rd c. BCE. ^157,158^

**Location:** 47.432845 N, 34.27259 E. Mamay-Gora tract, Velyka Znamianka village, Vasyliv district, Zaporizhzhia region.

**Excavations:** 1989, 2002.

**Excavation authors:** Svitlana Andrukh, Gennady Toshev, Zaporizhzhia National University, Zaporizhzhia.

**Storage of anthropological materials:** Zaporizhzhia National University, Zaporizhzhia.

**Description of samples:**

Two samples were taken from Mamay-Gora burials of the Scythian times ^157,158^.

**UKR013.** *Mamay-Gora, kurgan 33, burial 1.* Excavated in 1989. Attributed as nomadic. The burial was found at the depth of 1,2 m. The entrance pit of 2.8 m in length was extended along the W-E line. The roof collapsed. At the depth of 0.15 m there was a step along the long southern wall, 0.4–0.6 m in width. The burial chamber was located parallel to the entrance pit from the north. It was of an oval shape, 2.8×1.1 m. The chamber’s bottom was 0.25 m lower than the bottom of the entrance pit. The burial had been robbed in ancient times. Some human bones were found in the filling. A group of bones of the upper part of the body were found at the western end of the chamber. The bones of the legs and the right hand were *in situ*. Judging by their location, the skeleton lay stretched out on its back, with the head oriented to the west. Among the bones, a clay whorl, six grey beads, and a fragment of an iron bracelet were found. Chronology according to archaeology: 4th с. BCE. ^78^

**UKR014.** *Mamay-Gora, kurgan 173, burial 1.* Excavated in 2002. Attributed as nomadic. The burial had been robbed. It was in a catacomb of oval shape (2.6×0.7 m), elongated along the west-east line. The entrance was found at a depth of 1.2 m. At a depth of 0.85 m, on the south side, a 0.25 m wide step was traced, descending obliquely to a depth of 1.2 m. The depth of the bottom was 1.7 m. The chamber was oval, its length was 2.8 m, the width in the western part was 1 m and in the eastern part 0.55 m. The filling of the entrance pit and the chamber consisted of dense black soil with a slight admixture of loam. In the western part of the entrance, at a depth of 0.6 m, there were human bones (skull, shoulder blades, ribs) and an animal bone. Among them, a knife handle and fragments of a spear were found. In the bottom part of the chamber, the rest of the human bones lay. In the western corner, bones of a large animal (a young cow) with fragments of iron staples and remains of a wooden tray were found. In the centre of the chamber, closer to the northern wall, one three-bladed and one triangular arrowhead were found. Chronology according to archaeology: 4th с. BCE. ^81^

#### **<u>Late Scythian period of Crimea</u>** (Пізньоскіфський період у Криму)

The Late Scythian culture of Crimea (known as Taurica in the Roman era) is represented by settlements, kurgans and ground burials in the steppe part and foothills of the peninsula. It emerged during the late Hellenistic period (end of the 4th–3rd c. BCE) and persisted until the 3rd c. CE ^159,160^. The Late Scythian culture encompassed remnants of various ethnic groups that inhabited Taurica at that time: Scythians, Tauri (descendants of Cimmerians), Thracians, and Sarmatians. The formation of the culture is associated with the gradual sedentarisation of the barbarian population, accompanied by changes in economic activities.

Funerary structures of the Late Scythian culture included earthen and stone tombs (crypts), as well as pit-type graves with a niche. The material assemblage is represented by a wide range of clay vessels (pots, footed bowls, incense burners), antique ceramics (amphorae and Tableware), arrowheads, and tools. Anthropological materials have been studied at sites such as Naples, Belyaus, and Zavyetne ^159^.

#### **Maslyny** (Маслини)

*(Archaeology – O. Latysheva, text - V. Kotenko)*

The kurgan burial site belonging to Late Scythian culture is located in the steppe zone of the Northwestern coast of the Crimean Peninsula, near the Severne village, in the Chornomorsk district of the Autonomous Republic of Crimea. It is situated about 1 km southeast of the Greek settlement of Maslyny, which dates from the 4th to the mid-2nd century BCE. In the Hellenistic period, the settlement of Maslyny was part of the chora of Tauric Chersonesos ^161,162^.

The burial sites in the region are dated to the late 5th to the early 3rd c. BCE. The height of the barbarian kurgans at the Maslyny settlement ranged from 0.5 to 0.7 m. One of them was archaeologically examined in 1976. Artefacts typical of the Late Scythian archaeological culture were found in the burial. The nature of the material obtained (arrowhead, handmade vessel) allows us to speculate that it was not a Greek but a barbarian kurgan, which was located near the Greek settlement. Similar burial sites have been repeatedly investigated in the region and are associated with the presence of Scythians on these lands. In particular, such tombs are typical for kurgans in Western Crimea, dating from the late 5th to the early 3rd c. BCE ^163^. Based on known analogies, the concentration of bones on the flooring may suggest that it was a familial burial ground of sedentary Scythian population. However, with only one kurgan excavated, confirming these conclusions is currently challenging.

**Location:** 45.685922 N, 33.041920 E; Severne village, Chornomorsk district, AR Crimea, Ukraine.

**Excavations:** 1976.

**Excavation authors:** Latysheva V.O., V.N. Karazin Kharkiv National University.

**Storage of anthropological materials:** Museum of Archaeology of V.N. Karazin Kharkiv National University, Kharkiv.

**Description of samples:**

Four samples were taken for aDNA analysis, three yielded a sufficient amount of DNA for further study.

**UKR051, UKR052, UKR053.** *Maslyny, kurgan 1, burial.* The kurgan’s height was about 0.7 m. It was surrounded by a stone ring (width 0.2–1.2 m) and had a limestone pavement (2.2×2 m) at the level of the ancient ground. On the pavement, poorly preserved human bones were found, not in anatomical order (burial position and orientation are unknown). Among the bones, 7 skulls were found (including UKR051, UKR052, UKR053), two of which, judging by the teeth, belonged to children. The artefacts were characteristic of barbarian burials (clay bowl, triangular bronze arrowhead, 10 beads). At 2 m to the east from the centre of the kurgan, another burial was discovered under the stone ring (depth 0.5 m from the reference point). This burial exhibited better preservation, skeleton lay in a contracted position on the left side, head oriented to the northwest, facing to the northeast. The left arm was bent at the elbow, the hand near the face, while the right arm was slightly bent and lowered downward. Among the accompanying artefacts, only an iron knife was found near the right femur. Chronology according to archaeology – end of 2nd to 1st c. BCE.

#### Greeks of Antiquity on the Northern Black Sea coast *(Античні греки у Причорномор’ї)*

*(Text - O. Smyrnov)*

During the ancient Great Greek colonization (8th–6th c. BCE), one of the primary directions was the Northern Black Sea region (mid-7th to 5th c. BCE). For a thousand years, the Lower Southern Bug Region represented the most intense point of penetration for antique culture in Southern Ukraine.

In the mid-6th c. BCE, migrants from Miletus founded the city-state of Olbia. It was located on the right bank of the Southern Buh Estuary near the village of Parutyne in the Ochakov district of the Mykolaiv region. From the 6th c. BCE to the 3rd c. CE, Olbіa represented a typical Greek polis. It had a functioning temenos and an agora as centres of socio-economic, political, and cultural life. Local currency was minted, a temple was erected, along with trading rows and a gymnasium. Defensive stone walls and towers were built around the city. Olbіa had strong trade and cultural ties with the population of the South Black Sea region, as well as with Mediterranean ancient cities like Ionia, Chios, Samos, Rhodes, Corinth. Like other Greek cities, the Olbian state encompassed a substantial agricultural territory known as the chora. The archaeological sites of the chora are represented by over 150 objects (settlements, necropolises) from various periods of antiquity (archaic, classical, Hellenistic, and Roman times).

Olbia’s history is divided into two stages. The first stage (mid-6th c. to 49–44 BCE) is characterised by gradual economic and cultural development, which was interrupted by a Getae (Thracian-related tribe) attack (the mid-1st c. BCE). In the subsequent stage (second half of the 1st c. BCE to the middle of the 3rd c. CE), Olbia faced intense pressure from local tribes and eventually fell into political dependence on Rome. In the mid-3rd c. CE, presumably due to the Gothic invasion, the city ceased to exist as an ancient centre.

#### **Oleksandrivskyi necropolis** (Олександрівський некрополь) (Archaeology, text – O. Smyrnov)

The Oleksandrivskyi necropolis dates to the transition between the 4th and 3rd c. BCE. It is located within the territory of the Admiralteysky Park in the Mykolaiv city centre (the intersection of Naberezhna Street and 2nd Slobidska Street). This necropolis was situated near the ancient settlement known as “Zavod im. 61-Komunara”, which dates to the times from the 4th c. BCE to the 1st c. CE.

The necropolis was discovered in 2010, but its exact dimensions and boundaries have not been established due to urban development. A total of 25 burials were discovered at this site, arranged in a cluster-like pattern.

During the Hellenistic period, from the 4th to the 3rd c. BCE, the territory of present-day Mykolaiv was part of the Olbia state (Olbia chora). This period was characterised by the expansion of the agricultural region of Olbia, which served as a vital economic base for the city-state following the invasion of the Macedonian army. ^164–170^

**Location:** 46.9724 N, 32.00901 E. Naberezhna Street, Mykolaiv city.

**Excavations:** 2011.

**Excavation authors:** Olexandr Smyrnov, Petro Mohyla Black Sea National University, Mykolaiv, Ukraine.

**Storage of anthropological materials:** Laboratory of Archeology and Ethnology, Petro Mohyla Black Sea National University.

**Description of samples:**

Three samples were taken for aDNA analysis and two yielded a sufficient amount of DNA for further study.

**UKR152.** *Oleksandrivskyi necropolis, burial 5.* Burial with a grave niche. Partially disturbed by modern construction activity. The entrance pit and burial chamber were not traceable. The stone structure at the entrance to the burial chamber was composed of limestone slabs, with one of the slabs serving as a limestone altar of the ’Olbia type’. To the north, there was a skeleton extended on its back, with the head oriented to the northeast. The right arm was straight, and the left arm was on the pelvis. Only the femurs remain from the legs. Chronology according to archaeology: 4th–3rd с. BCE. Chronology according to ^14^C dating: 392–206 cal BCE (2253±29 BP).

**UKR153.** *Oleksandrivskyi necropolis, burial 6.* Burial with a grave niche. Partially disturbed by modern construction activity. The entrance pit measured 1.28 m in length and 0.26 m in width and was oriented along the W-E line. The grave-pit was separated from the burial chamber by a double-row stone masonry. The chamber was rectangular, measuring 2.32 m in length and 0.69 m in width, with an orientation along the W-E line. The skeleton lay in an extended position on the back, with the head oriented to the west, arms positioned near the pelvis, and straight legs. Near the left shin, bones of sacrificial food (shallow horned cattle) were found. Twelve arrowheads were discovered near the left femur (possibly the remains of a quiver), additionally, 9 arrowheads were found in other places of the burial. All 21 arrowheads were made of bronze and belonged to three types: 8 three-bladed with a prominent bushing, 6 three-bladed with a less prominent bushing, and 7 triangular ones with a concealed bushing. According to archaeological artefacts, the burial dates to the 4th–3rd c. BCE. The chronology according to ^14^C dating: 746–401 cal BCE (2415±30 BP).

### IRON AGE

#### Sarmatian Period

Sarmatians were a confederation of Iranian-speaking nomadic tribes that originated in the central Eurasian Steppe and were closely related to Scythians. They dominated the North Pontic steppe and forest-steppe from the 3rd c. BCE to the 4th c. CE. Sarmatian nomadic society, most likely, was a temporary military-political association with such attributes of statehood as a common territory, supreme power, a certain social stratification of society, possibly a professional army. Sarmatians had high military potential and imposed tribute and indemnities, controlled over trade routes, used mediation and protectionism in trade, and finally, robbery during raids.

The main occupation of the Sarmatian tribes on the territory of Ukraine was nomadic cattle breeding. They bred mainly horses and sheep. The fighting horse was an important part of the military culture of the Sarmatians. Early Sarmatian burials are scattered across the steppe without any system, most of them are the graves of warriors. This indicates that the initial advance of the Sarmatians into the North Pontic steppes was carried out by military detachments. The lack of settlements, the finds of weapons and horse equipment in ground burials indicate a nomadic lifestyle.

The Sarmatians had contact with other populations of the region: with the inhabitants of ancient cities, the Hellenes; Late Scythian tribes of the Lower Dnipro and Crimea; carriers of the Zarubinets culture and the population of the Chernyakhiv culture. The spread of Sarmatian-type items — weapons, horse equipment, some forms of clothing, and jewellery — in the material culture of the ancient centres of the first centuries BCE got the name Sarmatization of the ancient world.

In the 3th–5th c. BCE Sarmatians were conquered by Goths and Huns, later they were assimilated by Slavs.

#### Forest-steppe zone of Dnipro-Donets. Siversky Donets group of landmarks **(Дніпро-Донецький**

*лісостеп, Сіверсько-Донецька група пам’яток))*

The Siversky Donets group of landmarks has been described under the Scythian period section.

#### **Liubivka** (Любівка)

(Text – I. Shramko, S. Zadnikov)

The multi-layered settlement of Liubivka is located on the first floodplain terrace of the right bank of the Merla River (a right tributary of the Voronezh River within the Dnipro River basin). The surface of the settlement is subject to annual erosion, and there are visible ash spots on the surface. Artefacts from the Neolithic, Bronze Age and Scythian times, Saltiv, Bondarikha and Chernyakhiv cultures, early Slavs period were found in Liubivka and its surroundings. In 1974, three excavations were conducted at the settlement, revealing a cultural layer from the Scythian period (the latter half of the 6th c. BCE). Additionally, isolated fragments of pottery from the Bronze Age were discovered, representing various cultures such as Catacomb, Zrubna (Srubnaya), and Bondarikhinska. Furthermore, a unique Sarmatian girl’s burial was excavated. ^171,172^

**Location:** 50.007291 N, 35.038043 E. Liubivka village, Bohodukhiv district, Kharkiv region.

**Excavations:** 1974.

**Excavation authors:** Borys Shramko, V.N. Karazin Kharkiv National University.

**Storage of anthropological materials:** Museum of Archaeology of V.N. Karazin Kharkiv National University.

**Description of samples:**

One sample was taken for aDNA analysis.

**UKR160.** *Liubivka, Excavation 3, burial 1.* The burial was embedded in the cultural layer of a Scythian-period settlement and was defined as Sarmatian based on the artefacts found. The bottom of the trapezoid-shaped burial pit was in yellow sand at a depth of 1.35 m. The skeleton of a woman was found lying on her back in an extended position, with her head oriented to the north-northeast. On her right clavicle, there was a bronze fibula (brooch) with an iron pin. On the left side of her chest, there was another bronze fibula (brooch) with a suspension loop. Bronze bracelets were on both her arms. Around the neck, 43 glass, coral and amber beads were found. Chronology according to archaeology: 1st – 3rd с. CE.

#### Chernyakhіv Culture (Chernyakhiv-Sântana-de-Mureș Culture) *(Черняхівська культура)*

*(B. Magomedov, S. Didenko, O. Petrauskas, R. Reida, A. Heiko, S. Sapiehin)*

The Chernyakhiv culture (Chernyakhiv-Sântana de Mureș culture) was created by barbarian tribes in late Roman times in Southeastern Europe. It was widespread in most of the forest-steppe and steppe regions of Ukraine, Moldova, parts of Romania, and some neighbouring areas of the Russian Federation from the beginning of the 3rd to the middle of the 5th c. CE ^28^.

Most researchers consider the composition of this culture to be polyethnic. It included Eastern Germans, early Slavs, Getae-Dacians, Sarmatians, as well as late Scythians and Alans ^28^. It is also likely that some portion of the provincial Roman population and immigrants from ancient centres of the Northern Black Sea region were included in it ^27,173–176^. Now this is confirmed not only by archaeology but also by anthropological data ^177,178^. According to most researchers, the leading role in this tribal alliance was played by East Germanic tribes Goths and Gepids ^28,179,180^.

The analysis of archaeological materials from Chernyakhiv sites shows that there are strong traditions of the Wielbark culture and other Germanic cultures of Central Europe in the types of structures, burial rites, moulded ceramics, clothing details, ornaments, and everyday items. The style of pottery developed as a result of the synthesis of the provincial Roman craft tradition and the Wielbark tradition of clay pottery. Thanks to the strong influence of late antique centres, the material culture of Chernyakhiv-Sântana de Mureș reached a high level.

The dominant ethnic group within the Chernyakhiv population were the carriers of the Wielbark tradition – the Goths. They left behind “classic” Chernyakhiv artefacts, which can be conditionally referred to as “Kosanov type” landmarks.

Two local groups of antiquities were related to other ethnic groups. In the Upper Dniester region, there was a western group of Slavs, represented by the “Cherepin type” landmarks. In the Black Sea steppe, there were late Scythians and Sarmatians-Alans who formed mixed communities with the Germanic people. This population left behind “Black Sea type” landmarks. Some Chernyakhiv sites in Eastern Ukraine have elements of the neighbouring Kiev culture (eastern group of Slavs) whereas elements of the Thracian-Carpi culture are found in Moldova and Muntenia, but they do not form independent types.

After the arrival of the Huns, the Germanic and Scytho-Sarmatian population left the Chernyakhiv culture area. Slavic tribes began to dominate in the forest-steppe zone, eventually giving rise to the Prague and Penkivska cultures.

#### Chernyakhiv culture in the Dnipro left bank forest-steppe (Черняхівська культура лівобережного лісостепу)

The Dnipro left bank forest-steppe in late Roman times was part of the Chernyakhiv archaeological culture area. Hundreds of sites are known in this region now ^181,182^.

It is evident that the ethnic components within the Chernyakhiv culture had varying proportions and significance in different regions and during different periods of the culture. For the Dnipro left bank forest-steppe, the Eastern Germanic component is known (as seen in the Kompanitsi burial) ^183^ but relatively limited. Instead, an important ethnic component, at least from the middle of the 4th c. CE, consists of migrants from a nomadic (adjacent to the Chernyakhiv culture area) environment and possibly a late Scytho-Sarmatian ethnic component ^176,184^. The contact zone between nomadic and settled tribes at that time passed through the Vorskla River basin. The nomadic tribes were represented by the Alans. The result of these contacts was the active integration of this ethnic component into the Chernyakhiv community, which can be traced through specific archaeological sites from southeast to northwest of the Dnipro left bank forest-steppe. To observe an example of this process, one can examine the cemeteries of Storozhove ^185^, Kantemirivka ^186^, and Shyshaky ^184^.

The Chernyakhiv culture of the Dnipro left bank forest-steppe retained its character even after the arrival of the Huns in the region and, in some cases, continued to develop. Notably, the Shyshaky and Voitenky cemeteries serve as vivid examples of this. It is likely that the relationships with newcomers among the bearers of the Chernyakhiv culture in the region did not have dramatic consequences, and contacts with ancient centres of the Northern Black Sea and Roman provinces persisted (although the nature of these contacts — whether trade or military interactions — remains unclear).

#### Shyshaky *(Шишаки)*

*(Archaeology, text - A. Heiko, R. Reida, S. Sapiehin)*

The Shyshaky cemetery is one of the largest necropolises of the Chernyakhiv culture in the Dnipro left bank forest-steppe as well as in the entire Chernyakhiv culture area. It is located near the urban-type settlement of Shyshaky (Myrhorod district, Poltava region). Archaeological excavations at the site began in 2009 and continued until 2017. From the beginning of archaeological work, 156 burials were excavated.

The cemetery is characterised by both inhumations and cremations. The predominant group consists of inhumation burials with the heads oriented to the west. The next in terms of quantity are inhumation burials oriented to the north. There are individual inhumation burials with eastern and southern orientations. Burial-cremations constitute the smallest group. The inhumation burials were divided into four types: 1) rectangular pit; 2) ledge-pit; 3) catacomb; 4) tomb ^184^. Some burials exhibit signs of nomadic burial traditions, which, in our opinion, belong to the Alanic component. Among these signs are ledge-pits and catacombs, rituals involving fire, bird eggs, and bones. Elements that could be clearly attributed to other ethnic components, including Eastern Germanic, were not found at the Shyshaky cemetery.

Among other research of the site, it is worth mentioning the DNA analysis conducted on the content of the clay vessel from burial 124 ^187^.

All the examined burials at the Shyshaky cemetery, where chronological indicators were found, were dated within the period from the mid-4th to the first half of the 5th c. CE ^173,188^. These burials primarily include complexes with glassware of provincial Roman production ^173,189^.

**Location:** 49.856205 N, 33.981227 E. Shyshaky village, Shyshaky district, Poltava region.

**Excavations:** 2009–2017.

**Excavation authors:** Anatoliy Heiko (2009-2011), “Center for Protection and Research of Archaeological Monuments” of the Poltava Regional Council; Roman Reida (2012-2017), Institute of Archaeology, National Academy of Sciences of Ukraine, Kyiv, Ukraine.

**Storage of anthropological materials:** “Center for Protection and Research of Archaeological Monuments” of the Poltava Regional Council, Ukraine.

**Description of samples:**

Ten samples were taken from Shyshaky burial, seven of them yielded a sufficient amount of DNA for further study.

**UKR121.** *Shyshaky, burial 88.* Excavated in 2014. Burial of an adult woman. The skeleton was lying flat on its back, the head was oriented to the north. The upper part of the postcranial skeleton was broken. The upper limbs were stretched along the body, the bones of the lower limbs were slightly displaced. The burial had numerous accompanying artefacts: 11 vessels were found in the grave (a large vase-bowl, two-handled jug, decorated pottery cup, bowl-dish, pot, vase-bowl, a small single-handled jug of antique production, a medium-sized bowl, a small bowl, and two pots). Additionally, there were two silver fibulae, a necklace made of carnelian, glass, coral, and amber beads, a small silver belt buckle, a horn comb, a large spindle, a bone awl, and a miniature bronze knife. Next to the pelvic bones, there were bones of large horned cattle, presumably representing accompanying food. Based on the chronological indicators, the burial is dated to the second half of the 4th – the beginning of the 5th c. CE.

**UKR122.** *Shyshaky, burial 89.* Excavated in 2014. Burial of an adult individual. The skeleton was extended on the back, the head oriented to the west. The bones of the upper limbs were on top of the pelvic bones, and the lower limbs were extended. The cranial and postcranial skeleton was relatively well-preserved. No artefacts were found. Chronology according to archaeology: 4th с. CE.

**UKR123.** *Shyshaky, burial 94.* Excavated in 2014. Burial of an adult individual. The skeleton was extended on the back, the head oriented to the west with a slight deviation to the south. No torso bones were found in the burial, the bones of the upper limbs were positioned parallel to each other, while the legs were closely brought together. There were no accompanying artefacts. Chronology according to archaeology: 4th с. CE.

**UKR125.** *Shyshaky, burial 97.* Excavated in 2014. Burial of an adult individual. At a depth of 1.96 m, a cluster of anthropological remains was found, consisting of two femurs, a pelvic bone, several ribs, and vertebrae, chaotically arranged in the western sector of the pit. Subsequent research revealed another part of the burial that remained undisturbed. The skeleton was placed on the back in an extended position, the head was oriented to the west with a slight deviation to the south. The skull, pelvic bone, and possibly the lower jaw, had been displaced at some point after the burial. Traces of wood were found – remnants of the burial structures. No accompanying artefacts were found. Chronology according to archaeology: 4th с. CE.

**UKR126.** *Shyshaky, burial 103.* Excavated in 2014. Burial of an adult individual. The skeleton was extended on the back, the head oriented to the west. Near the right shoulder, a small pot was found, surrounded by animal bones, which were presumably remnants of accompanying food. Chronology according to archaeology: 4th с. CE. Chronology according to ^14^C dating: 131–325 cal CE (1820±26 BP).

**UKR128.** *Shyshaky, burial 114.* Excavated in 2014. The skeleton was of satisfactory preservation, laying on the back in an extended position, the head was oriented to the southwest. Some small bones from the upper and lower limbs were missing, likely due to the activities of burrowing animals. No accompanying artefacts were found. Chronology according to archaeology: 4th с. CE. Chronology according ^14^C dating: 229–361 cal CE (1771±26BP).

**UKR129.** *Shyshaky, burial 115.* Excavated in 2014. Burial of an adult individual. The skeleton was extended on the back, the head oriented to the west with a slight deviation to the south. The skeleton was severely damaged: only fragments of large bones from the upper and lower limbs were found, and the cranial skeleton was heavily damaged. The burial stands out for its relatively rich artefacts, including two belt buckles with fragments of leather belts (one silver, the other made of a copper alloy), remnants of heavily damaged fragments of a horn comb with copper alloy rivets, a pottery bowl with a conical glass cup inside, adorned with drops of blue and dark red colour; another three pottery bowls and a black-glazed jug of the left-bank type. Based on the chronological indicators found in the burial, the complex is dated to the late 4th to early 5th c. CE. ^190^. Chronology according ^14^C dating: 245–401 cal CE (1741±27BP).

#### **Komariv** (Комарів)

*(Archaeology, text – O. Petrauskas)*

The Komariv settlement is located in the middle reaches of the Dniester River – this river was a boundary between the eastern and western parts of the Chernyakhiv-Sântana de Mureș culture. The settlement existed from the beginning of the 3rd to the middle of the 5th c. CE and was a centre oriented towards mass production of glass items such as cups, beads, gaming pieces, etc ^191^. It was the only glass workshop outside the Roman Empire within the Barbaricum territory in Europe. It is highly likely that Komariv was not only influenced by provincial Roman technologies but also had a significant population of Romanized individuals ^69,192^.

The ethnic and cultural composition of the population in Komariv was heterogeneous and included various ethnic groups and social strata, originating from the late antique, Scytho-Sarmatian, early Slavic, Dacian, and East Germanic backgrounds. Dominant ethnic groups included the late Scythians, Sarmatians, and early Slavs, while to a lesser extent, there were Getae-Dacians and people from Roman provinces. The East Germanic component was very restricted in this context ^69^.

**Location:** 48.535183 N, 26.9676 E. Komariv village, Dnistrovskyi district, Chernivtsi region.

**Excavations:** 2013.

**Excavation authors:** Oleg Petrauskas, Institute of Archaeology, National Academy of Sciences of Ukraine, Kyiv, Ukraine.

**Storage of anthropological materials:** Archaeological research Laboratory, National Pedagogical Dragomanov University, Kyiv, Ukraine.

**Description of samples:**

One sample was taken for aDNA analysis.

**UKR049.** *Komariv, burial 2.* According to archaeological data, burial 2 exhibited traits typical for late Scytho-Sarmatian population (ledge-pit) and belonged to a woman under the age of 20. The grave depth was 2.67 m. Remains of wooden roofing were preserved. The head of the skeleton was oriented to the north. The skeleton was ritually destroyed in ancient times. In the burial, the following items were found: 15 glass beads of prismatic shape; a silver fibula clasp; two spinning wheels; six pottery vessels. In addition, the burial was accompanied by a sacrificial animal (a rooster). The artefacts and the position of the burial in the overall plan of the cemetery allow dating it to the second half of the 4th to the beginning of the 5th c. CE. Chronology according ^14^C dating: 169–338 cal CE (1805±26 BP).

#### **Lehedzyne** (Легедзине)

*(Archaeology, text - B. Magomedov, S. Didenko)*

The cemetery of Chernyakhiv culture near the village of Lehedzyne was excavated by B. V. Magomedov and S. V. Didenko in 2008–2009. Approximately half of the necropolis (690 sq. m) was excavated. A total of 81 burials were studied, including 47 cremations and 34 inhumations. The cremations included both with urn and without urn type. Among the inhumations, 2 were conducted in pits with niches, and 3 in ledge pits. The findings were mostly characteristic of the Chernyakhiv culture archaeological artefacts. Wheel-made pottery predominated among ceramics. Moulded vessels had analogies in the Wielbark culture. Imported pottery included fragments of amphorae, red-lacquered items, and glass cups. The site yielded clothing and items of everyday use, such as fibulae and buckles, various beads, pendant amulets, horny and iron combs.

The Lehedzyne burial site dates to the late 3rd – the end of the 4th c. CE. The locals were primarily Gothic people, who constituted the majority of the Chernyakhiv culture population. Burials in pits with a niche and with a ledge may suggest the presence of people of Sarmatian origin within the community.^28,193,194^

**Location:** 48.806513 N, 30.508557 E. Lehedzyne village, Talniv district, Cherkasy region.

**Excavations:** 2008-2019.

**Excavation authors:** Borys Magomedov, Institute of Archaeology, National Academy of Sciences of Ukraine, Kyiv; Serghii Didenko, National Museum of History of Ukraine, Kyiv.

**Storage of anthropological materials:** Institute of Archaeology, National Academy of Sciences of Ukraine.

**Description of samples:**

Three samples were taken for aDNA analysis, two yielded a sufficient amount of DNA for further study.

**UKR045.** *Lehedzyne, burial 16.* Excavated in 2009. Inhumation of a woman aged 25–35 years. The burial pit had a niche, oriented to the north. The rectangular entrance measured 1.7 x 0.65 m. The burial chamber was irregularly oval-shaped, measuring 1.8 x 0.6–1.1 m. The depth of the entrance pit was 1.32–1.40 m, and the depth of the burial chamber was 1.36–1.55 m. The skeleton was lying on the back in an extended position. On the chest, there were 18 carnelian beads and two mother-of-pearl beads. Around the neck, there was a leather cord with six silver and three leather pendants. Fragments of an iron fibula were found above the left clavicle. Chronology according to archaeology: the first to third quarters of the 4th c. CE.

**UKR047.** *Lehedzyne, burial 24.* Excavated in 2009. Inhumation, oriented to the north, partially destroyed. The burial of a woman aged 20–25 years. The pit’s depth was 1 m, and the pit’s contours were not traceable. The skeleton was lying in an extended position, with legs shifted to the east and arms outstretched. Cervical vertebrae and part of the chest were missing, the pelvis was damaged. The ulnas were fractured, and the lower part of the arms was missing. The skull was in place, but the mandible was found in the fill, near the feet. To the west of the skull, there were vessels: a bowl, under which fragments of a one-handed vase, under which fragments of a pot. A fibula lay on the left scapula of the skeleton. Near the left foot, there were a red-clay bowl, a cup, an amphora, and a bead. There were traces of a bronze object (possibly a buckle) on one of the lower vertebrae. Chronology according to archaeology: the second half of the 4th c. CE.

#### **Zolochiv** *(Золочів)*

*(Archaeology - I. Shramko, text – I. Shramko, S. Zadnikov)*

The burial was located on the western outskirts of Zolochiv town, on the edge of the right bank of the Udy River (Siverskyi Donets basin). It was accidentally discovered by local residents near a quarry while extracting sand for household purposes in October 2008. The object has been completely destroyed, but it was possible to ascertain some details of the burial and collect most of the items left in the grave during the burial ceremony. Fragments of pottery vessels were collected, as well as artefacts made of bone, glass, and metal. Only traces of part of the burial chamber were visible. ^195,196^

**Location:** 50.279939 N, 35.957792 E. Chalogo street, Zolochiv city, Bohodukhiv, Kharkiv region.

**Excavations:** 2008.

**Excavation authors:** Iryna Shramko, Museum of Archaeology of V.N. Karazin Kharkiv National University.

**Storage of anthropological materials:** Museum of Archaeology of V.N. Karazin Kharkiv National University.

**Description of samples:**

One sample was taken for aDNA analysis.

**UKR102.** *Zolochiv, ground burial 1.* The burial chamber was in the form of a rectangular pit with a niche. The burial chamber was excavated in sandy soil, measuring 2.10 m in length, and was oriented along an east-west axis, with its bottom situated at a depth of 2.70 m. There was a noticeable trace of a vertical shaft (dromos) filled with mixed soil. The human skeleton was oriented with the head to the west. The accompanying artefacts were located on the skeleton’s right side. A significant portion of the pots and glassware were broken into fragments, but some items remained intact: a large bi-conical-shaped bowl, a small dish, an iron sword, a dart point, a fragment of an iron knife, a bone comb, a product made of tubular bird bone, and a bronze dart. Human bones had been hidden by modern-day grave robbers in a pit near the burial. Chronology according to archaeology: 4th c. CE.

### EARLY MIDDLE AGE

#### Saltiv (Saltiv-Majaki) culture *(Салтівська культура)*

*(Archaeology – V. Borodulin, V. Aksonov, text – V. Aksenov)*

The Saltiv culture got its name from an archaeological site near the village of Verkhnii Saltiv, which is represented by a hillfort, a burial ground, and a series of settlements. The site was found in 1900 by the local educator V.O. Babenko. The excavated artefacts allowed to identify a distinct archaeological Saltiv culture associated with the Khazar Khaganate. The culture dates to the final quarter of the 1st millennium CE (the second half of the 8th to the first half of the 10th c. CE) ^197^.

The sites of Saltiv culture have been found across an extensive area, ranging from the Caucasus Mountains in the south to the Middle Volga region in the north, and from the Volga River in the east to the lowlands of the Dnieper River in the west, primarily encompassing steppe regions. Within this territory, researchers identify local types, including the forest-steppe, the mid-Don steppe, the lower Don steppe, the Azov Sea region, the area of the Caucasus foothills and plains, and the Crimean region^198,199^.

Within this area, diverse types of archaeological sites have been found, including hillforts, settlements, temporary camps, catacombs, pit burials, and kurgans. Hillforts have been found with stone walls as well as wooden and ground fortifications ^73,200–203^. At settlements, excavations have revealed dugouts, semi-dugouts ^198,204–206^, yurt-like dwellings ^207^, household buildings and pits ^202,208–210^, as well as religious complexes ^211^.

Various burial practices have been identified, including burials in catacombs ^201,212–214^, inhumation pits of various types ^215–219^, cremation burials with subsequent placement of human remains and artefacts in pits or urns ^220,221^, and inhumation burials in kurgans ^199^. The diversity of burial customs within the Saltiv culture indicates heterogeneity of population. The following burial practices are associated with specific ethnic groups: inhumations in underground catacombs with Alans, inhumations in pits with Bulgars and mixed Turkic-Ugric population, inhumations in kurgans with Khazars, cremations with Turkic, Slavic, and Adyghe-Abkhaz individuals.

The spread of the Saltiv culture within the forest-steppe and steppe zones led to a diverse range of economic activities, including various forms of agriculture and cattle breeding ^198,199,204,222^.

The presence of weaponry items in the materials from burials, such as sabers, armor-piercing spearheads, battle-axes, bone and horn arrowheads, iron arrowheads, as well as equestrian gear items (bits, stirrups, horse harness decorations), suggests that the Saltiv military predominantly consisted of light cavalry units ^223–225^.

The broad cultural and trade exchanges between the Saltiv culture, Byzantium and the Arab Caliphate are evidenced by the elements of lotus-like belt ornaments ^226^, ornaments crafted in a Crimean-Byzantine style ^227^, the presence of early medieval amphorae and pottery produced in Crimea ^228,229^, Arab silver dirhams and Byzantine gold soliduses ^230^.

#### **Verkhnii Saltiv** *(Верхній Салтів)*

The Verkhnii Saltiv catacomb burial ground was found in 1900 by V.O. Babenko. It is located northward of the Verkhnii Saltiv Hillfort, on the slopes of numerous ravines crossing the high right bank of the Siversky Donez River. From 1900 to 1917, V.O. Babenko excavated three separate sections with catacombs, which he designated as Verkhnii Saltiv I (main), Verkhnii Saltiv II, and Verkhnii Saltiv III burial grounds. An additional burial ground – Verkhnii Saltiv IV – was found in 1989 by the expedition of the Kharkiv Historical Museum led by V.G. Borodulin.

The Verkhnii Saltiv catacomb burial ground I (main) occupies the western and northwestern slopes of the Kapinosova Ravine. Up to 1960, about 417 catacomb burials were excavated there ^221^. It is considered that this section of the burial ground was used throughout the whole culture period, spanning from the second half of the 8th to the middle of the 10th c. CE ^231^. In 1984, excavations at the burial ground were initiated by an expedition from the Kharkiv Historical Museum under the leadership of V.G. Borodulin. From 1984 to 1989, the expedition studied 75 catacombs, one burial in a dromos, 16 burials in pits of various types, and 4 burials of horses in separate pits. These researched burial complexes are dated to the middle of the 8th–9th c. CE ^232,233^.

The Verkhnii Saltiv catacomb burial ground III is located on the eastern slope of the right bank of the Siversky Donez River northward of the settlement, beyond the ravine. The cemetery was found in 1902 by V.O. Babenko, who excavated 4 catacomb burials in 1902–1903. In 1959–1961, excavation was carried out by the Institute of Archaeology (Kyiv) led by D.T. Berezovets. During this expedition, 17 catacomb burials and 3 separate burials with horse remains were studied. The Kharkiv Historical Museum expedition under the leadership of V.G. Borodulin, conducted research in 1988–1992. The work resulted in the excavation of 25 catacombs, 5 pit burials, 4 horse burials in separate pits, and 1 burial in a dromos ^221^. Based on coins and elements of a belt set, the burial ground is believed to have originated in the second half of the 8th c. CE and was in use throughout the 9th c. CE ^231,232^.

The Verkhnii Saltiv catacomb burial ground IV is located on the eastern slope of the Netechinsky ravine and forms a single area with the Verkhnii Saltiv II burial ground ^234^. In 1989–1990 17 catacombs were explored by an expedition of the Kharkiv Historical Museum led by V.G. Borodulin. An additional 10 catacombs were excavated in 1996 by an expedition of G.S. Skovoroda Kharkiv State Pedagogical University led by V.V. Koloda. From 1998 to 2021, the burial ground was explored by the expedition of the Kharkiv Historical Museum led by V.S. Aksenov. To date, the total number of studied burial complexes is 164 plus 1 burial of a horse. Most of the studied complexes are dated to the 9th c. CE.

**Location:** 50.135123 N, 36.799162 E. Verkhnii Saltiv village, Chuhuev district, Kharkiv region, Ukraine.

**Excavations:** 1986, 1989, 2014.

**Excavation authors:** Vyacheslav Borodulin (1986, 1989), Viktor Aksonov (2014), M. F. Sumtsov Kharkiv Historical Museum, Kharkiv, Ukraine.

**Storage of anthropological materials:** M. F. Sumtsov Kharkiv Historical Museum, Kharkiv, Ukraine

**Description of samples:**

One sample was taken from burial ground I, one from burial ground III and four from burial ground IV. Individuals were referred to as Alans based on the catacomb type of burials.

**UKR134.** *Verkhnii Saltiv catacomb burial ground І, catacomb 42.* Excavated in 1986. T-shaped catacomb with a floor size of 2.25 x 1.45 m. On the floor, the remains of three individuals were found: two women and a child. The skeletal remains showed deliberate damage, which had occurred in ancient times during a ritual neutralization of the deceased people. One female burial (UKR134) was accompanied by a bronze trapezoid-shaped pendant, a bronze wire bracelet, and a bronze ring. The other female burial contained bronze wire decorations, bronze bells, plate-shaped trapezoid pendants, a glass necklace, and silver earrings. The child’s remains were accompanied by glass beads. Chronology according to archaeology: second half of 8th–9th c. CE.

**UKR135.** *Verkhnii Saltiv catacomb burial ground III, catacomb 11.* Excavated in 1989. The chamber was oriented longitudinally in relation to the dromos, had a size of 2.4 x 1.6 m. The remains of two individuals, a man and a woman, were found on the floor. The skeletons exhibited deliberate damage, which had occurred in ancient times during a ritual neutralization of deceased people. The man (UKR135) was accompanied by the following artefacts: iron battle-hammer, an iron battle-axe, silver decorations from footwear straps, a bronze belt buckle, an iron knife, and a bronze pendant-seal. The female burial was accompanied by bronze earrings, a glass necklace, a bronze mirror, a bronze medallion, 3 bronze wire bracelets, 2 rings, a bronze toiletry case, a bronze belt buckle and a tip, silver decorations from shoe straps. Chronology according to archaeology: second half of 8th–9th c. CE.

**UKR136.** *Verkhnii Saltiv catacomb burial ground, catacomb 120.* Excavated in 2014. T-shaped catacomb with a floor size of 2.3 x 1.83 m. On the floor, the remains of three individuals (two men and one woman) were found. In the wall of the dromos, a niche had been made, where the remains of a child aged 1–2 years were found, along with a bronze wire bracelet, a bronze bell, and a bronze stamped button. The burial of one man (UKR136) was accompanied by a bronze ring and bronze elements of a belt. The artefacts associated with the other man consisted of bronze elements of a belt and metal elements from shoe straps. On the woman’s remains, bronze earrings, a bronze ring, and a glass necklace were found. Chronology according to archaeology: second half of 8th–9th c. CE.

**UKR137.** *Verkhnii Saltiv catacomb burial ground, catacomb 122.* Excavated in 2014. T-shaped catacomb with a floor size of 1.8 x 0.9 m. On the floor, the remains of two individuals (a man and a child) were found. The man’s (UKR137) burial was accompanied by three iron knives, three cast bronze bells, an iron fibula, iron belt buckle, iron tweezers, and one glass bead. The child’s burial was accompanied by a hand-formed clay pot. Chronology according to archaeology: second half of 8th–9^th^ c. CE.

**UKR138.** *Verkhnii Saltiv catacomb burial ground, catacomb 124.* Excavated in 2014. The catacomb was incomplete and consisted only of a dromos, where the remains of an adult man were found, accompanied by artefacts: a bronze belt buckle, a bronze belt tip, an iron knife, an iron adze-hoe, and a reliquary made of animal bone. Chronology according to archaeology: second half of the 8th–9th c. CE.

**UKR139.** *Verkhnii Saltiv catacomb burial ground, catacomb 125.* Excavated in 2014. T-shaped catacomb with a floor size of 1.82 x 1.42 m. On the floor, the remains of an adult woman were found, accompanied by bronze butterfly-shaped decorations, two bronze chumbar blocks, two iron knives, a bronze duck-shaped pendant-amulet, a glass bead necklace, bronze wire decorations, and a bronze bell. Chronology according to archaeology: second half of 8th–9th c. CE. Chronology according to ^14^C dating: 671–874 cal CE (1256±27BP).

#### **Bochkove** *(Бочкове)*

*(Text, archaeology – Oleksiy Laptev)*

The burial ground was found 1.3 km west of the modern western outskirts of the village of Bochkove. The burial ground occupies a kurgan-like elevation on the right bank of the Vovcha River, located 23 km from its confluence with the Siversky Donez River. ^235,236^

**Location:** 50.314768 N, 37.11063 E. Bochkove village, Chuhuev district, Kharkiv region.

**Excavations:** 2014.

**Excavation author:** Oleksii Laptev, M. F. Sumtsov Kharkiv Historical Museum, Kharkiv, Ukraine. **Storage of anthropological materials:** M. F. Sumtsov Kharkiv Historical Museum, Kharkiv, Ukraine. **Description of samples:**

Eight samples were taken, DNA was successfully extracted from four and three of them yielded a sufficient amount of DNA for further study. Individuals were referred to as Bulgars based on the pit type of burials.

**UKR143.** *Bochkove, burial 3.* The rectangular-shaped burial pit was oriented along the east-west axis. The filling of the pit was stratified: in its upper part, there was marl, and in the lower part, humus. The burial pit had a circular widening: at a distance of 0.6–0.7 m from the surface, it expanded in all directions. The buried adult woman lay in a wooden coffin, curled up on the right side with her head to the west. The bone preservation was poor and the skull was heavily decomposed. The neck was unnaturally curved: the first cervical vertebrae were oriented along the ’west-east’ axis and those adjacent to the thoracic region were oriented along the ’north-south’ axis. It is possible to speculate that such a fracture occurred as a result of arranging the required ritual position after death. The burial contained modest but interesting items: two earrings under the skull, a quarter of a dirhem in front of the jaw, a second quarter of a dirhem (which could be combined with the first to form half a coin) in the abdominal area under the left forearm. In the feet area, in the northwest corner, there was a mug.

**UKR144.** *Bochkove, burial 4.* An inhumation in a ledge-pit with a niche. The filling of the burial pit consisted of marl in the upper part and humus in the lower part (excavated in compact marl). The grave-pit had dimensions of 2.21×0.84 m, the depth was 1.65–1.8 m. The remains of a woman (height 1.45 m) lay stretched out on the back with the head to the north. The right arm was extended along the body with the palm facing down. The left arm was bent at the elbow, with the hand resting on the chest. The bone preservation was poor. Behind the head, there was a bowl with the handle broken in ancient times, and a thin metal mirror. A quarter of an Arabic dirhem coin was found in the mouth, whereas a one-eighth part of the coin was found on the lumbar vertebra. Two cast buttons were found on the collarbones, and a ring with a glass insert was found on a right hand finger. Chronology according to archaeology: 9th с. CE. Chronology according ^14^C dating: 671–874 cal CE (1256±27BP).

**UKR147.** *Bochkove, burial 6.* An inhumation in a ledge-pit with a niche. The bottom of the burial pit was found at a depth of 1.6–1.65 m. The burial contained the skeleton of an adult male (height 1.55 m), lying stretched out on his back with his head to the northwest. The bone preservation was good. Foot bones were absent in situ, but were found in the filling above the left elbow. The left leg was turned outward. The right arm lay along the body with the palm facing up. Beneath the right forearm, a long iron knife in a wooden sheath and an iron awl with remains of a wooden handle and sheath were found.

There were no other burial goods. Chronology according to archaeology: 9th с. CE. Chronology according ^14^C dating: 683–883 cal CE (1232±27BP).

### MIDDLE AGES

#### Сumans, Cuman culture *(Половці)*

*(Archaeology, text - S. Andrukh, G. Toshev)*

Cumans (Polovtsi, Kipchaks) were a medieval nomadic Turkic-speaking people. During the 11th to 14th c. CE, Cumans inhabited the Eurasian steppes from the Irtysh River to the Danube River. As they spread, they displaced the Pechenegs and Khazars. Later, in the 13th c. CE, became a part of the Mongol Empire. We know about Cumans from written sources and archaeological materials. Judging by these, Cumans had close military, political, cultural, and trade contacts with their neighbours, including Byzantium, Kievan Rus, Volga Bulgaria, and others. Their main occupation was cattle breeding. Cumans became a component of the ethnogenesis of Kipchak-speaking peoples, such as the Tatars, Karaites, Nogais, etc.

In the mid-11th c. CE Cumans appeared in the Northern Black Sea region ^237^. In the southern steppes of the Northern Black Sea region, information about Cumans is primarily known from archaeological data. For burials they used both kurgans and ground graves. The most famous complex is a leader’s burial in the Chingulsky Kurgan (Zaporizhzhia region). The ordinary people were buried in pits, placing bodies stretched out on their back accompanied by funerary goods, including weapons, everyday objects, jewellery, etc. To the north, in the basin of the Siversky Donets River, the Cuman cities of Balin, Suhrov, and Sharukan are known, which served as settlements for the Cuman khans and were inhabited by a settled agricultural population of Alans. The Alans in these towns engaged in craft production and agriculture for the needs of the Cuman nomads.

Written and archaeological sources indicate that Cuman horde confederations in the 12th c. CE were multi-ethnic. In addition to the Cumans, these unions brought together various steppe and forest-steppe populations that lived in the region before Cuman arrival. These included Alans and Bulgars from the forest-steppe zone of the Siversky Donets basin, Pechenegs, and Torks from steppe regions who were completely subjugated by Cumans. In the 12th c. CE Cumans underwent a process of forming an ethnic community in which Cumans themselves were the political core around which all other ethnic groups united. The Cumans gave this community its ethnic name. The unity of this political community was primarily determined by a common language, and unified cultural traditions including material culture, funerary rites, religious beliefs, shared epic narratives, and songs ^238^.

Аccording to anthropological data, Cumans, who initially had a distinct South Siberian anthropological type, quickly lost it, dissolving into the larger population of the Northern Black Sea steppes ^238^. During their first 50 years in the Northern Pontic region, Cumans, like preceding Pechenegs, obtained significant profit from the local Slavic agricultural population through simple plunder, often accompanied by the capture of prisoners for their sale on the slave markets of the Crimea. In the early 13th c. CE, the situation changed, and relative peace and military balance were established between the Cumans and the Slavs. The prolonged coexistence led not only to mutual raids but also to joint allied campaigns and marriages. The nomads actively absorbed the culture of the settled agricultural population, including aspects of daily life, elements of clothing, and household items. The long-term coexistence of Slavs and Cumans resulted in bilingualism among the local population. This process was interrupted by the Tatar-Mongol invasion (Golden Horde). ^239^

#### **Mamay-Gora** (Мамай-Гора) (Archaeology, text – S. Andrukh, G. Toshev)

The archaeological site has been described under the Neolithic Azov-Dnieper culture section.

On the territory of Mamay-Gora, over 60 Cuman burial complexes have been identified. According to the artefacts and coin finds, the ground and kurgan complexes of Mamay-Gora fit into the framework of the 12th–first third of 15th c. CE ^82,240^.

**Location:** 47.432845 N, 34.27259 E. Mamay-Gora tract, Velyka Znamianka village, Vasylivsky District, Zaporizhzhia Region.

**Excavations:** 1997.

**Excavation authors:** Svitlana I. Andrukh, Gennadi M. Toshev, Zaporizhzhia National University, Zaporizhzhia, Ukraine.

**Storage of anthropological materials:** Zaporizhzhia National University, Zaporizhzhia, Ukraine.

**Description of samples:**

One sample was taken from a Cuman ground burial.

**UKR012.** *Mamay-Gora, kurgan 162, burial 13.* Excavated in 1997. Inlet burial without artefacts. The depth was 1.35 m. The burial was made in a pit with a niche. The entrance pit was covered with planks and had an elongated oval shape, measuring 2.4 m in length. It was oriented along a north-south line. Along the eastern sloping wall, at a depth of 0.3–0.4 m from the contour fixation level, a step with a width of 0.35 m was observed, sloping down towards the chamber. The chamber, measuring 2.25×0.7 m, was located on the western side, elongated along the south-north line. The depth from the contour fixation level was 0.7 m. The vault was destroyed in ancient times. The skeleton lay stretched out on its back with the head oriented to the north. A ram bone was found behind the skull. Initially, the burial was dated to the Scythian period, but later the time was defined as medieval. ^80^.

#### **Velyka Znamianka** (Велика Знам’янка) (Archaeology, text – S. Andrukh, G. Toshev)

Kurgan group II, consisting of four kurgans, is located on the western outskirts of the village of Velyka Znamenka, 0.2 km south of the Zaporizhia-Kherson highway. These kurgans were excavated in connection with the irrigation system reconstruction project in 1996 by an archaeological expedition from Zaporizhzhia State University. The excavated kurgans contained 30 burials from different time periods. In kurgan 18, measuring 0.45 m in height and 19 m in diameter, seven burials were identified (including two Scythian and five medieval burials). ^241^

**Location:** 47.439358 N, 34.335757E. Velyka Znamianka village, Vasylivsky District, Zaporizhzhia Region.

**Excavations:** 1996.

**Excavation authors:** Svitlana I. Andrukh, Gennadi M. Toshev, Zaporizhzhia National University, Zaporizhzhia, Ukraine.

**Storage of anthropological materials:** Zaporizhzhia National University, Zaporizhzhia, Ukraine.

**Description of samples:**

One sample was taken from a Cuman period burial.

**UKR027.** *Velyka Znamianka, kurgan 18, burial 1.* Inlet burial. Excavated in 1996. The burial was robbed and completely destroyed during the robbery. It was located at a depth of 0.7 m. The pit had a rectangular shape, measuring 2.35×1.1 m, elongated in a west-east direction. In the southeast part of the pit, the remains of a wooden coffin were preserved. Above the burial, at a depth of 0.05–0.15 m, a horse’s skull was found, at the base of which was a bone buckle and fragments of reins. At the same depth, small fragments of an iron sword were found. There were two bone clusters near the southern wall of the pit. The eastern cluster contained a skull, ulnas and radiuses, pelvis, femurs, and vertebrae. In the western cluster, located 0.4 m away, tibia bones were arranged. Many small fragments of a sword were found on the bones. To the north of the skull, a silver plaque was found. Under the western cluster, gold foil, two bone inlays, two bronze buttons, fragments of a sword, a fragment of an item with rivets, a fragment of a ring, three iron arrowheads in a decayed leather case, and a bone cheek-piece were found. Chronology according to archaeology: 12th c. CE.

#### **Kumy** (Куми)

*(Archeology – I. Shramko; text – I. Shramko, S. Zadnikov)*

The archaeological site has been described under the Early Iron Age Cimmerian culture section.

The kurgan was found in 2010 during a scientific archaeological expertise ^107^. The site where the kurgans were located is situated in the fields of the Krasnograd Research Station, near Kumy village in the Krasnograd District of the Kharkiv Region. The kurgan, located on the edge of a watershed plateau on the right bank of the Berestovaya River (right tributary of the Orel River), consists of nine kurgans of varying sizes, most of which were plowed and are scarcely visible in the field. ^109^

**Location:** 49.3246500 N, 35.3687167 E. Kumy village, Krasnograd district, Kharkiv region, Ukraine.

**Excavations:** 2010.

**Excavation authors:** Iryna Shramko, Museum of Archaeology of V.N. Karazin Kharkiv National University, Kharkiv, Ukraine.

**Storage of anthropological materials:** Museum of Archaeology of V.N. Karazin Kharkiv National University, Kharkiv, Ukraine.

**Description of samples:**

One sample was taken from a Cuman ground burial.

**UKR056.** *Kumy, kurgan 1, burial 1b.* A damaged complex containing the remains of two destroyed burials dated to the Late Bronze Age (2nd period of the Berezhnevo-Mayivska Zrubna Culture) (1a) and the Middle Ages (1b) (the inlet burial). It is impossible to determine the sizes of the burial pits and the level of their bottom because these complexes have been disturbed by burrowing animals. The human bones and burial goods were displaced from their original places and were found in mixed soil layers. Fragments of Bronze Age ceramics were found among the artefacts from the late Middle Ages. It was difficult to establish the orientation of the burials and the skeletons. An iron knife, remains of heavily corroded iron stirrups, and a fragment of a bone plate date to the Middle Ages. In the southern sector of the pit, at a depth of 0.95 m, a human skull (without the lower jaw) and clavicle bones were found. Chronology according to archaeology: 9th–11th c. CE. Chronology according to ^14^C dating: 991–1149 cal CE (1010±25 BP).

#### Kyivan Rus and Golden Horde: Slavs and eastern nomads *(Київська Русь і Золота Орда)*

*(Text – V. S. Tersky, S. Zadnikov)*

Starting from the 8th c. CE, Slavic tribes in Eastern Europe began to unite into political alliances, and in the 9th c. CE the state of Kyivan Rus was formed, with its center in the city of Kyiv, which emerged in the 6th c. CE. From the second half of the 11th c. CE, Kyivan Rus experienced a period of feudal fragmentation. The state disintegrated into separate principalities – independent state structures with their own elites, trade and political relations with neighbours, and other local features that determined the direction of their historical development.

From the mid-8th to the last third of the 10th c. CE, Slavic tribes in the southeast were economically dependent on the Khazars, carriers of the Saltiv culture. From the second half of the 13th to the last quarter/end of the 14th c. CE, the development of Slavic statehood was complicated by the presence of eastern nomads, Turks and Mongols (Golden Horde).

The main Slavic burial rite until the 9th c. CE was cremation. However, after the adoption of Christianity, cremation gradually gave way to the rite of inhumation. In the 9th–10th c. CE, a kurgan culture known as the “warrior culture” emerged, belonging to representatives of the Slavic military-political elite. Anthropological material from this period is very poor.

Burials of Golden Horde representatives in the forest-steppe were usually found accidentally and typically were without artefacts, making their identification problematic. Large separate burial grounds in the forest-steppe have not been found yet. In the steppe, the most well-known landmark is Mamay-Surka. Many burials have been uncovered in Mamay-Gora, as well as in other steppe sites ^242^.

#### **Zvenyhorod** (Звенигород)

*(Archaeology – H. Vlasova, V. Shelomentsev-Tersky, text – S. Tersky)*

The landmark of Zvenyhorod is related to the Slavic culture of the Kingdom of Galicia-Volhynia (Kingdom of Ruthenia), which emerged as one of political centres after the collapse of Kyivan Rus and included western Ukraine with some areas in Belarus, Poland, Moldova, and Lithuania.

The Galician group of landmarks is represented by cities, fortified and unfortified settlements, and inhumation burials. From the 10th to the 12th c. CE, kurgans practically were not used in the burial rite of Eastern Carpathians. The main type of burial is ground pits, and occasionally stone sarcophagi (predominantly in princely churches). Among ground burials, there were ones under stone plates, which indicates more southern customs. The traditional skeletal position is supine, with the head oriented mainly westward.

Zvenyhorod as a city with a princely court arose at the end of the 11th c. CE from several local unfortified settlements of 10–11 c. CE. The city is located near the Main European Watershed, which is conventionally considered to be the border between Eastern (Carpathian) Croats and Volhynians (Buzhans). These Slavic tribes settled the basins of the upper Western Bug River and the upper Dniester River in early medieval times. Therefore, the local population in the 12th and 13th c. CE could have descendants of these two tribal groups.

The Zvenyhorod fortifications covered approximately 50 hectares. Within this area, the remains of three churches and nine cemeteries have been studied, and another seven cemeteries were found within a radius of up to 2 km. A significant agglomeration developed around the city, extending 15 km from west to east and at least 7 km from north to south. Until 1241, this area may have housed at least 12 churches and five monasteries, each with their mainly small cemeteries. The two large (city-wide) cemeteries of the 11th–12th c. CE, located at Mount Hoyeva Hora and Zahumenki, have been investigated most thoroughly. All the burials found in Zvenyhorod are inhumations in grave pits. ^243–245^

**Location:** 49.73363 N, 24.247629 E. Zvenyhorod village, Lviv district, Lviv region.

**Excavations:** 1960, 1962.

**Excavation authors:** Halina Vlasova, Volodymyr S. Shelomentsev-Tersky, Lviv Historical Museum, Lviv, Ukraine.

**Storage of anthropological materials:** Lviv Historical Museum, Lviv, Ukraine.

**Description of samples:**

One sample was taken for aDNA analysis.

**UKR166.** *Squares 12 and 13 К (№ 23353), cultural layer I-II, Excavation No. 1.* The burial was found at the depth of 30 cm in the northwestern part of Okolny town. The burial was associated with the Slavic population on the border between Volynians and White Chroats. Chronology according to archaeology: the first half of the 13th c. CE.

#### **Donets hillfort** (Донецьке городище)

*(Archaeology – B. Shramko, text – I. Shramko, S. Zadnikov)*

The Donets hillfort is related to the Slavic culture of Kyivan Rus in the Siversky Donets basin in the Golden Horde period (13th–14th c. CE). The territory of the upper Siversky Donets River was on the border with nomadic tribes, which influenced the population’s lifestyle and the development of settlement infrastructure. Archaeological excavations revealed dwellings, workshops, farm buildings and numerous artefacts from two periods of the settlement’s existence. During the early Middle Ages (8th–10th c. CE), fortified settlements of early Slavs emerged here, later evolving into cities of the Kyivan Rus (10th–13th c. CE).

In the 11th c. CE, the territory belonged to the Principality of Pereyaslavl. From the 10th to the 13th c. CE, the city played an important military-defensive role on the southern border of the ancient Rus state and had significance for international trade along ancient overland and water routes along the Donets and Don rivers ^246^. In 1239, during the Tatar-Mongol invasion, the city, abandoned by residents, was burned ^246^.

During the Mongol invasion, the city was abandoned by its population. Some parts of the city were used as a burial ground, but no cultural artefacts have been documented from the second half of the 13th– 14th c. CE. On the territory of the settlement, burials dated to this period were found ^247,248^. At that time the southeastern territories of the state fell under the rule of the Golden Horde.

**Location:** 49,921928N, 36,192856 E. Kharkiv city, Novobavarsky district.

**Excavations:** 1960.

**Excavation authors:** Borys A. Shramko, V. N. Karazin Kharkiv National University.

**Storage of anthropological materials:** Museum of Archaeology of V.N. Karazin Kharkiv National University.

**Description of samples:**

Five samples from excavation 4 of the Donets hillfort belonging to the Golden Horde period were taken. Three samples yielded a sufficient amount of DNA for analysis.

**UKR068.** *Donets hillfort. Вurial 1, field number 1132/IV-1960*. The burial was found at a depth of 0.4–0.45 m. The pit had a rectangular shape, with a length of 1.7–2.0 m and a width of 0.55–0.6 m. Remains of wood were found along the wall. In the burial, there was a skeleton that was osteologically estimated to be of a middle-aged man (but genetically female). The skeleton laid out on the back with the head oriented to the west. The right arm was placed on the chest, the left on the abdomen. There were no artefacts. Chronology according to archaeology: the end of 13th–beginning of 14th c. CE.

**UKR069.** *Donets hillfort, Вurial 4.* The burial was found at a depth of 0.4–0.45 m. The pit had a rectangular shape, with a length of 1.7–2.0 m and a width of 0.55–0.6 m. An elderly man was buried, laid out in an extended position on the back, with the head oriented to the west. There were visible traces of two strong blows on the skull. The right arm was placed on the chest, the left on the abdomen. There were no artefacts. Chronology according to archaeology: the end of 13th–beginning of 14th c. CE.

**UKR070.** *Donets hillfort, Вurial 2, field number 1150/IV-1960.* The burial was found at a depth of 0.4–0.45 m. The pit had a rectangular shape, with a length of 1.7–2.0 m and a width of 0.55–0.6 m. A very elderly individual with toothless jaws, where tooth sockets in the gums long healed, was buried. The skeleton lay on the back, with the head oriented to the west. The right hand was on the chest, the left on the abdomen. No artefacts were found. The burial pit was located above the household pit No. 81 with filling from the period of Kyivan Rus. Chronology according to archaeology: the end of 13th–beginning of 14th c. CE.

#### **Kumy** (Куми)

*(Archaeology, text – I. Shramko, S. Zadnikov)*

The archaeological site has been described under the Early Iron Age Cimmerian culture section.

Historians associate the decline of life in certain ancient Rus settlements to the time of the Mongol Golden Horde invasion ^246,248^. The buffer zone of the Golden Horde in the forest-steppe of the Dnieper left bank covered an area of about 200 km in the meridional direction, and in the Siversky Donets basin, its northern border ran somewhat south of Kharkiv ^249^. Numerous monuments from the post-Mongol period were found in this zone ^109,250–252^. Materials from the Kumy site significantly expand our understanding of the life of the population in the middle reaches of the Siversky Donets in the post-Golden Horde period, challenging the notion of a long existence of an uninhabited “Wild Field” in this territory. It is more likely that cohabitation occurred in the post-Golden Horde period within the contact territory of both sedentary Slavic farmers and steppe nomads.

**Location:** 49.3246500 N, 35.3687167 E. Kumy village, Krasnograd district, Kharkiv region, Ukraine.

**Excavations:** 2010.

**Excavation authors:** Iryna Shramko, Museum of Archaeology of V.N. Karazin Kharkiv National University.

**Storage of anthropological materials:** Museum of Archaeology of V.N. Karazin Kharkiv National University.

**Description of the samples:**

One sample was taken from Kumy ground burial of the Golden Horde period.

**UKR063.** *Kumy, kurgan 1, burial 3.* An inlet burial attributed to medieval Golden Horde nomads. The burial pit was found at a depth of 0.80 m, at the level of loam. The pit had a rectangular shape with rounded corners, and its longer axis was oriented in the west-east line. There was a small ledge in the southern part. The skeleton of a child was found, which lay on the back in an extended position with the head facing west. The right arm was extended along the body, the left arm was slightly bent at the elbow, likely the hand was resting on the pelvic bones. The legs were extended and brought together at the feet. No artifacts were found in the burial. Chronology according to archaeology: 14th c. CE.

#### **Ploske** (Плоске)

*(Archaeology - V. Mikheev, text – I. Shramko, S. Zadnikov)*

The ground burial near Ploske village belonging to the settlement of the Saltiv culture is placed on the elevated right bank of the Siversky Donets River in the immediate vicinity of the Mayaki hillfort. The Mayaki hillfort was a fortified settlement of the early medieval period (9th–10th c. CE). On the edge of the settlement, burial ground No. 4 of the Golden Horde period was explored ^205^. The ground burial 9 was found in excavation № 9, in the eastern part of the settlement. According to archaeological data, it was preliminarily dated to the 13th–14th c. CE ^253^.

**Excavation authors:** Volodymir K. Mikheev, V. N. Karazin Kharkiv National University.

**Excavations:** 1965.

**Storage of anthropological materials:** Museum of Archaeology of V.N. Karazin Kharkiv National University, Kharkiv, Ukraine.

**Location:** 48.940698 N, 37.666706 E. Mayaki village, Kramatorsk district, Donetsk region.

**Description of samples:**

One sample was taken for aDNA analysis.

**UKR074.** *Ploske, Burial 9, excavation 9.* The pit was found at a depth of 0.85 m. The human skeleton was badly damaged. It lay on the back in an extended position, with the head oriented to the west. To the right of the skull, there was a fragment of a thick-walled clay vessel. To the left side of the skull, a conical bone amulet with two holes for hanging was found. ^253^

#### **Mamay-Surka** (Мамай-Сурка) (Archaeology – G. Toshev, S. Andrukh)

The burial site of Mamay-Surka is located on the high left bank of the Kakhovka Reservoir, west of the village of Velyka Znamianka in Vasylivka District, Zaporizhia Oblast, Ukraine. It has been under excavation since 1989 by an archaeological expedition from Zaporizhzhia National University. By 2006, 1,162 burials of the 14th c. CE had been found there, within a total area of 7,166 m^2^. The individuals were buried in pits, stretched out on their backs with the heads oriented to the west. The burial artefacts consisted of jewellery, household items, and rare findings of coins. According to anthropology, the site represents a mixed settled population of post-Cuman time ^254–256^.

**Location:** 47.430345N, 34.280355E. Ground burial Mamay-Surka 14th c. near the Velyka Znamianka village, Vasylivsky District, Zaporizhzhia Region.

**Excavations:** 1997.

**Excavation authors:** Svitlana I. Andrukh, Gennadi M. Toshev, Zaporizhzhia National University, Zaporizhzhia, Ukraine.

**Storage of anthropological materials:** Zaporizhzhia National University, Zaporizhzhia, Ukraine.

**Description of samples:**

Two samples were taken for aDNA analysis, one of them yielded enough DNA to be included in subsequent analysis.

**UKR028.** *Mamay-Surka, burial 188*. The burial was found at a depth of 1.40 m. The contours of the pit were not discernible. Th filling material was black soil. The skeleton of a woman aged 30–35 years lay stretched out on the back witeh the head oriented to the southwest. The arms were bent at the elbows at a right angle, and the hands lay on the pelvis. Between the right elbow and the ribs, the bones of a child were found. On the left side near the temple bone, an earring was found. Among the bones, an iron needle was found. Chronology according to archaeology: 14th c. CE.

#### **Nogai** (Ногаї, ногайцї)

*(Archaeology, text – S. Andrukh, G. Toshev)*

The Nogai Horde was the successor of the Mangit Yurt. It arose on the territory of modern northern Kazakhstan at the end of the 14th c. and the beginning of the 15th c. CE amid the disintegration of the Golden Horde. It was formed by late nomads from the Lower Volga and the Ural regions. The first waves of Mangit people in the Northern Black Sea region are documented at the beginning of the 15^th^c. CE. However, at that time, they did not have significant ethno-political power.

In the steppe regions of the Northern Black Sea, in the early stages, only Nogai kurgans are known, such as the Balkivsky Kurgan, Berdiansky Kurgan, Mamay-Gora burial ground. The Mamay-Gora burial ground had around 200 burials. The individuals were placed in pits or ledge-pits, stretched out on their backs with their heads oriented to the west. The findings of coins in these burials are a basis for dating this largest burial ground in the Northern Black Sea region to the second half of the 15th c. CE.

The situation changes from the mid-16th c. CE when a significant number of Nogais migrated to the Northern Black Sea region. They form autonomous hordes, including the Yedisanska, Yedichkulska, Jambuylutska, and Budzhakska ^33,257,258^.

**Mamay-Gora (Mamai-Hora)** (Мамай-Гора) (Archaeology, text – G. Toshev, S. Andrukh)

The archaeological site has been described under the Neolithic Azov-Dnieper culture section.

**Location:** 47.432845 N, 34.27259 E. Mamay-Gora tract, Velyka Znamianka village, Vasylivsky District, Zaporizhzhia Region.

**Excavations:** 2006**–**2008, 2012–2013.

**Excavation authors:** Svitlana I. Andrukh, Gennadi M. Toshev, Zaporizhzhia National University, Zaporizhzhia, Ukraine.

**Storage of anthropological materials:** Zaporizhzhia National University, Zaporizhzhia, Ukraine.

**Description of samples:**

Nine samples affiliated to Nogais were taken from Mamay-Gora ^242,257,258^, DNA was extracted from eight and seven of them yielded enough DNA to be included in subsequent analysis.

**UKR016.** *Mamay Gora, object 242, burial 1.* Excavated in 2012. The burial pit was found at a depth of 0.65 m. It had an oval shape, measuring 2.5×0.6 m, with a depth of 0.45 m. The pit was filled with black soil. The skeleton was lying stretched out on the back with the head oriented to the west. The right arm was placed along the body, the left arm was bent at the elbow with the hand on the pelvis. ^259^. Chronology according to archaeology: 15th c. CE.

**UKR017.** *Mamay Gora, object 266, burial 1.* Excavated in 2013. Found at a depth of 0.9 m. The pit had an elongated oval shape (2.25×0.7×0.5 m) and a depth of 0.4 m. It was filled with black soil mixed with clay. The skeleton of a 50–55-year-old man was lying stretched out on his back with the head oriented to the west (slightly turned to the south). The arms were placed along the body. ^259^. Chronology according to archaeology: 15th c. CE.

**UKR018.** *Mamay Gora, object 206, burial 1.* Excavated in 2008. The rectangular shaped pit was found at a depth of 0.85 m and measured 2.3×0.85 m. The filling was mixed. In the northern part of the pit, remains of wood were found. Along the southern wall, at a depth of 0.2 m from the contour of the pit, there was a step about 0.25 m wide. The total depth of the pit was 0.4 m. The burial chamber had a size of 2.3×0.6 m. The skeleton of an adult individual lay stretched out on the back with the arms placed along the body. The knees were slightly bent to the right. ^260^. Chronology according to archaeology: 15th c. CE.

**UKR020.** *Mamay Gora, object 260, burial 1.* Excavated in 2013. Found at a depth of 0.9 m. The pit had an elongated oval shape, measuring 2.3×1×0.8 m, with a depth of 0.7 m. The pit filling was mixed. The skeleton of a 40–45-year-old man lay stretched out on the back with the head oriented to the west (slightly turned to the south). The right arm was bent at the elbow, with the hand on the pelvis, the left arm was placed along the body. ^259^. Chronology according to archaeology: 15th c. CE.

**UKR021.** *Mamay Gora, object 268, burial 1.* Excavated in 2013. The burial was found at a depth of 1.05 m and was made in a pit with a niche. The entrance pit measured 2.1×0.6 m and had a depth of 0.1 m from the contour fixation level. The filling of the pit was mixed. Below the level of the entrance pit, there was an oval-shaped burial chamber measuring 2.2×0.6 m. The skeleton of a 45–50-year-old woman was lying stretched out on the back with the head oriented to the east (slightly turned to the south). The left arm was bent at the elbow at a right angle, with the hand on the pelvis, while the right arm was placed along the body. ^259^. Chronology according to archaeology: 15th c. CE.

**UKR022.** *Mamay Gora, object 194, burial 1*. Excavated in 2007 The pit was found at a depth of 0.85m. It had an almost rectangular shape, with a length of 2.15 m and a width of 0.75 m. At a depth of 0.2 m from the pit contour, along the northern wall, a step up to 0.18 m wide was found, and in the southern wall, there was a niche up to 0.1 m wide. The bottom of the burial chamber was 0.08 m below the step. The skeleton of a woman lay stretched out on the back with the head oriented to the west. The arms were placed along the body. The skull was slightly turned with the facial part to the left. There were wooden remains under the skeleton. ^259^. Chronology according to archaeology: 15th c. CE.

**UKR024.** *Mamay Gora. object 189, burial 1.* Excavated in 2006. The burial outline was found at a depth of 0.84 m. It was made in a ledge-pit. The upper pit had an oval shape, measuring 2.1×0.9 m, with lateral pits that were 0.2–0.3 m wide and 8 cm high. The lower pit was also oval and measured 2.14×0.43 m with a depth of 0.28 m. The skeleton was lying stretched out on its back with the head oriented to the west. The arms were placed along the body. ^259^. Chronology according to archaeology: 15th c. CE.

### EARLY MODERN PERIOD

#### Ukrainian Cossack Culture, lower Desna, left bank of Dniprо, Slavs *(Українське козацтво*

*Лівобережжя, нижня Десна)*

*(Archaeology – V. Skorokhod, text – V. Skorokhod, Y. Sytyi, V. Zhyhola)*

The 17th and 18th c. CE in Ukraine can be considered as the early modern period. Permanent settlements were represented by fortresses and open settlements. In the 18th c. CE, fortresses were built in the Dutch manner, with bastions located outside of the fortifications’ line. The urban constructions on the Left Bank of Dnipro in the 17th c. CE were primarily wooden, but in the 18th c. CE brick buildings appeared next to wooden ones. Traditional dwellings in towns were represented by houses with non-residential basements and the above-ground living floor, which was heated with tiled stoves.

The future resident received a building site within the fortress under the condition of defending the fortifications and participating in military campaigns with own weapons and means of transportation. In addition, residents received land outside the city for cultivating to earn income for living. Taxes were paid from these profits. Both peasants and townspeople were engaged in agriculture, and various crafts were well developed.

As in any city, various ethnic groups lived together. The main population consisted of Ukrainians, but there were also Poles, Germans, people from the Turkish steppe, Russians, Belarusians, Lithuanians, and others. ^261^

#### **Vypovziv** (Виповзів)

Vypovziv Hillfort (village of Vypovziv, Chernihiv district, Chernihiv region) is a multi-layered archaeological site. It contains artefacts from the late Neolithic-Chalcolithic period (Kyiv-Pechersk culture and sites of the Pustynka-V type, which were part of the Dnipro-Donetsk ethno-cultural community 7000–4000 BCE), the Bronze Age (Middle Dnipro and Sosnytska cultures, 3000–1500 BCE), and the early Slavic period in the early Iron Age (1st–3rd c. CE).

The most intensive activity at the site took place from the 10th to the 13th c. CE, when it was inhabited by the population of Kievan Rus. Later, in 17th–18th c. CE, there was a wooden Christian church and a cemetery around it. The cemetery was excavated in 1889 and then in 2013–2014 by V. L. Berenshtam and V. M. Skorokhod. In total, 16 burials have been investigated to date, all of them were carried out according to the Christian rite, in graves and in wooden coffins. Bodies were put in an extended position on backs with heads oriented to the west. Burials also could be oriented parallel to the long axis of the church. Some burials included accompanying items such as coins and rings. ^262–264^

**Location:** 50.945951 N, 30.8039 E. Vypovziv village, Chernihiv district, Chernihiv region, Ukraine.

**Excavations:** 2013.

**Excavation authors:** Viacheslav M. Skorokhod, Institute of Archaeology of the National Academy of Sciences of Ukraine, Kyiv, Ukraine.

**Storage of anthropological materials:** Institute of Archaeology of the National Academy of Sciences of Ukraine, Kyiv, Ukraine.

**Description of samples:**

Two samples of the early modern period were taken for aDNA analyses.

**UKR164.** *Vypovziv, burial 12.* A woman aged 25–30 years old. The grave was approximately 0.5 m wide and studied to a length of 1.63 m. The grave was oriented along the east-west axis with a slight deviation to the south. Decayed wooden planks from a coffin were found within the grave. The skeletal remains of an adult individual were in extended position on the back, with the head facing the southwest. The hands were folded on the abdomen. The preservation of the bones was satisfactory. On the one hand, two rings made of coloured metal with square and round-shaped stone inserts were found. The bottom of the grave was at a depth of 0.2 m from the level of grave identification. The individual was identified as Ukrainian based on Orthodox Christian burial rite. Chronology according to archaeology: 18th c. CE.

**UKR165.** *Vypovziv, burial 13*. A woman aged 30–40 years old. The grave was approximately 0.4 m wide and studied to a length of 0.83 m. Decayed wooden planks from a coffin, about 0.35 m wide and up to 0.2 m in height, were identified. The upper part of skeletal remains of an adult individual were found, that lay in an extended position on the back, with the head facing the southwest. The preservation of bones was satisfactory. The bottom of the burial was at a depth of 0.25 m from the upper contour of the pit. The individual was identified as Ukrainian based on Orthodox Christian burial rite. Chronology according to archaeology: 18th c. CE.

### METHOD DETAILS

Sampling and DNA extraction were performed in dedicated ancient DNA laboratories of the Institute of Genomics, University of Tartu. Extract purification and library preparation were performed in dedicated ancient DNA laboratories of the Francis Crick Institute. DNA sequencing was performed at the Advanced Sequencing Facility, the Francis Crick Institute. The main steps of the laboratory work are detailed below. For 6 samples (UKR008, UKR030, UKR053, UKR070, UKR119, UKR147), sampling, DNA extraction, extract purification and library preparation were performed in dedicated ancient DNA laboratories of the Department of Archaeology and Anthropology, University of Cambridge as described in Järve et al. 2019 ^7^ and sequencing was performed at the Cambridge Biochemistry DNA Sequencing Facility and the Institute of Genomics Core Facility, University of Tartu.

#### DNA extraction

For 109 teeth, the apical tooth roots were cut off using a drill with a cutting wheel attached and used for extraction since root cementum has been shown to contain more endogenous DNA than crown dentine^265^. The root pieces were used whole to avoid heat damage during powdering with a drill and to reduce the risk of cross-contamination between samples. Contaminants were removed from the surface of tooth roots by soaking in 6% bleach for 5 minutes, then rinsing three times with milli-Q water (Millipore) and lastly soaking in 70% ethanol for 2 minutes, shaking the tubes during each round to dislodge particles. Then, the samples were left to dry under a UV light for 2 hours. Next, the samples were weighed, [20 * sample mass (mg)] µl of EDTA and [sample mass (mg) / 2] µl of proteinase K was added and the samples were left to digest for 72 hours on a rotating mixer at 20 °C to compensate for the smaller surface area of the whole root compared to powder.

For 14 bone fragments, the surface of the bone fragment was cleaned using a drill with a ball-end drill bit attached and powder from the inner part of the cortical bone was collected for extraction. Next, the samples were weighed, [20 * sample mass (mg)] µl or 1 ml (if >50 mg of powder) of EDTA was added and the samples were left to digest for 24 hours on a rotating mixer at 20 °C.

The lysate was concentrated to 250 µl (Vivaspin Turbo 15, 30,000 MWCO PES, Sartorius), 140 µl of concentrated lysate was transferred to a clean tube, frozen and shipped to the Francis Crick Institute on dry ice. Undigested material and leftover lysate was stored for a second DNA extraction if need be.

At the Francis Crick Institute, the concentrated lysate was transferred into FluidX tubes where it underwent automated purification on an Agilent Bravo Workstation ^266^.

#### Library preparation

Single-stranded DNA libraries were prepared from the extracts using an automated system ^267^. No treatment to remove uracils was used. The libraries were then double-indexed ^268^.

#### DNA sequencing

DNA was first sequenced using the Illumina HiSeq 4000 platform and around 2.5 million sequencing reads were generated per sample. Based on these data, 88 out of 123 samples were chosen to be included in further analyses. For most of the additional sequencing, fragments smaller than 35 bp were removed from the library ^267^. For this, 100 ng of the initial library was biotinylated and the non-biotinylated strand was isolated using streptavidin beads to obtain a single-stranded library. The samples were pooled and loaded onto a denaturing polyacrylamide gel along with 30 bp, 35 bp and 150bp insert markers and fragments with the desired length were excised and eluted from the gel after incubating overnight. Around 15.5 billion sequencing reads were generated using the Illumina NovaSeq platform with the 100 bp paired-end method.

The 6 samples processed in Cambridge were sequenced using the Illumina NextSeq 500 platform with the 75 bp single-end method and 4 were chosen to be included in further analyses.

### QUANTIFICATION AND STATISTICAL ANALYSIS

#### Mapping and quality filtering

Sequencing data was processed using the nf-core/eager v2 pipeline ^269^. First, adapters were removed, paired-end reads merged and bases with a quality below 20 trimmed using AdapterRemoval v2 ^270^ with options –trimns –trimqualities –collapse –minadapteroverlap 1 and –preserve5p. Next, reads with a minimum length of 35 bp were mapped to the hs37d5 human reference genome using Burrows-Wheeler Aligner (BWA-0.7.17 aln) ^271^ and parameters -l 16500 -n 0.01 -o 2 -t 1 ^268,272^. Then, the mapped BAM files were deduplicated using Dedup ^273^. Lastly, data from different sequencing runs was merged and duplicates were removed with picard 2.20.3 (https://broadinstitute.github.io/picard/) and reads with mapping quality under 10 were filtered out with samtools 1.9 ^274^.

The average endogenous DNA content (proportion of reads mapping to the human genome) for the 129 samples is 35% (Table S1). The endogenous DNA content is extremely variable as is common in aDNA studies, ranging from under 0.01% to almost 95% (Table S1). The average endogenous content for the 92 samples chosen for further analyses is 49%, varying between 1% and almost 95% (Table S1).

#### aDNA authentication

Ancient DNA damage in the form of C=>T substitutions in the 5’ ends of sequences due to cytosine deamination was estimated using DamageProfiler ^275^ from the nf-core/eager v2 pipeline ^269^. mtDNA contamination estimation was performed using schmutzi ^276^.

For the male individuals, contamination was also estimated based on chrX using the two contamination estimation methods first described in Rasmussen et al. 2011 ^277^ and incorporated in the ANGSD software ^278^ in the script contamination.R.

The samples show 23% C=>T substitutions at the 5’ ends on average, ranging from 0% to 61% in all samples and from 8% to 50% in the samples chosen for further analyses (Table S1). The mtDNA contamination point estimate for the samples chosen for further analyses ranges from 1% to 8% with an average of 2% (Table S1). The average of the two chrX contamination methods of male individuals chosen for further analyses with average chrX coverage >0.1x is 2% on average, ranging between 0% and 3% for most individuals (Table S1). For two individuals, the chrX contamination estimate is >10% (Table S1), but given that the mtDNA contamination estimate is low, average read length is similar to other samples and the C=>T substitution rate is high for both individuals, it was decided to include them in analyses, keeping the potential contamination in mind.

#### Kinship analysis

A total of 4,375,438 biallelic single nucleotide variant sites, with minor allele frequency (MAF) >0.1 in a set of over 2,000 high coverage genomes of Estonian Genome Center (EGC) ^279^ were identified and called with ANGSD ^278^ command --doHaploCall from the BAM files of all Iron Age samples (n=67), including 13 from the same groups published in Järve et al. 2019 ^7^, and all medieval (and early modern) samples (n=28) that were chosen for further analyses. Furthermore, the process was repeated for all 39 Scythian period samples and for smaller groups of samples with specific archaeological associations – 7 Thracian Hallstatt, 10 right bank of Dnipro Schytian, 10 left bank of Dnipro Scythian, 15 Siversky Donets basin Scythian, 4 Northern Black Sea region Scythian, 14 Chernyakhiv, 9 Saltiv, 7 Nogai. The Neolithic and Bronze Age samples were too few per group and spread over a too long time period to be included in kinship analyses. The ANGSD output files were converted to .tped format as an input for the analyses with READ script to infer pairs with up to 2^nd^ degree relatedness ^280^.

The results based on smaller groups and wider time periods are consistent with each other. Next to 1^st^ and 2^nd^ degree relative pairs, two identical pairs of samples are revealed (Figure S5). The first pair of identical samples – UKR035 and UKR038 (Figure S5A) – were thought to come from two skeletons in the same burial so we deduce that they most likely come from the same individual and the data were merged (UKR035AB) with samtools 1.9 ^274^ option merge. The second pair – UKR089 and UKR091 (Figure S5B) – come from the same site, but from two different kurgans excavated on different years so we consider that these could be identical twins. Hence, data from 91 newly sequenced individuals was used in further analyses. For group-based analyses, only the highest coverage individual of each group of relatives was used.

#### Calculating general statistics and determining genetic sex

Samtools 1.9 ^274^ option stats was used to determine the number of final reads, average read length, average coverage, etc for the 91 genomes.

Genetic sex was calculated using the script sexing.py from Skoglund et al. 2013 ^281^, estimating the fraction of reads mapping to Y chromosome out of all reads mapping to either X or Y chromosome. The average coverage of the whole genome for the 91 individuals is between 0.019x and 1.95x (Table S1). Of these, 13 genomes have an average coverage of less than 0.1x, 9 genomes >0.1x, 34 genomes >0.3x, 27 genomes >0.5x and 8 genomes >1x (Table S1). Genetic sexing reveals that the study involves 44 females and 47 males (Table 1, Table S1).

#### Determining mtDNA haplogroups

The program bcftools ^282^ was used to produce VCF files for mitochondrial positions – genotype likelihoods were calculated using the option mpileup and genotype calls were made using the option call. mtDNA haplogroups were determined by submitting the mtDNA VCF files to HaploGrep2 ^283,284^. Haplogroups were successfully determined for all 91 individuals (Table 1, Table S1).

#### Y chromosome variant calling and haplogroup determination

In total, 273,059 haplogroup informative Y chromosome variants from regions that uniquely map to Y chromosome ^285–289^ were called as haploid from the BAM files of the samples using the --doHaploCall function in ANGSD ^278^. Derived and ancestral allele and haplogroup annotations for each of the called variants were added using BEDTools 2.29.2 ^290^ intersect option. Haplogroup assignments of each individual sample were made manually by determining the haplogroup with the highest proportion of informative positions called in the derived state in the given sample. Y chromosome haplogrouping was blindly performed on all samples regardless of their sex assignment.

No female samples have reads on the Y chromosome consistent with a haplogroup, indicating that levels of male contamination are negligible. Haplogroups for 45 (with coverage >0.01x) out of the 47 males were successfully determined (Table 1, Tables S1–S2).

#### Genome-wide variant calling

Genome-wide variants were called with the ANGSD software ^278^ command --doHaploCall, sampling a random base for the positions that are present in the Allen Ancient DNA Resource (AADR) ^43^ version 54.1.

#### Preparing the datasets for autosomal analyses

Individuals from the Human Origins (HO) array dataset from AADR ^43^ version 54.1 was used as the modern DNA background. Individuals from the 1240K dataset from AADR ^43^ version 54.1 were used as the ancient DNA background.

The data of the comparison datasets and of the individuals of this study was converted to BED format using PLINK 1.90 (https://www.cog-genomics.org/plink/) ^291^ and the datasets were merged. Two datasets were prepared for analyses: one with HO and 1240K individuals and the individuals of this study, where 577,902 autosomal SNPs of the HO dataset were kept; the other with 1240K individuals and the individuals of this study, where 1,127,956 autosomal and 45,218 chrX SNPs of the 1240K dataset were kept. Individuals with <10,000 SNPs overlapping with the HO autosomal dataset were removed from further autosomal analyses.

#### Principal component analysis

To prepare for principal component analysis (PCA), two reduced comparison sample-sets were assembled, both including 3,298 ancient individuals from 400 populations, while one composed of 813 modern individuals from 54 populations of Europe, Caucasus and Near East (West Eurasia) and the other included in addition 1,230 modern individuals from 86 populations (Eurasia) (Tables S3–S4). Only SNPs with MAF >0.05 in the HO dataset were used (n=409,113). The data was converted to EIGENSTRAT format using the program convertf from the EIGENSOFT 8.0.0 package ^66^. PCA was performed with the program smartpca from the same package, projecting ancient individuals onto the components constructed based on the modern genotypes using the option lsqproject.

#### Admixture analysis

For Admixture analysis ^292^, the same ancient sample-set was used as for PCA and the modern sample-set included 1861 individuals from 144 populations from all over the world (Tables S3–S4). Only SNPs with MAF >0.05 in the HO dataset were used and the HO dataset of modern individuals was further pruned to decrease linkage disequilibrium using the option indep-pairwise with parameters 1000 250 0.4 in PLINK 1.90 (https://www.cog-genomics.org/plink/) ^291^. This resulted in a set of 161,774 SNPs.

Admixture was run using ADMIXTURE 1.3 ^292^ on the modern dataset using K=3 to K=14 in 100 replicates. This enabled us to assess convergence of the different models. K=10 was the model with the largest number of inferred genetic clusters for which >10% of the runs that reached the highest Log Likelihood values yielded very similar results. This was used as a proxy to assume that the global Likelihood maximum for this particular model was indeed reached. Then the inferred genetic cluster proportions and allele frequencies of the best run at K=10 were used to run Admixture using ADMIXTURE 1.3 ^292^ with the P option to project the aDNA individuals. The same projecting approach was taken for all models (K3–14). We present all ancient individuals on Figure S4AD but only population averages on Figure 4.

Admixture was also run using only ancient individuals and without projection. For this, the same ancient sample-set was used, but with the SNPs of the 1240K dataset. The dataset was pruned to decrease linkage disequilibrium using the option indep-pairwise with parameters 50 10 0.1, resulting in a set of 366,718 SNPs. Samples with more than 90% of missing data were removed with the mind option in PLINK 1.90 (https://www.cog-genomics.org/plink/) ^291^. Next, the comparative dataset was revised to include a maximum of 5 individuals per group, resulting in 1,436 individuals to be used in the analysis, including 79 individuals of this study. Admixture was run using ADMIXTURE 1.3 ^292^ using K=2 to K=6 in 100 replicates. This enabled us to assess convergence of the different models. K=4 was the model with the largest number of inferred genetic clusters for which >10% of the runs that reached the highest Log Likelihood values yielded very similar results. The inferred genetic cluster proportions and allele frequencies were also used to run Admixture using ADMIXTURE 1.3 ^292^ with the P option to project all aDNA individuals for comparison. We present all ancient individuals on Figure S4BCEF but only population averages on Figure S3 – based on non-projected results for groups that could be included in that analysis, based on projected results for other groups.

#### f4 statistics

For calculating f4 statistics, the SNPs of the 1240K dataset were used with Mbuti as outgroup and the same ancient sample-set as for previous analyses, but only including comparative groups with at least 3 individuals, resulting in 3,138 individuals from 314 populations (Tables S3–S4). The data was converted to EIGENSTRAT format using the program convertf from the EIGENSOFT 8.0.0 package^66^. f4 statistics of the form f4(Mbuti, other ancient; Ukraine ancient, Ukraine/other relevant ancient) were calculated using the ADMIXTOOLS 2 ^293,294^ package program qpDstat.

#### Outgroup f3 statistics

Autosomal outgroup f3 statistics were calculated on the same dataset as f4 statistics (Tables S3–S4). To allow for chrX *versus* autosomes comparison for ancient populations, outgroup f3 statistics were run both using 1,127,956 autosomal SNPs and also 45,218 chrX positions available in the 1240K dataset. Since all children inherit half of their autosomal material from their father but only female children inherit their chrX from their father then in this comparison chrX data gives more information about the female and autosomal data about the male ancestors of a population. Outgroup f3 statistics of the form f3(Mbuti; Ukraine ancient, other ancient) were computed using the ADMIXTOOLS 2 ^293,294^ package program qp3Pop. Results for combinations including UkrN_?, UkrFBA/EIA_Vysotska_Early, UkrEIA_Scythian_SivDon_NomEl_2, SaltovoMayaki and China_Wuzhuangguoliang_LN were not included in chrX *vs* autosomes comparisons since <1,000 SNPs were used in many of these combinations based on chrX data.

#### qpAdm

The ADMIXTOOLS 2 ^293,294^ package program qpAdm was used to estimate which populations and in which proportions are suiTable proxies of admixture to form the groups of this study. The autosomal positions of the 1240K dataset were used. Only samples with more than 100,000 SNPs were used in the analyses. Mota, Ust-Ishim, Kostenki14, GoyetQ116, Vestonice16, MA1, AfontovaGora3, ElMiron, Villabruna, Afanasievo, Anatolia_N, Mongolia_North_N were used as “right” populations. For distal models, Yamnaya_Samara, Yamnaya_Kalmykia or Ukraine_Yamnaya was used as the “left” population representing Steppe ancestry, LBK_EN, Central_MN, Poland_GlobularAmphora, Ukraine_GlobularAmphora or Ukraine_Trypillia was used as the “left” population representing early farmer ancestry, and Mongolia_LBA_Khovsgol_6, Mongolia_LBA_CenterWest_4 or Mongolia_EIA_SlabGrave_1 was used as the “left” population representing East Asian ancestry. For proximal models, groups of this study from earlier time periods were used as the “left” populations representing local ancestry, and Mongolia_EIA_SlabGrave_1 was used as the “left” population representing East Asian ancestry.

#### Comparing genetic heterogeneity

Analysis of genetic heterogeneity made use of eigenvectors obtained from the preceding PCA analysis to compare Euclidean distances between genomes in multidimensional principal components space. Up to 100 PCs were generated using the smartpca method ^66^ from the EIGENSOFT package (detailed above). Ukraine genomes generated in this study were subset into the following date ranges based on the midpoints of the 95% calibrated date estimates or archaeological date range estimates: Late Bronze Age and pre-Scythian Early Iron Age (LBAEIA, 3,000–700 BCE, n=15); Scythian period Early Iron Age (SEIA, 699–300 BCE, n=27); post-Scythian Iron Age to Early Middle Ages (IAEMA, 299 BCE– 900 CE, n=26); and Middle Ages and early modern period (MAEM, 901–1,800 CE, n=19). One Neolithic (UKR008), two Scythian-associated (UKR095 and UKR096) and one potentially Greek-associated genome (UKR153) were excluded from the analysis due to falling outside of the specified date range. To contextualize these PCA-based heterogeneity estimates, comparative datasets were generated from 3,298 previously published ancient genomes from Western Eurasia. We subset these genomes into the same four date ranges as for the Ukraine data. From the geographic coordinates of the individuals of this study, the radius of the minimal bounding circle was obtained for each date range, using the haversine formula implemented in R to calculate great-circle distances. From each of the comparative data date ranges described above, sets of subsamples were generated by randomly selecting up to 100 individuals, and then for each of these individuals, retaining all other reference genomes within the same radius distance as the corresponding Ukraine data. This resulted in up to 100 randomly positioned subsets of equivalently proximate individuals, setting a minimum of 10 individuals per subset. As this was limited by the number of unique individuals with >10 others within the radius distance, the resulting number of subsets was in some cases <100 (LBAEIA n=100, SEIA n=63, IAEMA n=100, MAEM n=28). For each date range, pairwise Euclidean distances in PC-space were then determined between the genomes of this study and also between the genomes in each comparative subset using the first 25 principal components, scaling each eigenvector by its eigenvalue. Boxplots and kernel density estimates were then generated in R to visually compare the resulting sets of distances. All these methods are documented in a reproducible Rscript available on Github.

## Notes

### Competing Interest Statement

The authors have declared no competing interest.

