## Supplemental Information for "North Pontic crossroads: Mobility in Ukraine from the Bronze Age to the early modern period"

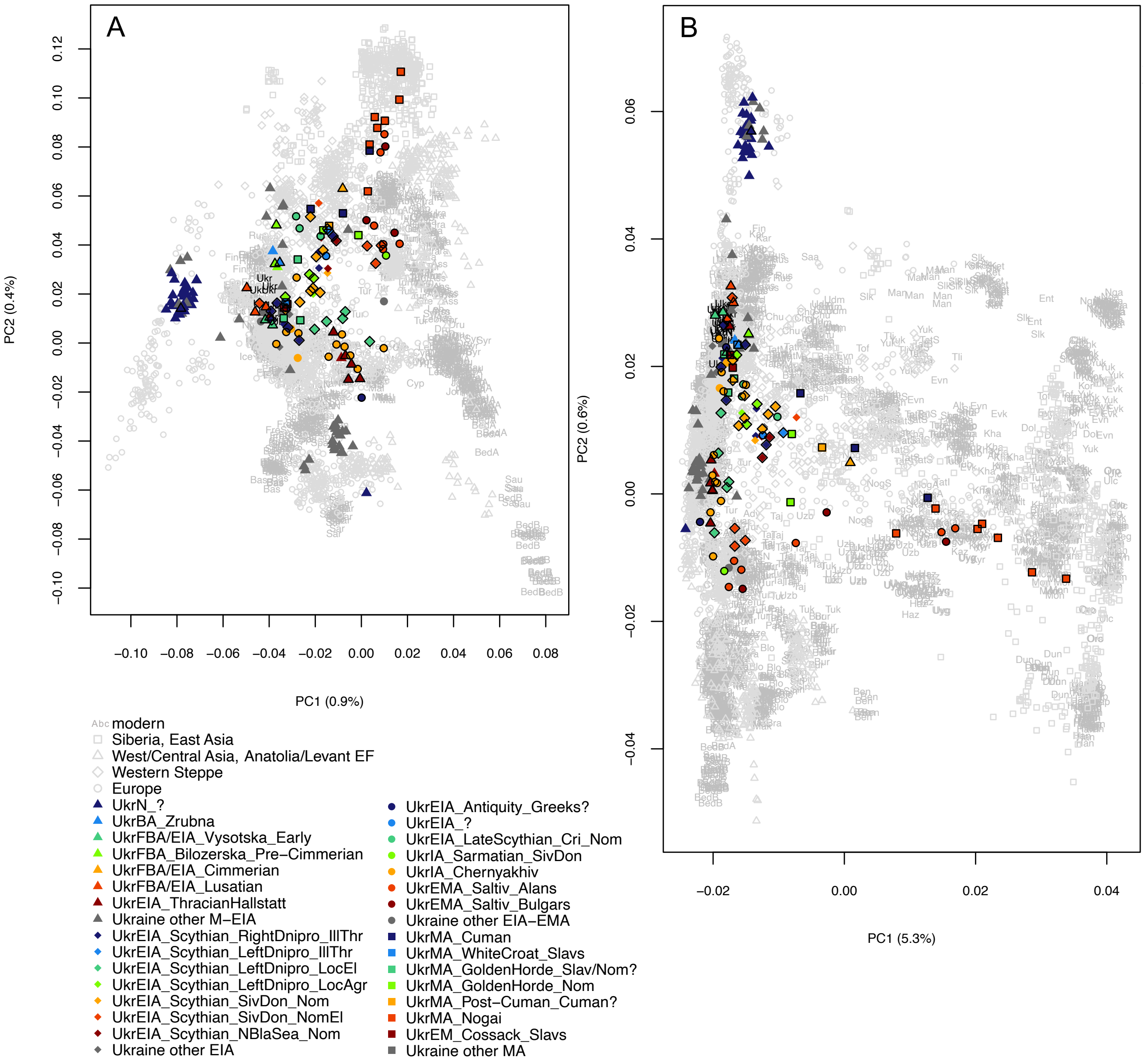


**Figure S1. Full principal component analysis results.** Principal component analysis results of modern (**A**) West Eurasians, (**B**) Eurasians with ancient individuals projected onto the first two components (PC1 and PC2). Newly reported individuals are indicated with a black outline. Modern Ukrainians are shown in black.

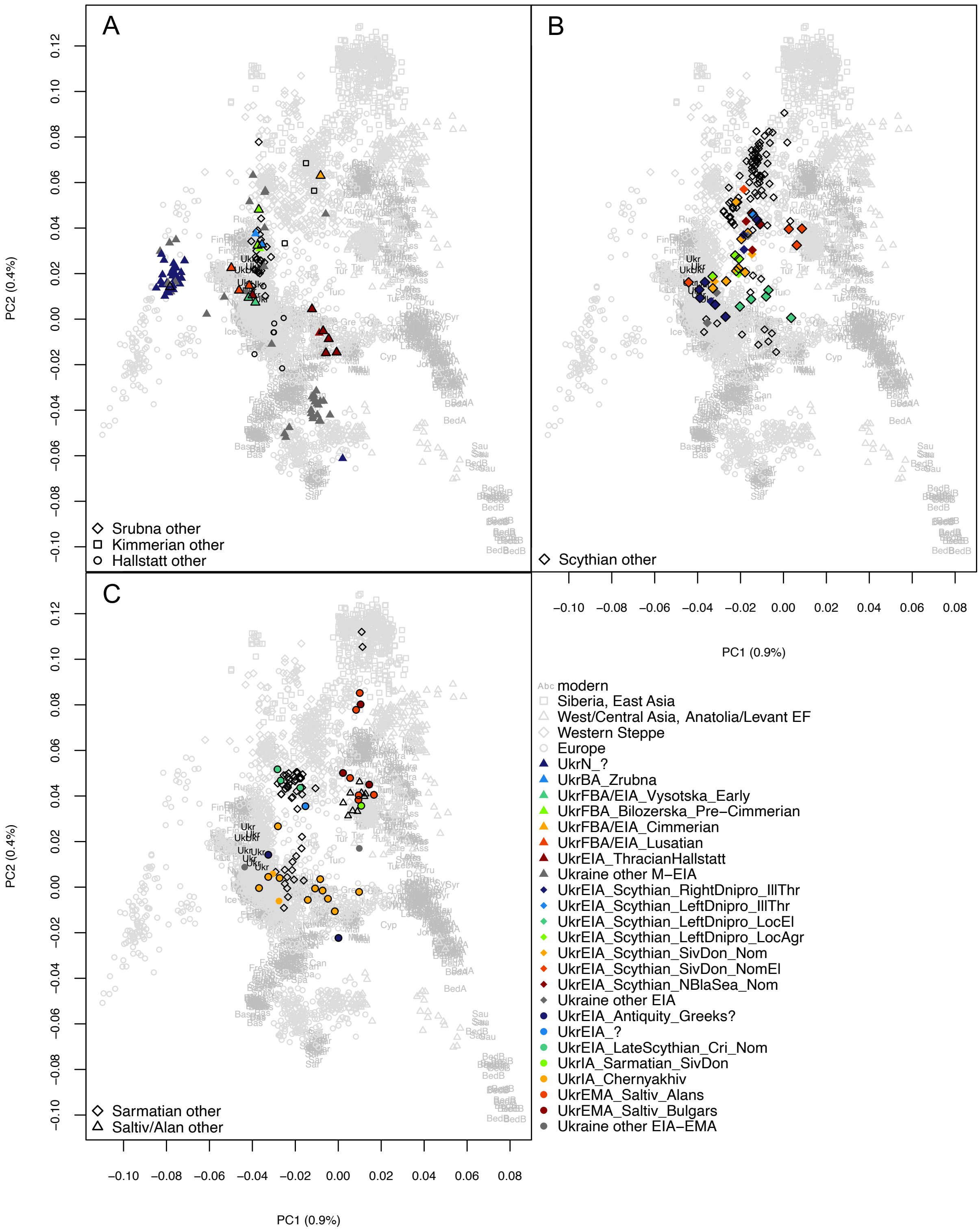


**Figure S2. Principal component analysis results with relevant previously published individuals outlined.** Principal component analysis results of modern West Eurasians with ancient individuals projected onto the first two components (PC1 and PC2). Ukrainian and previously published groups from (**A**) Late Bronze Age and pre-Scythian Iron Age (3,000–700 BCE), (**B**) the Scythian period of Early Iron Age (700–300 BCE), (**C**) post-Scythian Iron Age until Early Middle Ages (400 BCE–900 CE). Newly reported individuals are indicated with a black outline. Modern Ukrainians are shown in black.

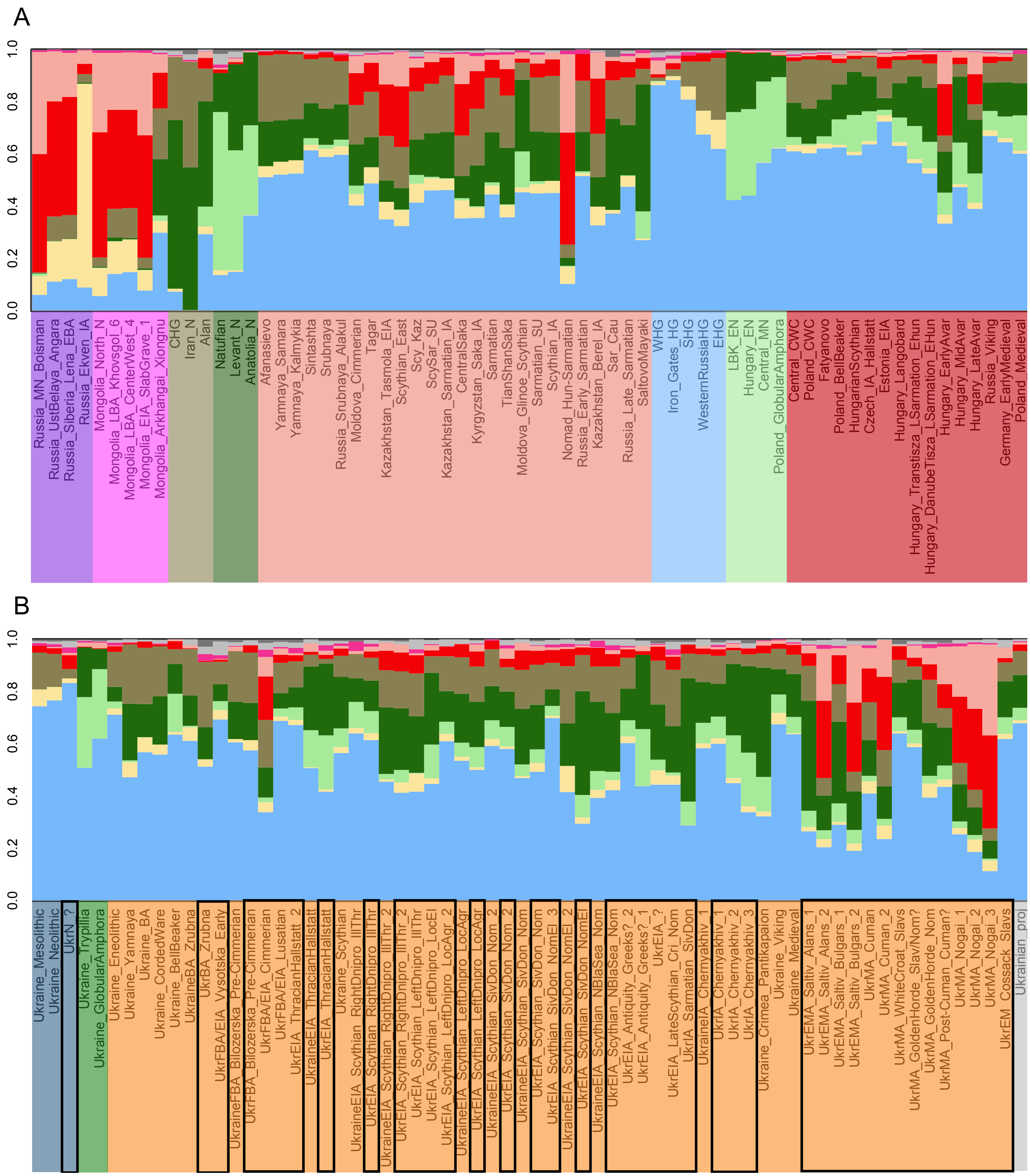


**Figure S3. Subset of ADMIXTURE analysis results with ancient individuals projected onto modern structure.** Population averages of (**A**) Eurasia-wide, (**B**) Ukrainian ancient groups at K10.

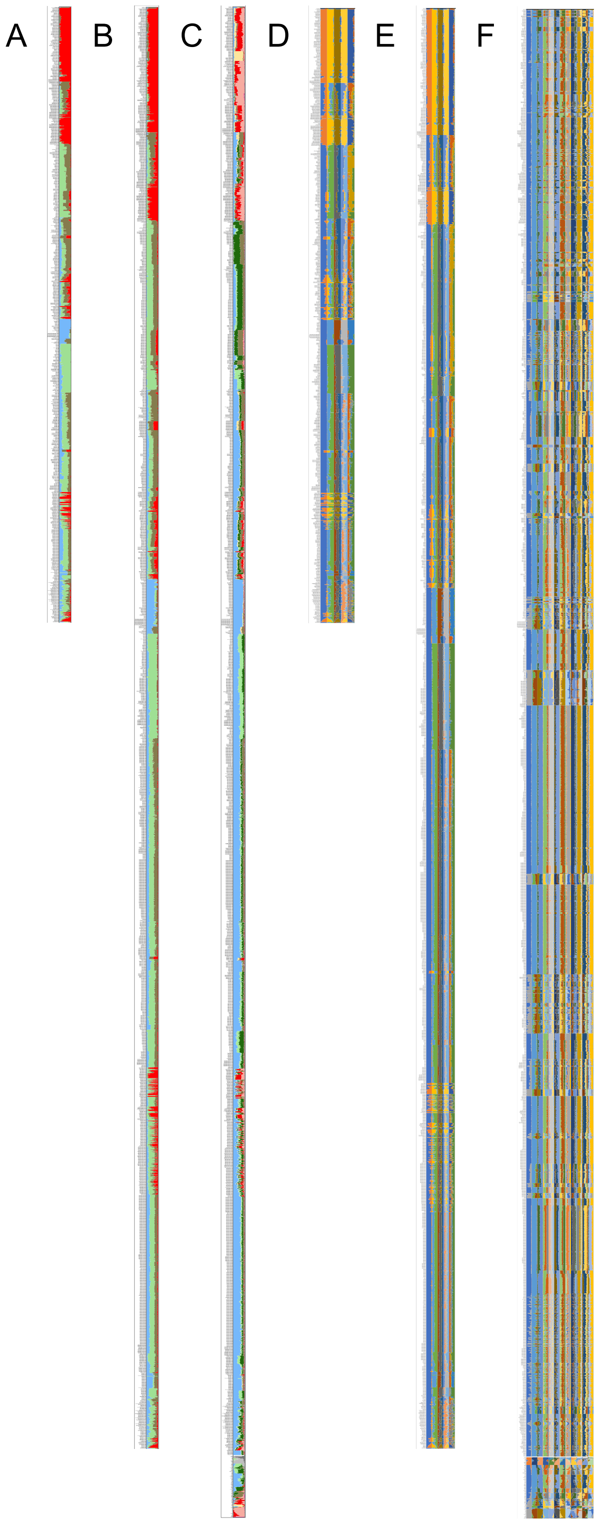


**Figure S4. Full ADMIXTURE analysis results.** (**A**) ancient individuals at K4, (**B**) ancient individuals projected onto ancient structure at K4, (**C**) projected ancient individuals and modern population averages at K10, (**D**) ancient individuals at K2 to K6, (**E**) ancient individuals projected onto ancient structure at K2 to K6, (**F**) projected ancient individuals and modern population averages at K3 to K14.

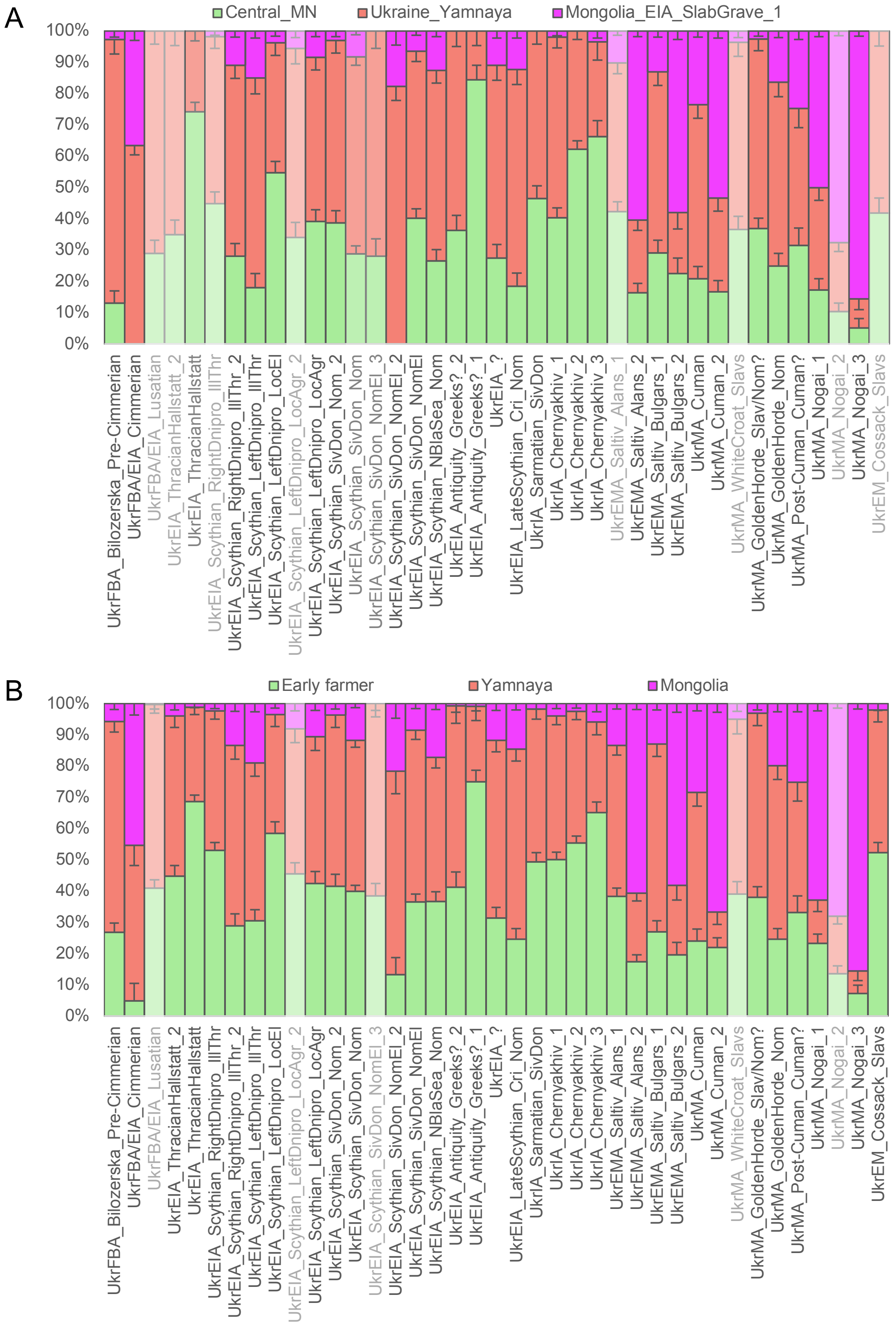


**Figure S5. Additional qpAdm admixture modelling results.** Distal qpAdm models of admixture between (**A**) Central_MN, Ukraine_Yamnaya and Mongolia_EIA_SlabGrave_1, (**B**) a European early farmer group, a Yamnaya group and a Mongolian group resulting in the highest p value for each target group, tested using the autosomal positions of the 1240K dataset. Models with non-significant p-values ($p<0.05$) are semi-transparent.

A

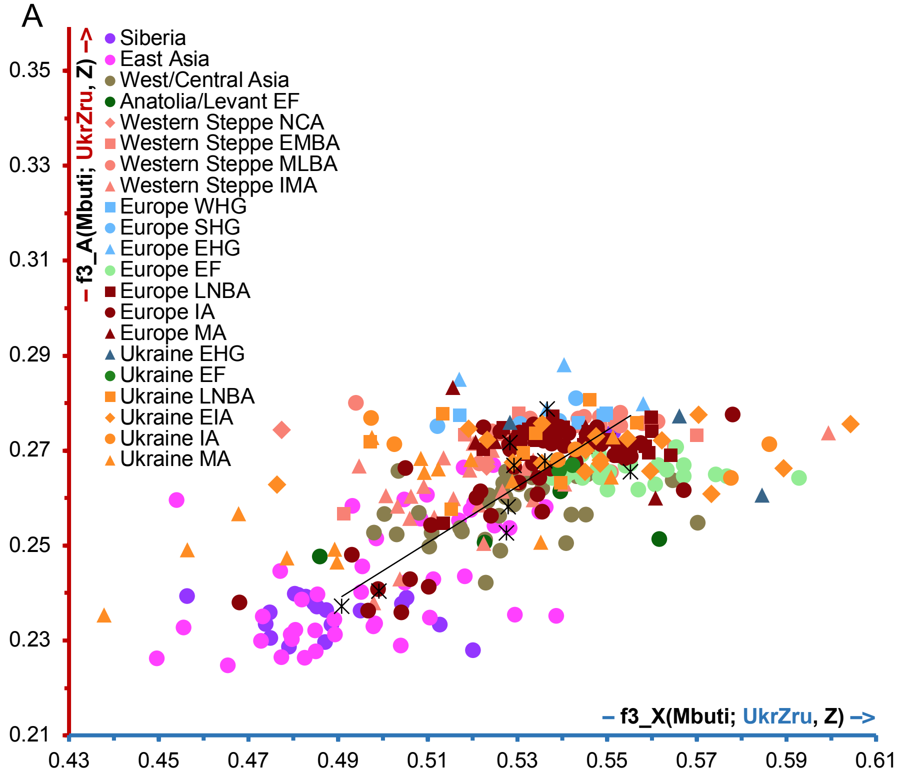


B

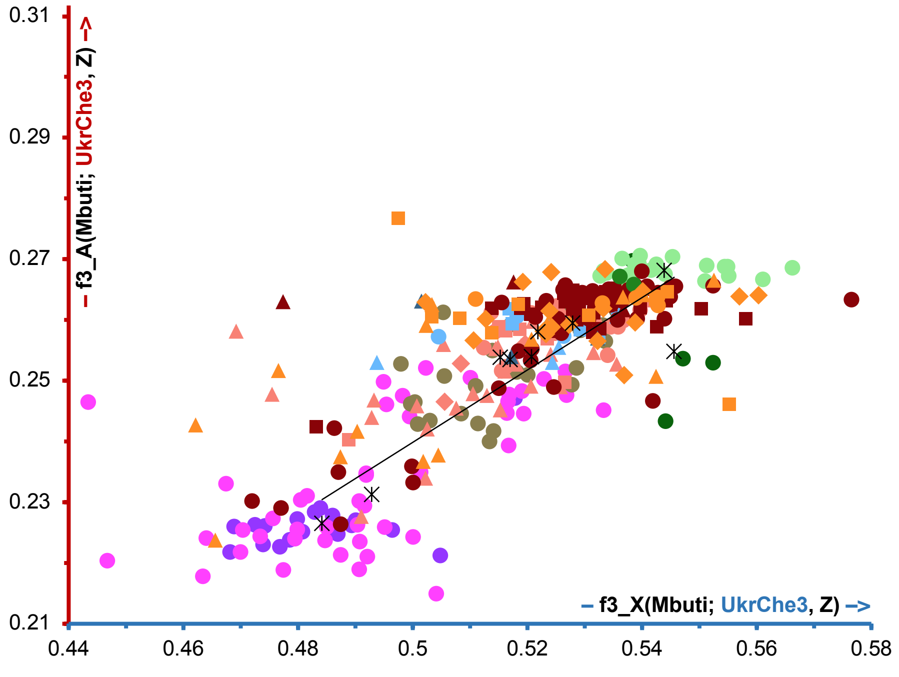


**Figure S6. Outgroup f3 statistics' results of comparisons with ancient populations.** Outgroup f3 statistics' values of form f3(Mbuti; Ukrainian population, ancient) using chr X/autosomal SNPs of the 1240K dataset. Ukrainian population: (**A**) Zrubna, (**B**) Chernyakhiv_3. EF – early farmers; NCA – Neolithic/Copper Age; EMBA – Early/Middle Bronze Age; MLBA – Middle/Late Bronze Age; IMA – Iron/Middle Ages; HG – hunter-gatherers, W – Western, S – Scandinavian, E – Eastern; LNBA – Late Neolithic/Bronze Age; IA – Iron Age; MA – Middle Ages. The average values for groups shown in the legend are shown with black asterisks. The trendline for the black asterisks is shown in black.

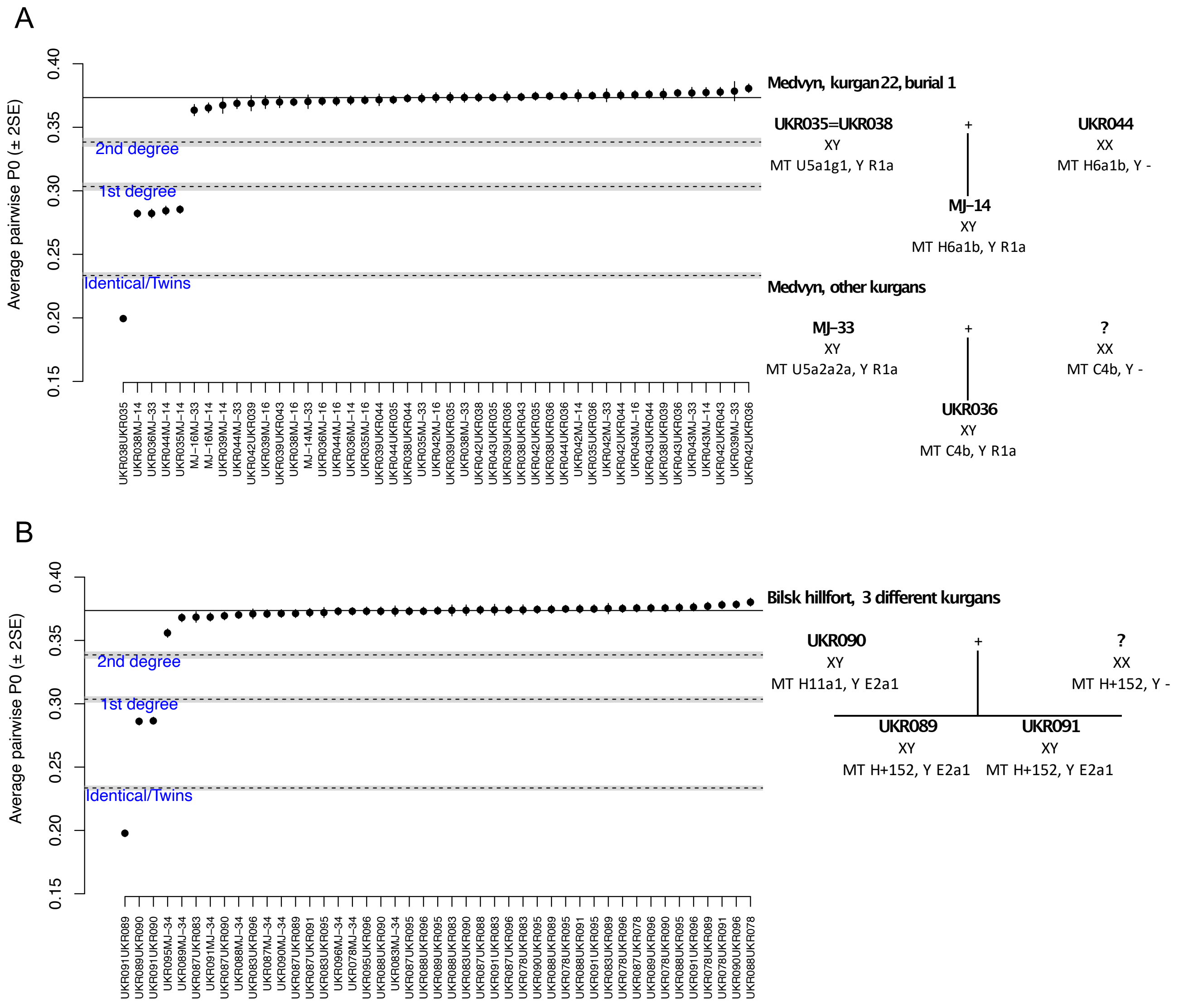


**Figure S7. Full kinship test results.** (**A**) right bank of Dnipro Scythians' kinship test results, (**B**) left bank of Dnipro Scythians' kinship test results.
